# Higher-order epistasis and phenotypic prediction

**DOI:** 10.1101/2020.10.14.339804

**Authors:** Juannan Zhou, Mandy S. Wong, Wei-Chia Chen, Adrian R. Krainer, Justin B. Kinney, David M. McCandlish

## Abstract

Contemporary high-throughput mutagenesis experiments are providing an increasingly detailed view of the complex patterns of genetic interaction that occur between multiple mutations within a single protein or regulatory element. By simultaneously measuring the effects of thousands of combinations of mutations, these experiments have revealed that the genotype-phenotype relationship typically reflects genetic interactions not only between pairs of sites, but also higher-order interactions among larger numbers of sites. However, modeling and understanding these higher-order interactions remains challenging. Here, we present a method for reconstructing sequence-to-function mappings from partially observed data that can accommodate all orders of genetic interaction. The main idea is to make predictions for unobserved genotypes that match the type and extent of epistasis found in the observed data. This information on the type and extent of epistasis can be extracted by considering how phenotypic correlations change as a function of mutational distance, which is equivalent to estimating the fraction of phenotypic variance due to each order of genetic interaction (additive, pairwise, three-way, etc.). Using these estimated variance components, we then define an empirical Bayes prior that in expectation matches the observed pattern of epistasis, and reconstruct the genotype-phenotype mapping by conducting Gaussian process regression under this prior. To demonstrate the power of this approach, we present an application to the antibody-binding domain GB1 and also provide a detailed exploration of a dataset consisting of high-throughput measurements for the splicing efficiency of human pre-mRNA **5**′ splice sites, for which we also validate our model predictions via additional low-throughput experiments.

## Introduction

Understanding the relationship between genotype and phenotype is difficult because the effects of a mutation often depend on which other mutations are already present in the sequence, a phenomenon known as epistasis [1–3]. Recent advances in high-throughput mutagenesis and phenotyping have for the first time provided a detailed view of these complex genetic interactions, by allowing phenotypic measurements for the effects of tens of thousands of combinations of mutations within individual proteins [4–18], RNAs [19–24], and regulatory or splicing elements [25–31]. Importantly, it has now become clear that the data from these experiments cannot be captured by considering simple pairwise interactions, but rather higher-order genetic interactions between three, four, or even all sites within a functional element are empirically common [2, 12, 32–44] and indeed often expected based on first-principles biophysical considerations [12, 23, 32, 35, 36, 41, 45, 46]. However, the enormous number of possible combinations of mutations makes these higher-order interactions both difficult to conceptualize and challenging to incorporate into predictive models.

From a very basic perspective, data from combinatorial mutagenesis experiments provide us with observations of phenotypic values for individual genotypes, the effects of specific mutations on specific genetic backgrounds, epistatic coefficients between pairs of mutations on specific backgrounds, etc. The essential problem in modeling data like this then comes down to the question of how to combine these observed quantities to make phenotypic predictions for unobserved genotypes. That is, given that we have already seen the results of a specific mutation in several different genetic backgrounds, how should we combine these observations to predict its effect in a new background?

Here, we provide an answer to this question based on the intuition that, when making these predictions, we should focus on the observed effects of mutations that are nearby in sequence space to the genetic background we are making a prediction for, rather than observations of mutational effects that are more distant. We do this by considering a key comprehensible aspect of higher-order epistasis, namely the decay in the predictability of mutational effects, epistatic coefficients of double mutants, and observed phenotypes, as one moves through sequence space. We show analytically that the shape of how this predictability decays as a function of distance is completely determined by the fraction of phenotypic variance due to each order of genetic interaction (additive, pair-wise, three-way, etc.). Thus, rather than conceptualizing higher-order epistasis in terms of innumerable interaction terms between larger and larger numbers of sites, we suggest that: (1) we can understand a great deal about higher-order epistasis by considering simple diagrams showing how the correlations between mutational effects, epistatic coefficients, etc. decay as a function of genetic distance; and (2) these same diagrams suggest a method for making phenotypic predictions by weighting our observations in terms of the degree of information they provide for mutations on a genetic background of interest.

We implement these ideas in terms of a Gaussian process regression [47] framework with an empirical Bayes [48] prior. Specifically, we use the observed pattern of decay in phenotypic correlation as a function of genetic distance to estimate the fraction of variance due to each order of interaction in our observed data. We then use these point estimates of the variance components to construct a prior distribution over all possible sequence-to-function mappings, where the expected decay in the predictability of mutational effects in the prior matches that observed in the data. Finally, we conduct Bayesian inference under this prior, using Hamiltonian Monte Carlo [49] to sample from the resulting high-dimensional posterior distribution. The end result is a procedure that automatically weights the contributions of our observations to our predictions in the manner suggested by the overall form of higher-order epistasis present in the data, while simultaneously accounting for the effects of measurement noise and quantifying the uncertainty in our predictions.

To demonstrate the performance of this technique, we apply our method to two datasets. The first dataset is derived from a combinatorial mutagenesis experiment for protein G [37], a streptococcal antibody-binding protein that has served as a model system for studies of the genotype-phenotype map in proteins. The second dataset consists of high-throughput measurements of the splicing efficiency of human 5′ splice sites [31], which are RNA sequence elements crucial for the assembly of the spliceosome during pre-mRNA splicing. For this latter dataset, we also present low-throughput validation measurements for our model predictions, as well as a qualitative exploration of the complex patterns of epistasis in splicing efficiency observed in this system.

## Results

Experimental observations of the genotype-phenotype map typically consist of measurements of phenotypic values for a subset of possible genotypes. From these observations, we can calculate a number of more derived quantities, such as mutational effects (i.e., the difference in phenotype between two mutationally adjacent genotypes) and double-mutant epistatic coefficients (i.e., the difference between the observed phenotype of a double mutant and its expected phenotype based on the sum of the single-mutant effects). The central question of phenotypic prediction is then deciding how to combine these various mutational effects, local epistatic coefficients and individual phenotypic values to produce accurate predictions for the phenotypic values of unmeasured genotypes.

Different prediction methods typically reflect different overall strategies for how to combine these experimentally derived quantities. For example, when we fit an additive or non-epistatic model [50], we are implementing a strategy based on the assumption that the phenotypic effects of observed mutations are the same regardless of the presence/absence of other mutations. Thus, fitting an additive model can be thought of as a generalization of the simple heuristic procedure of making predictions by: (1) averaging over all the times the effect of each possible point mutation is observed; and then (2) adding up these average effects to make a prediction for any given genotype. In a similar way, it is easy to show that while a pairwise interaction model [51] allows the effects of individual mutations to vary across genetic backgrounds, the epistatic interaction observed in double mutants for any specific pair of mutations is constant across backgrounds (see *SI Appendix*). Thus, fitting a pairwise model is conceptually closely related to the heuristic of determining the interaction between a pair of mutations by averaging over the local epistatic coefficients for this pair of mutations that are observed in the data and then assuming that this pair of mutations has the same interaction regardless of what genetic background these mutations occur on.

Putting the underlying strategies of additive and pairwise interaction models in these terms helps clarify the deficiencies of these models. Both models assume that only interactions between a certain number of mutations are relevant to prediction (i.e., additive effects of single mutations in non-epistatic models and interactions between two sites in pairwise interaction models). And both models assume that these interactions or mutational effects do not vary over sequence space, first by pooling information across all observed sequences to estimate these interactions or mutational effects, and then by making predictions that extrapolate these observations to all possible genetic backgrounds—even to areas of sequence space where we have little or no data.

Here we introduce a prediction method corresponding to a different heuristic, one that implements the intuitions that: (1) all orders of genetic interaction can be important and helpful in making predictions; and (2) observations of mutational effects and epistatic coefficients in nearby genetic backgrounds should influence our predictions more than observations in distant genetic backgrounds.

### Genetic interactions and the predictability of mutational effects

To implement a strategy of this type, it will be helpful to present some general results concerning higher-order epistasis. We begin by considering the case where all phenotypic values are known, before proceeding to our main problem of predicting unknown phenotypic values.

Our first task is to understand the relationship between the spatial scale of smoothness or predictability in the genotype-phenotype map, the amount of higher order epistasis, and the typical magnitude of epistatic interactions of various orders. These features of the genotype-phenotype map are illustrated for a simulated complete genotype-phenotype mapping in Figure 1A-C. Figure 1A shows the distance correlation function (ref. [33, 52, 53] and *SI Appendix*), which plots how correlations between phenotypic values drop off as one moves through sequence space. Figure 1B shows the decomposition of the genotype-phenotype map into variance components, where the fraction of variance due to a particular interaction order is defined to be increase in the *R*^2^ of a least squares fit when one adds interaction terms of that order to a model that already includes all interactions of lower order (*SI Appendix*). This set of variance components is also known in the literature as the normalized amplitude spectrum [33, 53]. Figure 1C shows how large the individual regression coefficients of a given order (ref. [33, 53, 54] and *SI Appendix*) tend to be, by plotting the mean square regression coefficient size as a function of interaction order (more precisely, the regression coefficients for order *k* are expressed with respect to an arbitrary orthonormal basis that spans the orthogonal complement of the space of all (*k* - 1)th order interaction models within the space of *k*-th order models).

**Figure 1:**
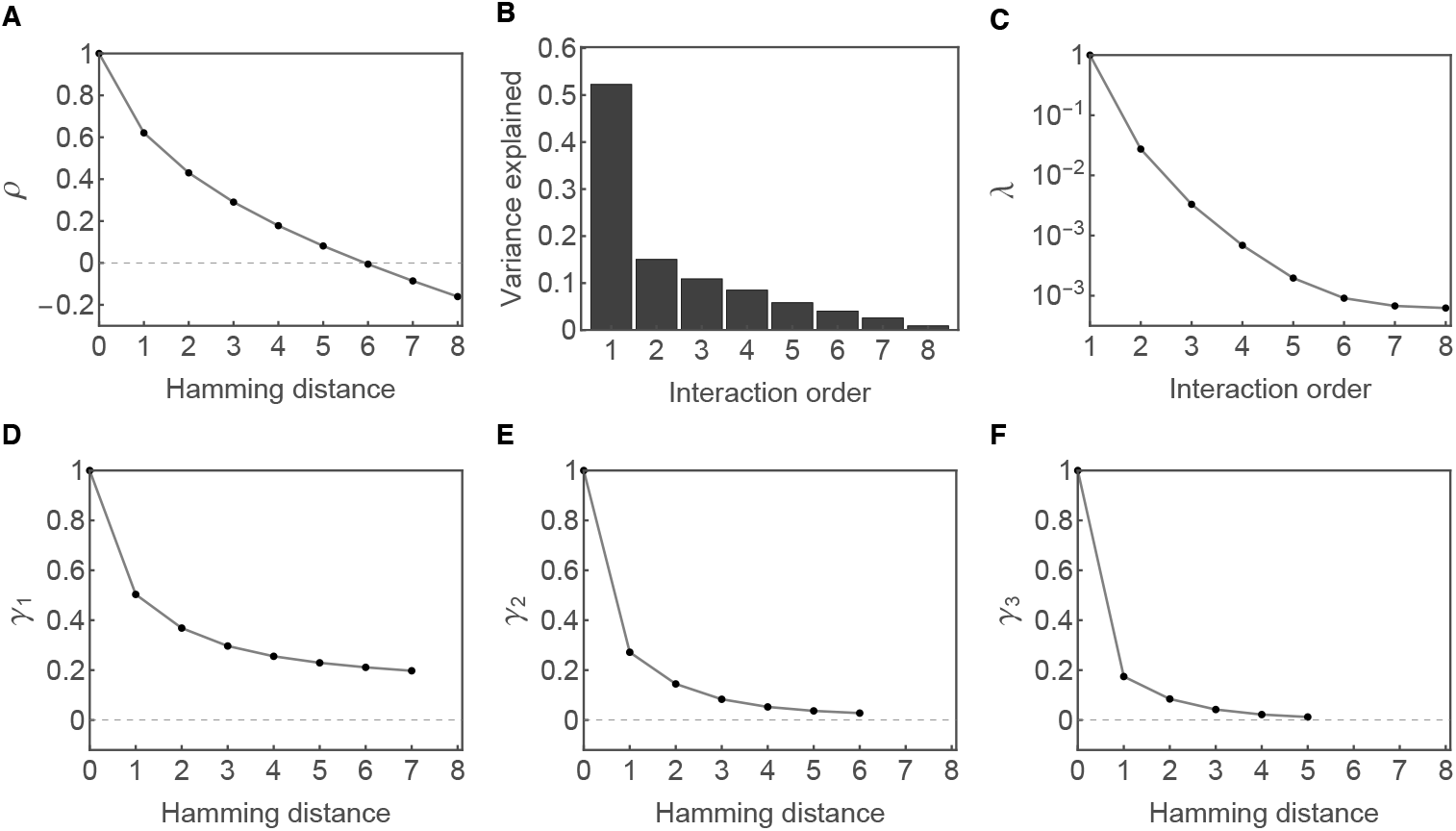
Summary statistics of a simulated genotype-phenotype map on sequences of length 8 with 4 alleles per site. (A) Distance correlation function for phenotypes (*ρ*). This is equal to the correlation between phenotypic values averaged over all pairs of sequences separated by the specified number of mutations (Hamming distance). (B) Decomposition of the genotype-phenotype map into variance components. Specifically, the *k*-th order variance component is equal to the increase in *R*^2^ of a least squares fit when when one adds *k*-th order interaction terms to a model only containing interactions up to order *k* − 1. (C) Mean squared regression coefficients of different orders (*λ*). (D) Distance correlation function for mutational effects (*γ*_1_), (E) distance correlation function for local pairwise epistatic coefficients (*γ*_2_), and (F) distance correlation function for local three-way interactions (*γ*_3_). Formulas for calculating these statistics can be found in *SI Appendix*.

Because our goal is to understand how to combine the mutational effects, observed double-mutant epistatic coefficients, etc., we can also plot how the predictability of these effects drops off as we move through sequence space [12, 55]. The distance correlation functions for mutational effects, local pairwise epistatic coefficients, and local three-way interactions are shown in Figure 1D-F, respectively.

These pictures, particularly the plots of correlations as a function of distance in genotypic space, are quite informative for our intuitive goal of determining how to combine our observations of mutational effects, local epistatic coefficients, etc. when making predictions. We see for example from Figure 1D that, for this particular genotype-phenotype map, mutational effects remain moderately correlated across all of sequence space, dropping from having a Pearson correlation coefficient of roughly 0.5 in adjacent genetic backgrounds to a correlation coefficient of roughly 0.2 in maximally distant backgrounds. However, from Figure 1E we see that the predictability of interactions in double-mutants decays much more rapidly, and so our observations of double-mutant interactions are only really informative in genetic backgrounds up to approximately two mutations away. Similarly, Figure 1F shows that observed triple-mutant interactions are only substantially informative in immediately adjacent genetic backgrounds. These results suggest that when making predictions, it might, e.g., be sensible to extrapolate our observations of mutational effects throughout sequence space, but only allow our observations of interactions in local double and triple mutant cycles to influence our predictions in relatively nearby genetic backgrounds.

How can we convert these intuitions based on examining the decay in the consistency of observed interactions into a rigorous method of phenotypic prediction? The key in answering this question lies in the fact that all 6 panels of Figure 1 are actually intimately related with each other and with previously proposed methods for phenotypic prediction. In particular, it is classically known that the three pictures in Figure 1A-C actually contain identical information, in the sense that for any given genotype-phenotype map, having any one of the panels in the top row of Figure 1 allows us to compute the other two (ref. [33, 53, 56], *SI Appendix*). Moreover, recent results on the distance correlation function of mutational effects (previously denoted by *γ*, [12, 55]) have shown that Figure 1D can be derived from any of Figure 1A-C. Here, we extend these results to arbitrary orders of interaction, so that Figure 1E-F and their higher-order analogs can likewise be derived from any of Figure 1A-C (*SI Appendix*). In total, these results show that all the information on how mutational effects and local epistatic interactions (of arbitrary order) generalize across genetic backgrounds is in fact captured by any one of the pictures in Figure 1A-C and can hence be specified using only *ℓ* − 1 parameters, where *ℓ* is the sequence length.

### Higher-order epistasis and the geometry of genotype-phenotype maps

To see how exactly Figures 1A-F are related, we need to develop a deeper understanding of the relationship between higher-order epistasis and the decay of phenotypic predictability across increasingly divergent genetic backgrounds. If we again consider constructing a least-squares fit of a genotype-phenotype map using up to *k*-th order interactions and compare it to the least squares fit using only up to (*k* − 1)th order interactions, the difference between these two fits represents the pure *k*-th order interaction component of the genotype-phenotype map, in the sense that it is the component that can be expressed using *k*-th order interactions but cannot be constructed using only lower-order interactions. The crucial observation is that these pure *k*-th order components each have a very specific geometry, and as a result the distance correlation function for any such *k*-th order component must take a very specific shape. Technically, these shapes are given by a set of orthogonal polynomials known as the Krawtchouk polynomials [33, 56–58], but for our purposes it suffices to look at these functions visually, Figures 2A and 2B.

**Figure 2:**
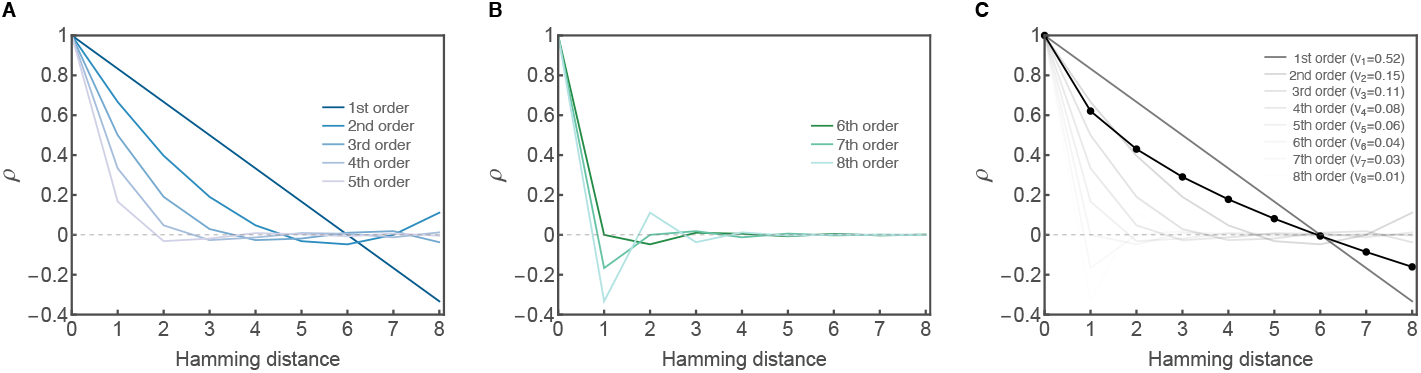
The relationship between the distance correlation function (*ρ*) and higher-order epistasis. The example shown considers sequences of length 8 with 4 alleles per site. (A) Distance correlation functions for interaction orders contributing to positive local correlations. (B) Distance correlation functions for interactions contributing to negative local correlations. The critical interaction order dividing these two groups of interactions is given by the expected number of mutational differences between two random sequences (which is 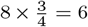 in this case). Note that the values plotted in (A) and (B) do not depend on the specifics of the genotype-phenotype map, but rather depend only on the sequence length *ℓ* and number of alleles per site. These values are given by the appropriately normalized Krawtchouk polynomials (*SI Appendix*). (C) The distance correlation function of a complete genotype-phenotype map is a weighted average of curves shown in (A) and (B), with the weights given by the list of variance components (*v*_*k*_, *k* = 1, ···, *ℓ*, shown in parentheses in the legend). The distance correlation function and weights shown correspond to the genotype-phenotype map from Figure 1 and each curve is shaded proportionally to its weight.

We see that these functions split naturally into two groups, with different qualitative interpretations depending on the order of the interaction. The first of these groups consists of orders of interaction that contribute positive local correlations, so that genotypes that are near to each other in sequence space tend to have similar phenotypes. These functions are shown in Figure 2A, and we can see that the main qualitative effect of increasing interaction order among this group is that these locally positive correlations decay increasingly rapidly.

The second of these groups consists of orders of interaction that exhibit negative local correlations, i.e., they make mutationally adjacent sequences tend to have anti-correlated values. These orders of interaction are shown in Figure 2B, and qualitatively they oscillate increasingly rapidly as the order increases. There is an easy mnemonic for whether a particular interaction order contributes locally positive or locally negative correlations, since if the order of interaction is less than the expected distance between two random sequences, then that order contributes locally positive correlations, otherwise it contributes negative local correlations. (If the order is equal to the expected distance between random sequences, then the correlation at distance 1 is zero, but it will be negative at distance 2, see *SI Appendix*).

It turns out that the distance correlation function of a complete genotype-phenotype map is simply a weighted average of these curves, with the weights given by the list of variance components (illustrated by Figure 2C). Therefore, having Figure 1B allows us to draw Figure 1A. Conversely, given the distance correlation in Figure 1A, we can identify the unique linear combination of curves in Figure 2A-B that produce it, which allows us to draw Figure 1B. At the same time, given the mean squared regression coefficient *λ*_*k*_ for each order *k*, we can calculate the total variance due to *k*-th order interaction by multiplying *λ*_*k*_ with the number of interaction terms of order *k*. This allows us to convert between the quantities in Figure 1B and C. Finally, once we have the vector of *λ*_*k*_, we can calculate the distance correlation function for how local genetic interactions of order *j* generalize over sequence space, *γ*_*j*_. Specifically, we find that the contribution of *k*th-order interaction to *γ*_*j*_ takes a fixed shape that is given by a Krawtchouk polynomial of order *k* − *j* for a sequence of length *ℓ* − *j*, so that the calculation of the curve *γ*_*j*_ is exactly the same as calculating the distance correlation function for a sequence of length *ℓ* − *j* whose *ℓ* − *j λ*_*k*_s are simply *λ*_*j*_,…, *λ*_*ℓ*_ (see Supplemental Figure S1 for an example and *SI Appendix* for details).

In summary, we see that the reason the fraction of variance due to each order of interaction determines how phenotypic observations, mutational effects, local double-mutant epistatic coefficients, etc. generalize is that each order of interaction makes a characteristic contribution to the shape of the distance correlation functions for each of these quantities. Moreover, the qualitative nature of these contributions is that the correlations decay more and more quickly with genetic background as the order of interactions increases, so that we can think of an increasing fraction of higher-order interactions as resulting in an increasingly rapid decay of predictability with genetic distance, with the exact values of the variance components determining the precise shape of the decay.

### Bayesian phenotypic prediction

Now that we understand how higher-order epistasis relates to how far in sequence space we can generalize our experimental observations, our next goal is to develop an inference procedure that can incorporate our specific beliefs about how quickly phenotypic predictability decays. One way to incorporate such beliefs is via Bayesian inference, where these beliefs can be incorporated by choosing an appropriate prior. In this context, the prior takes the form of a probability distribution over all possible genotype-phenotype maps, and we wish to construct a prior whose mass is concentrated on genotype-phenotype maps where the phenotypic predictability decays in the desired manner.

In fact, our analysis so far suggests a natural family of such priors. Specifically, given the fraction of variance due to each order of interaction (as shown schematically in Figure 1B), we can draw epistatic interaction coefficients from a zero-mean normal distribution with variance given by the values in Figure 1C, which results in a genotype-phenotype map that in expectation produces the patterns of correlation shown in Figure 1A and Figure 1D-F. Models of this form are actually well-studied in the theoretical literature on fitness landscapes, where they are known as “random field models” [53, 56], and take the form of a family of multivariate Gaussian distributions parameterized in terms of the amount of variance due to each order of genetic interaction.

Importantly, various previously developed methods can be subsumed as particular (limiting) cases of Bayesian inference under this family of priors. For example, the solutions of the additive model and our recently proposed method of minimum epistasis interpolation [59] both arise as the maximum a posteriori (MAP) estimates in particular limiting cases where the prior fraction of variance due to additive effects goes to 1 (see Supplemental Figure S2). Similarly, the pairwise model [51] arises as the MAP estimate in the limit where the total fraction of variance due to additive and pairwise effects goes to 1 (Supplemental Figure S2). Thus, in a rigorous manner we can view these previously proposed methods as encoding specific assumptions about how the predictability of mutational effects, epistatic coefficients and phenotypic values changes as we move through sequence space, where these assumptions take the form of particular shapes for the curves in Figure 1 or equivalently specific prior assumptions about the fraction of phenotypic variance due to each order of interaction.

Finally, a key fact about this family of priors is that they are Gaussian, and so under the assumption that experimental errors are normally distributed, we can do inference under this prior using Gaussian process regression (see [47] for a review), which allows us to write down analytical expressions for the corresponding posterior distribution. In particular, suppose our prior distribution is a mean zero Gaussian with covariance matrix **K**, **y** is our vector of observations and **E** is a diagonal matrix with estimates of the variance due to experimental noise for each of our observations down the main diagonal. Then the posterior distribution for our vector of predicted phenotypes **f** is normally distributed with mean

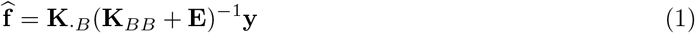

and covariance matrix

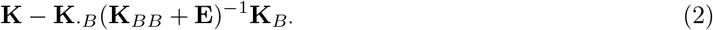

where **K**_*BB*_ is the submatrix of **K** indexed by the set of observed sequences B, and **K**_*B·*_ and **K**_*·B*_ are the submatrices of **K** consisting, respectively, of the rows and columns indexed by members of B.

### Estimating variance components from partial data

To summarize the previous section, if we know the fraction of phenotypic variation due to each order of epistatic interaction, then we can derive a simple method of making phenotypic predictions that uses the corresponding covariance structure to appropriately generalize from observed phenotypic effects, double-mutant epistatic interactions, phenotypic values, etc. While several existing methods of phenotypic prediction essentially come down to making specific assumptions about these variance components, our analysis suggests that a natural approach would be to make our predictions using variance components estimated from the data itself, i.e., an empirical Bayes approach in which we determine what prior to use by looking at the covariance structure of our observations. Conceptually, we want to make phenotypic predictions by assuming that the decay in the predictability of mutational effects, local epistatic interactions, etc. for the true genotype-phenotype mapping is similar outside our data as it is within our data. Practically, we can implement this idea by doing inference under a prior consisting of random genotype-phenotype maps where the effects of mutations (and epistatic coefficients, etc.) decay with genetic distance in the same way as in our data.

A naïve implementation of this approach would be to simply use our observed distance correlation function to build the covariance matrix **K** for our prior by setting the covariance between each pair of sequences at distance d equal to the covariance between sequences at distance d in our data. However, there is a subtle problem with this idea. To see what the difficulty is, recall that the distance correlation in phenotypes is a weighted sum of distance correlation curves of various orders with the weights equal to the fraction of variance due to each order of interaction, as shown in Figure 2.

The fact that these weights need to be positive and sum to one puts strong constraints on the shape that the correlation function can take for a function defined over all of sequence space. For example, positive local correlations cannot decay any more slowly than they would for a purely additive model. However, for incompletely sampled sequence spaces, these constraints need not hold (e.g., if the sampling consisted of several clusters of sequences with identical phenotypes separated from each other with missing sequences, one could have a perfect correlation within the smaller distance classes). Unfortunately, using such a function to define a the matrix **K** would not result in a valid prior (in particular, **K** would not be positive definite, see *SI Appendix*). Thus, rather than using the observed covariance function to define our prior, we instead attempt to find the closest valid prior. We do this using weighted least squares, where the squared error for the correlation at distance *d* is weighted by the number of pairs of sequences at distance *d* (*SI Appendix*); this technique is formally equivalent to the idea of choosing a prior based on “kernel alignment” in the Gaussian processes literature, see ref. [60]. In addition, when producing this weighted least squares estimate, we also take into account the magnitude of the experimental noise so as to distinguish between experimental uncertainty and the influence of any true uncorrelated component of the genotype-phenotype map, and apply regularization to ensure that the resulting prior includes interactions of all orders (*SI Appendix*).

### Practical implementation

So far we have described a two-step method for phenotypic prediction: in the first step we estimate the variance components of the data via a weighted least squares fit to the observed phenotypic correlation function; then, in the second step, we conduct Gaussian process regression under the corresponding prior, so as to encapsulate the assumption that the effects of mutations, etc. generalize outside the data in a manner similar to how they generalize within the data. However, to construct a practical implementation of this approach we must also address certain computational difficulties that arise as a consequence of the very large size of biological sequence space.

Specifically, the major challenge in solving Eq. 1 and 2 is that the computation involves inverting the *m* × *m* dense matrix **K**_*BB*_, where m is the number of observed sequences. The complexity of this problem scales cubically with *m* in time and quadratically with *m* in space. As a result, Gaussian process regression becomes computationally expensive when the number of observed sequences *m* is larger than several thousand [61].

To circumvent this difficulty, we provide an implementation that leverages the symmetries of sequence space to allow practical computations for sequence spaces containing up to low millions of sequences. The basic strategy is to rephrase our problem so that the solution can be found iteratively using only sparse matrix-vector multiplication.

In particular, notice that Eq. 1 can be solved by first finding a vector **α**that satisfies (**K**_*BB*_ +**E**)**α**= **y**. Also, notice that the matrix **K**_*BB*_ is a principle submatrix of **K**, so that we can write **K**_*BB*_ = (**I**_*·B*_)^T^**KI**_*·B*_ where **I**_*·B*_ consists of the columns of the identity matrix **I** that correspond to our set of observed sequences *B*. Since the entries of **K** depend only on the Hamming distance between the corresponding sequences, **K** can expressed as a polynomial in the graph Laplacian (i.e., the matrix **L** whose *i*, *j*-th entry is −1 if *i* is adjacent to *j*, *ℓ*(**α** − 1) if *i* = *j*, and 0 otherwise) that is, 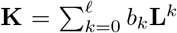 [62, 63] for some *b*_0_,…, *b*_*ℓ*_ that we can find analytically, and thus **Kx** can be found by iteratively applying the sparse matrix **L** to **x** at most *ℓ* times. Using these results, we can rewrite our original equation (**K**_*BB*_ + **E**)**α**= **y** using only sparse matrices as 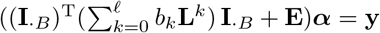, which we solve using the conjugate gradient algorithm.

### Application to protein G

We first apply our method to a dataset derived from a deep mutational scanning study of the IgG-binding domain of streptococcal protein G (GB1) [37]. This experiment assayed nearly all possible combinations of mutations at four sites (V39, D40, G41, and V54; 20^4^ = 160, 000 protein variants), where pairs of mutations at these sites were previously shown to exhibit strong interactions [7]. The library of protein variants was sequenced before and after binding to IgG-Fc beads and the binding scores were determined as the log enrichment ratio (logarithm of ratio of counts before and after selection, normalized by subtracting the log ratio of the wild-type). Due to incomplete coverage of the input library, these data lack binding scores for 6.6% of possible variants.

We began by inferring the variance components of the GB1 landscape from the empirical autocorrelation function (Figure 3A) using our least squares procedure applied to the empirical distance correlation function of all available data (93.6% of all possible sequences, see also *SI Appendix* for details). In Figure 3B, we see that the vast majority of the variance in the data is estimated to be explained by the additive, pairwise, and 3-way components (59%, 37%, and 4% of total variance, respectively), while the estimated contribution of the 4th order component is negligible.

**Figure 3:**
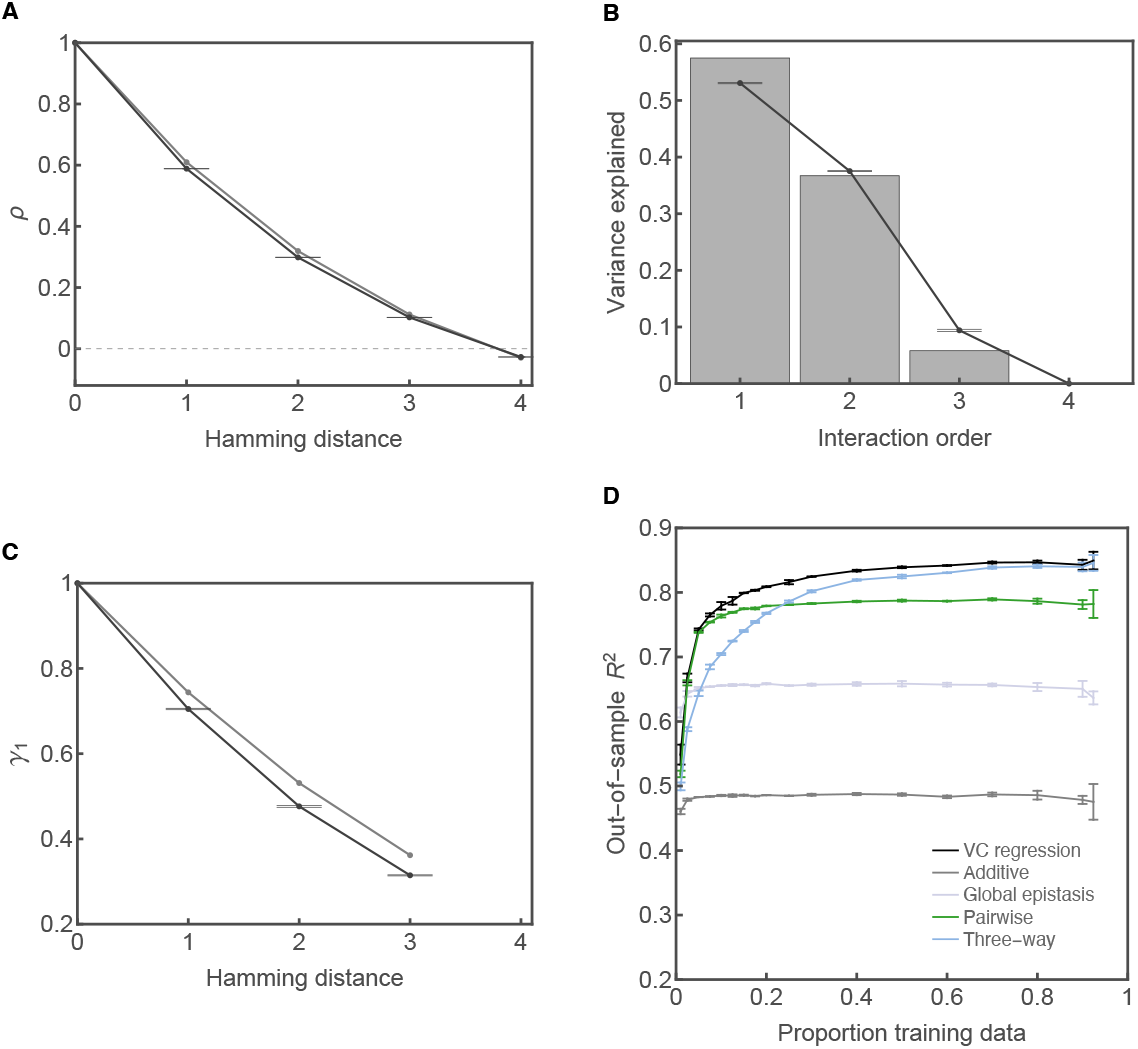
Analyses of the GB1 combinatorial mutagenesis dataset [37]. (A) Distance correlation of phenotypic values. (B) Variance components. (C) Distance correlation of the effects of single mutations. In A-C, gray represents statistics of the prior distribution inferred from the full dataset consisting of 149, 361 genotypes (93.6% of all possible sequences), black represents the posterior statistics estimated based on 2, 000 Hamiltonian Monte Carlo samples. Error bars indicate 95% credible intervals. (D) Comparison of model performance in terms of out-of-sample *R*^2^ for a range of training sample sizes calculated for 5 replicates. Additive models were fit using ordinary least squares. Pairwise and 3-way regression models were fit using elastic net regularization with regularization parameters chosen by 10-fold cross-validation (*SI Appendix*). The global epistasis model assumes the binding score is a nonlinear transformation of an unobserved additive phenotype and was fit following ref. [45]. Error bars represent one standard deviation.

We can use the results from the previous section to understand the practical meaning of these estimates for our task of phenotypic prediction. For example, in Figure 3C, we plot the correlation of mutational effects as a function of Hamming distance [55] (*SI Appendix*). We observe that the correlation of the effect of a random mutation is 0.72 between two genetic backgrounds that differ by one mutation, and 0.32 for two maximally distinct backgrounds (Hamming distance = 3). This decay is characteristic of non-additivity and shows that while the effects of point mutations remain positively correlated across sequence space, the extent of this correlation is approximately twice as high in nearby sequences as opposed to maximally distant sequences, and that therefore, when making predictions, we should give local observations of mutational effects approximately twice as strong a weight as distant observations of mutational effects.

At a broader scale, our analysis above also provides qualitative insights into the overall structure of the genotype-phenotype map. For example, we stated above that the orders of epistatic interaction can be divided into locally correlated and locally anti-correlated groups, depending on whether the order of the interaction is less or greater than the expected distance between two random sequences. Random protein sequences of length 4 differ at 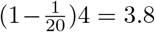 sites on average, so interaction orders 1 through 3 correspond to the genotype-phenotype map being locally correlated, whereas order 4 controls the strength of local anti-correlation. Thus, our estimated variance components suggest that the GB1 genotype-phenotype map is dominated by locally positive correlations, with essentially no anti-correlated component.

Within our overall inference procedure, the estimated variance components discussed above are used to construct a prior probability distribution over all genotype-phenotype maps where, in expectation, mutational effects, epistatic interactions and observed phenotypes generalize across sequence space in the same manner as observed in the data. The resulting prior distribution is based only based on the coarse summary statistics provided by the distance correlation function, and so the next step is to calculate the posterior distribution using the fine-scale information from the individual observations. Specifically, we calculated the MAP solution using all available data and drew 2000 samples from the resulting posterior distribution. Using a 2021 MacBook laptop with 32 Gb RAM, calculating the MAP solution took less than one minute, and we could also produce samples from the posterior distribution at a rate of 1-2 samples per minute.

One immediate question about this posterior distribution is the extent to which its distance correlation function and variance components are similar or different from that of the prior (Figure 3A-C). We find that the posterior gives very tight estimates of the variance components and correlation structure of the true genotype-phenotype map, but that these estimates differ somewhat from the prior, with the 3rd order interactions being roughly 1.6 times as strong in the posterior (Figure 3B), which results in a slightly faster decay in the predictability of mutational effects as we move through sequence space (Figure 3C). Thus, we conclude that our prior distribution provided a qualitatively reasonable estimate of the overall statistical features of the data, but that the rough estimates used to define the prior can be further refined using our full inference procedure.

Obviously, another important question is the performance of the predictions made by our method. Since the GB1 landscape is relatively well sampled, we were able to assess this performance for a large range of sampling regimes, from quite sparse to extremely dense, by using our method to make predictions for randomly sampled held-out data with increasing amounts of training data. Critically, the variance component estimates were re-computed for each of these random samples in order to provide a realistic test of the entire inference pipeline in the low-data regime. For comparison we also fit an additive model using ordinary least squares and regularized pairwise and 3-way regression models. Since both *L*_1_ and *L*_2_ regularized regression have been used to infer genotype-phenotype maps [39, 51, 64–66], here we fit the pairwise and three-way models using elastic net regression (see *SI Appendix*) where the penalty term for model complexity is a mixture of *L*_1_ and *L*_2_ norms [67] and the relative weight of the two penalties is chosen via crossvalidation. In addition to the linear regression models, we also fit a global epistasis model [45] where the binding score is modeled as a nonlinear transformation of a latent additive phenotype on which each possible mutation has a background-independent effect (*SI Appendix*).

We compared the predictive accuracy of these five models by plotting out-of-sample *R*^2^ against a wide range of training sample sizes (Figure 3D). We see that the out-of-sample *R*^2^ of the additive model and the global epistasis model stay nearly constant, regardless of training sample size, consistent with their low number of model parameters and low flexibility. The modest *R*^2^ for the global epistasis model also indicates a substantial degree of specific epistasis (i.e., interactions between specific subsets of sites, as opposed to interactions due to a global nonlinearity [34]). In terms of the regression models that do include these specific interactions, the pairwise model is among the top models for low training sample size, but fails to improve beyond 20% training data, while the 3-way model performs strongly with a large amount of data, but under-performs when data are sparse. We see that our empirical variance component regression (VC regression) method performs equivalently to the pairwise model at low data density and similarly to the three-way model at high data density (remaining marginally superior at very high sampling), and thus provides the strongest overall performance.

### Application to human 5′ splice site data

To provide an application of our method to a nucleic acid genotype-phenotype map, we turn to an analysis of a high-throughput splicing assay that measured the activity of nearly all possible 5′ splice sites [31]. The 5′ splice site (5′ss) is a 9-nucleotide sequence that spans the exon-intron junction. It comprises 3 nt at the end of the upstream exon (denoted as positions −3 to −1) and 6 nt at the beginning of the intron (coded +1 to +6). The consensus 5′ss sequence in humans is CAG/GUAAGU, with the slash denoting the exon-intron junction. At the beginning of the splicing reaction, the 5′ss is recognized by the U1 snRNP of the spliceosome through direct base pairing between the 5′ss and the U1 snRNA [68], whose 5′ sequence is complementary to the consensus 5′ss sequence. In ref. [31], the authors used a massively parallel splicing assay to estimate the splicing efficiency of 93.8% of the 32,768 possible 5′ss sequences of the form NNN/GYNNNN for intron 7 of the gene *SMN1*, using a barcoded minigene library transiently transfected into human cells. Splicing efficiency was measured in units of percent spliced in (PSI), which was estimated as the ratio between the exon inclusion read count and the total read count (which comprises both exon inclusion and exon skipping reads) divided by the corresponding ratio for the consensus sequence, expressed as a percentage. Computational times were somewhat faster for this dataset than for GB1, with the MAP estimate taking roughly 25 seconds on a 2021 MacBook and samples from the posterior distribution being produced at a rate of roughly 10 samples per minute.

Figure 4A shows the distance correlation function of PSI for the observed sequences. These correlations drop off quite rapidly, with sequences differing at 5 or more positions having PSIs that are essentially uncorrelated. The associated estimated variance components are shown in Figure 4B. These indicate that pairwise interactions account for the largest proportion of the sample variance (42.2%), but that there are also substantial higher-order interactions, with the variance due to 5-way interactions (13.7%) being comparable to those of the additive and three-way component. The orders of genetic interaction corresponding to locally negative correlations (order ≥ 6, since the Hamming distance between two random sequences is equal to 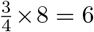) are estimated to play a relatively small but perhaps non-negligible role, accounting for 6.0% of the total variance. In Figure 4C, we found the correlation of mutational effects for two backgrounds that differ by one mutation is roughly 50% but decays to roughly zero for distant genetic backgrounds. Sampling from the posterior distribution, we see that the statistical characteristics of the posterior again have very small credible intervals and remain similar to those estimated using our least squares procedure, with a slightly increased contribution of pair-wise and three-way interactions and a decreased contribution of five-way interactions (Figure 4B). Overall, the splicing landscape appears to be dominated by interactions of order 2 through 5, resulting in positive correlations between the splicing activity of nearby genotypes, but a relatively limited ability to generalize our observations to distant regions of sequence space. This qualitative behavior is consistent with our mechanistic intuition, e.g., mutations that substantially decrease U1 snRNA binding in the context of a functional splice site are likely to have no impact in an already non-functional sequence context.

**Figure 4:**
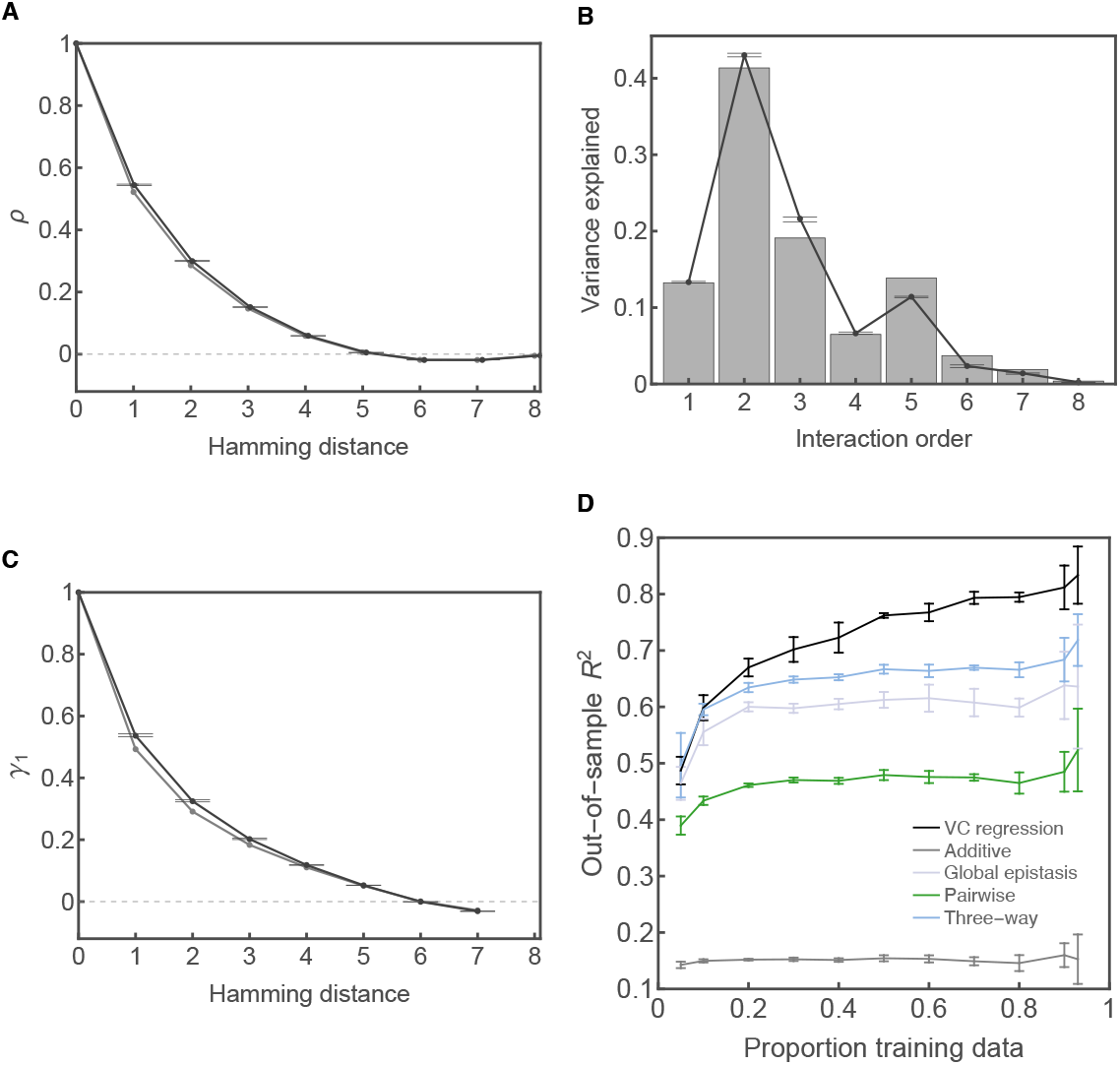
Analyses of the *SMN1* 5′ss combinatorial mutagenesis dataset. (A) Distance correlation function of the splicing phenotype (PSI). (B) Variance components. (C) Distance correlation of single-mutant effects. Gray represents statistics of the prior distribution inferred from the full dataset consisting of 30, 732 genotypes (93.8% of all possible splice sites), black represents the posterior statistics estimated using 2000 Hamiltonian Monte Carlo samples. Error bars indicate 95% credible intervals. (D) Out-of-sample *R*^2^ of the five models plotted against a range of training sample sizes. Error bars represent one standard deviation calculated for 5 replicates for each sample size.

To see how our model performs when greater or lesser amounts of data are available, we compared the predictive power of the same five models as in Figure 3D by plotting their out-of-sample *R*^2^ against a wide range of training sample sizes (Figure 4D). The rank order of the models is largely consistent throughout the sampling range. More importantly, we see that the variance component model adapts to increasing data density at a much faster rate than the other models. For example, with low sampling density (training sample size < 20% of all possible sequences) the three-way model has similar performance as our model, but the performance gap between the two models quickly widens as the training data become dense. The variance component model is able to achieve a final *R*^2^ = 0.83 with 93% of the sequence space assigned as training data (*n* = 30474), compared with the three-way model *R*^2^ = 0.72. This difference in model performance is consistent with the observation of substantial contribution of higher-order interactions (*k* > 3), which the low-order regression model is unable to accommodate.

Another question is the qualitative nature of the genetic interactions captured by our model. We note that the global epistasis model provides a remarkably good fit to the data, considering that it has only a few more parameters than a simple additive model. In Supplemental Figure S3, we see that the global epistasis model approximates the splicing landscape with a sigmoid-like function that maps an unobserved additive trait to the PSI scale. This is as we might expect under a simple biophysical model where each position in the splice site makes a context-independent contribution to the binding energy of the U1 snRNA to the 5′ss, and then this binding energy is mapped via a nonlinear function to PSI [3]. However, we also note that this simple model fails to capture some important features of the data, most notably a group of false-negative sequences that are predicted to be non-functional by the global epistasis model but experimentally show moderate to high measured PSI (Supplemental Figure S4A). Using the variance component regression, we were able to accurately predict these outlier sequences (Supplemental Figure S4B). We thus conclude that while the global epistasis model provides a relatively simple first-pass understanding of the landscape of 5′ss activity, our empirical variance component regression is able to capture more of the fine-scale features of this particular genotype-phenotype map.

Although predictions on held-out data provide one means of testing model performance, a stronger test is to conduct low-throughput experiments to validate the predictions of our method on sequences that were not measured in the original experiment. The *SMN1* dataset provides a suitable case study for this application, since the original dataset does not report the PSI of 2036 sequences (6.2% of all possible 5′ss) due to low read counts. To assess the predictive power of our method for these truly missing sequences, we first made predictions for all unsampled sequences using all available data. We then selected 40 unsampled sequences whose predicted values are evenly distributed on the PSI scale. The true PSIs of these sequences were then quantitatively measured using low-throughput radioactive RT-PCR [31]; see *Materials and Methods*and Figure 5A. Overall, our method achieved a reasonable qualitative agreement with the low-throughput measurements (Figure 5B), but differs systematically in that the transition between nearly 0 and nearly 100 PSI is more rapid in the low-throughput measurement than in our predictions. Intuitively, we can understand the source of this discrepancy in terms of the geometry of the splicing landscape, which features a bimodal distribution of PSIs with separate modes near 0 and 100 [31] and a sharp transition between these two sets of sequences in sequence space (Supplemental Figure S3). Because phenotypic observations generalize farther in most regions of sequence space than they do near this boundary between low and high PSI, our method tends to smooth anomalously sharp features of this type. This results in out-of-sample predictions that are more smoothly graded, rather than threshold-like, in the vicinity of this boundary.

**Figure 5:**
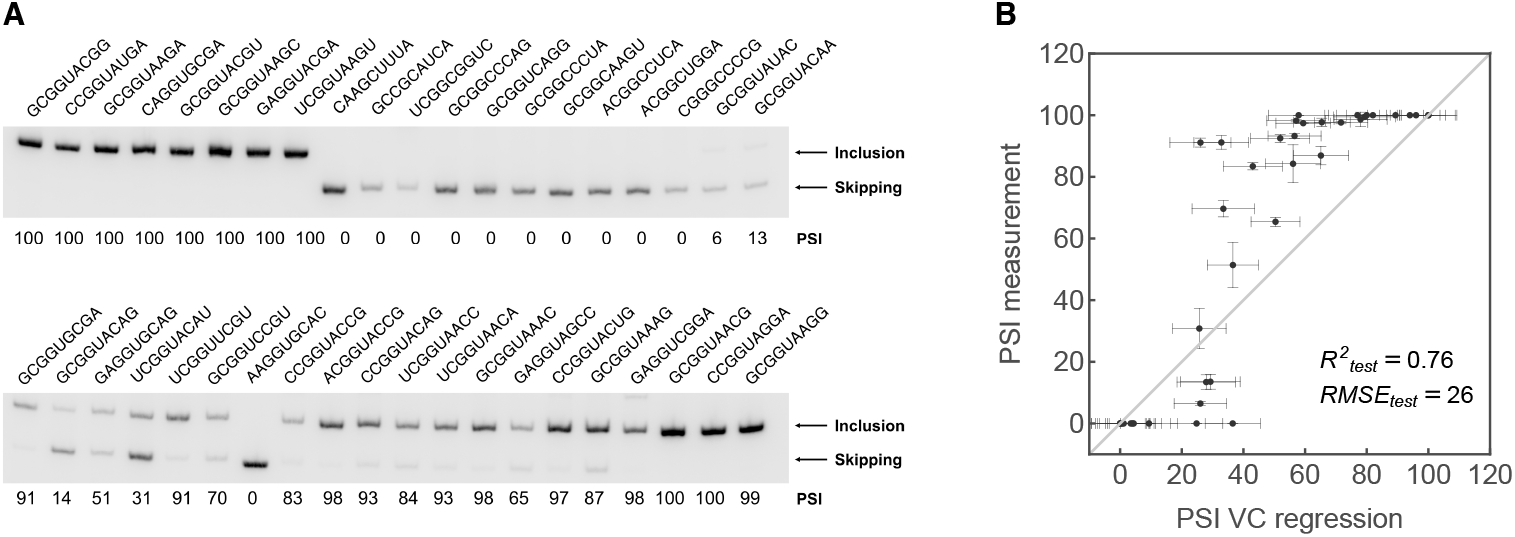
Manual validation of predicted PSI for 40 unmeasured *SMN1* 5′ss. (A) Gel images of manually validated sequences. For each lane, the top band corresponds to mRNA product containing exon 7 (exon inclusion), while the bottom band correspond to mRNA product without exon 7 (exon skipping). Percent spliced in (PSI) is indicated below each lane. Gel images are representative of triplicates. (B) Scatterplot showing measured PSI values versus PSI values predicted by the variance component regression. Horizontal error bars correspond to one standard deviation of the posterior distribution. Vertical error bars correspond to one standard deviation around the mean PSI estimated using three replicates in the manual validation. Since unlike the high-throughput measurements, the low-throughput PSIs are inherently restricted to the range 0–100, in this analysis we likewise cap the predicted PSIs to lie in this same range (see Supplemental Figure S5 for the unrestricted predictions).

### Structure of the *SMN1* splicing landscape

Besides making accurate phenotypic predictions, it is important to understand the qualitative features of a genotype-phenotype map, both with regard to how the underlying mechanisms result in observed genetic interactions and how these genetic interactions affect other processes, such as molecular evolution and disease. For simple models, such as pairwise interaction models or global epistasis models, extracting these qualitative insights can often be achieved by examining the inferred model parameters. Here, we take a different approach and attempt to understand these major qualitative features by constructing visualizations based on the entire inferred activity landscape. Because we previously conducted a detailed analysis of this type for the GB1 dataset [see 59], we focus here on the inferred activity landscape for the 5′ss.

In particular, our visualization method [69] is based on constructing a model of molecular evolution under the assumption that natural selection is acting to preserve the molecular function measured in the assay. The resulting visualization optimally represents the expected time it takes to evolve from one sequence to another (*SI Appendix*), and naturally produces clusters of genotypes where the long-term evolutionary dynamics are similar for a population starting at any genotype in that cluster (e.g., genotypes on the slopes leading up to a fitness peak will tend to be plotted near that peak). To make such a visualization for our splicing data, we built a model of molecular evolution based on the MAP estimate obtained above (*SI Appendix*). We then used the subdominant eigenvectors of the transition matrix for this model as coordinates for the genotypes in a low-dimensional representation; these coordinates are known as diffusion axes [70], since they relate closely to how the probability distribution describing the genotypic state of a population evolving under the combined action of selection, mutation, and genetic drift is likely to diffuse through sequence space [69, 71].

The resulting visualization using the first three diffusion axes is shown in Figure 6A and B. Here, genotypes are points (colored by the number of times that particular 5′ss is used in the human genome, *Materials and Methods*) and edges connect genotypes that differ by single point mutations. Remarkably, each of these diffusion axes turns out to have an interpretable meaning. First, diffusion axis 1 separates functional 5′ss (large positive values) from non-functional 5′ss (negative values), as can be seen in Figure 6C, which plots the estimated PSI against diffusion axis 1. Second, diffusion axis 2 captures the typical physical location of consensus nucleotides within high-activity 5′ss. Specifically, as one moves up diffusion axis 2, the mean position of consensus nucleotides shifts from the exonic portion (5′-end) of the splice site to the intronic (3′) portion (Figure 6D). This reflects a previously observed “seesaw” linkage pattern [72–75] between the intronic and exonic portions of the splice site, where non-consensus nucleotides are typically clustered in one or the other of these regions, but not both. Finally, we find that diffusion axis 3 encodes whether or not mutations are present at the +3 position (Figure 6B inset), where 5′ss with the consensus A are plotted at negative values on diffusion axis 3, and 5′ss with mutant nucleotides on +3 are plotted at positive values of diffusion axis 3.

**Figure 6:**
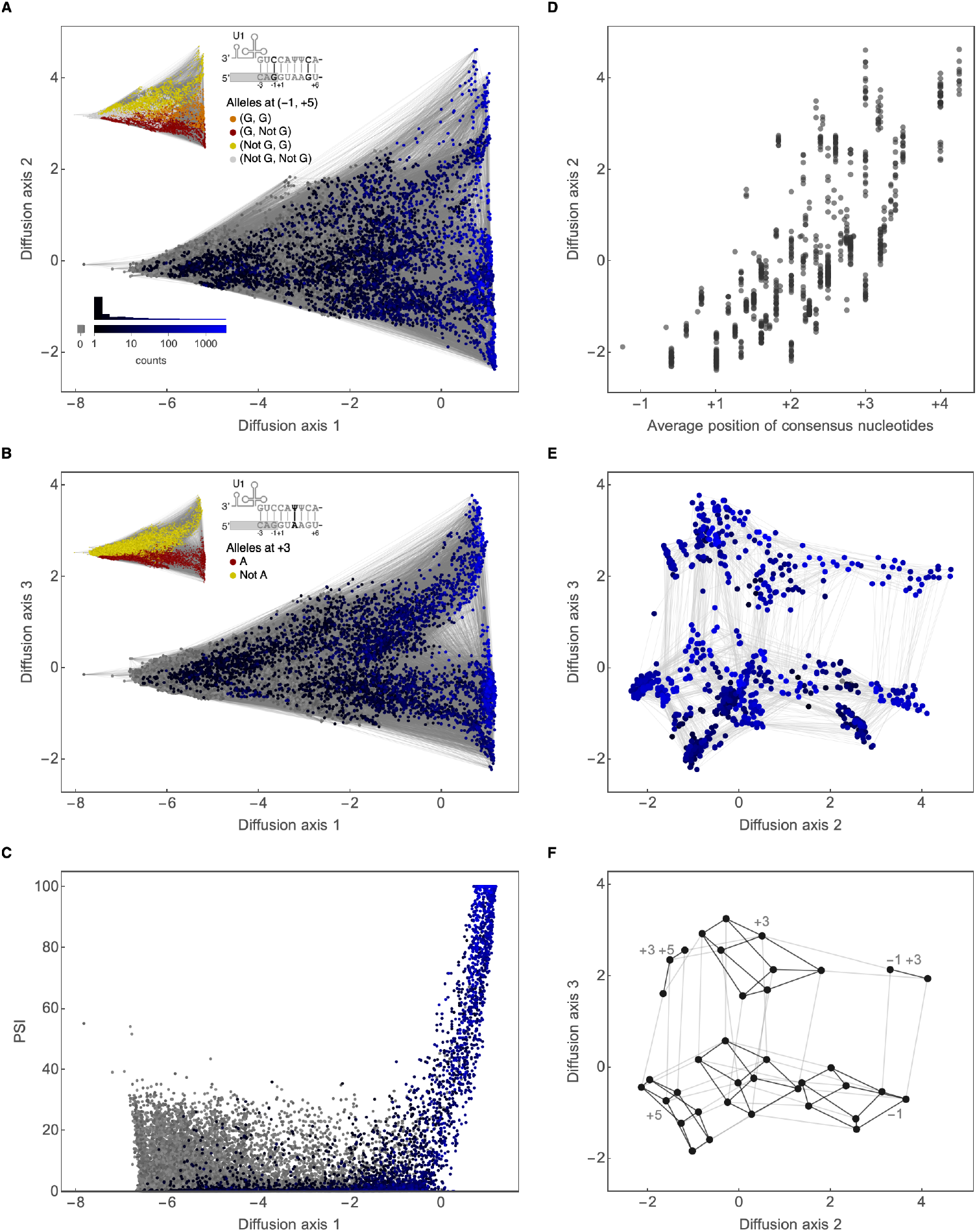
Visualization of the *SMN1* splicing landscape reconstructed using Empirical Variance Component regression. Genotypes are plotted using the dimensionality reduction technique from [69] (see *SI Appendix*). (A) Visualization of all 32,768 splice sites using diffusion axes 1 and 2. Two splice sites are connected by an edge if they differ by a point mutation. (B) Visualization of all 32,768 splice sites using diffusion axes 1 and 3. (C) PSI versus diffusion axis 1; we see that diffusion axis 1 separates high PSI versus low PSI sequences. Genotypes are colored in (A-C) according to the number of times they are observed as annotated splice sites in the human reference genome (hg38); see panel (A) inset for the color scale and a histogram of the numbers of counts. Gray dots represent sequences not present as functional splice sites (65.9% of all sequences of the form NNN/GYNNNN). (D) Diffusion axis 2 versus the average physical position of the consensus nucleotides of the 818 splice sites with predicted PSI > 80%. (E) Visualization of the 818 splice sites with predicted PSI > 80% using diffusion axes 2 and 3. Genotypes colored as in (A-C). (F) Abstracted version of panel E. Splice sites are grouped by mutational states (consensus vs. mutated) at positions -1, -2, +3, +4, +5, and +6. Each dot corresponds to a group of sequences with a prescribed pattern of consensus or mutated states on the six sites. Two groups are connected by an edge if they differ in mutational state at exactly one site. Gray lines represent differences at position -1, +3, and +5. Black lines represent differences at positions -2, +4, and +6. Only groups containing splice sites with > 80% PSI are shown, resulting in six (in)complete cubes with black edges, each representing a combination of mutational states on the three major sites -1, +3, and +5. The incompleteness of a cube indicates the absence of a combination of mutational states at position -2, +4, and +6. Note that no cubes contain both -1 and +5 mutant states, indicating a major incompatibility between mutations at these two sites.

To reveal more detailed structures in the sub-landscape of high-activity 5′ss, we focus on the 818 5′ss with PSI>80. These sequences are plotted using diffusion axis 2 and 3 in Figure 6E. Figure 6F further groups these high-activity 5′ss by their mutant states (consensus vs. mutant) at six positions −1, −2, +3, +4, +5, and +6 and represents each group by a dot (see also Supplemental Figure S6). On a coarse level, we see two main groups of 5′ss separated along diffusion axis 3 that correspond to sequences with the canonical A (bottom) and mutant nucleotides (predominantly G, top) at position +3, and also a distinction into three main groups along diffusion axis 2, where the three groups correspond to mutant +5 (left), consensus for both +5 and −1 (center), and mutant −1 (right). This separation between sequences containing mutations at positions +5 and −1 (see also Figure 6A, inset) arises due to a previously noted incompatibility between mutations at these two sites [31, 73–75], so that evolutionary trajectories that maintain high levels of splicing must typically wait for a reversion of a +5 mutation before fixing a mutation at the −1 position (and vice versa).

On a finer scale, Figure 6F and Supplemental Figure S6 show that each functional combination of mutant states on the three major positions (−1, +3, +5) can be thought of as defining a cube (plotted with dark edges) corresponding to the 2^3^ = 8 possible mutant states on the three minor positions (+2, +4, +6). Whereas having either a single −1 or +5 mutation is compatible with having many different combinations of mutations at the three minor positions (complete cubes on the bottom half of Figure 6F and Supplemental Figure S6), in a +3 mutant background combined with either a −1 or +5 mutation, most combinations of minor mutations result in low activity (two partial cubes on the top half of Figure 6F and Supplemental Figure S6). While in broad strokes this pattern is consistent with a global epistasis model in that minor mutations are mostly tolerable except in weak genetic backgrounds, the global epistasis model further requires that any specific mutation that is tolerated in a weaker background must also be tolerated in a stronger background. However, we instead observe a pattern where in weak backgrounds mutations are only tolerable if an adjacent major site is already mutated (Supplemental Figure S6). In particular, −2 mutations are often tolerable in a −1+3 background but never tolerable in a +3+5 background, and +6 mutations are often tolerable in a +3+5 background but not in a −1+3 background. More specifically, the deleterious effect of the +6 mutation in the −1+3 mutant background is almost completely abrogated in the +3+5 mutant background (Supplemental Figure S7), consistent with previous findings [74]. Likewise, we observe that −2 mutations are often tolerated in the −1+3 mutant background (median effect=−18.6 PSI, calculated for sequences with consensus bases on all other positions and with PSI > 80), but have much larger effects when +3+5 are mutated (median effect= 93.8 PSI). The global epistasis model is even more clearly violated by the fact that the +3+5+6 mutant background can also tolerate certain +4 mutations that would not be tolerable in the absence of a +6 mutation (Supplemental Figure S6). Specifically, these +4 mutations result in two highly functional 5′ss: CAG/GUUGUA, which binds to U1 snRNA using a noncanonical geometry known as an asymmetric loop [76], and AAG/GUGGAC, which does not seem to correspond to any known alternative binding geometry, but occurs as an annotated splice site 14 times in the human genome (*Materials and Methods*) and has a high level of splicing activity as confirmed via low-throughput validation (Supplemental Figure S8).

In summary, we conclude that the 5′ss activity landscape contains many qualitatively different types of genetic interactions. At broad scale, the splicing landscape can be understood in light of the global epistasis model, where PSI is modeled as a nonlinear function of an underlying additive phenotype, and interactions between major mutations arise due to the sharp threshold of the global nonlinearity. However, at a finer scale we discover that the effect of a mutation can be strongly modulated by other mutations in ways that are incompatible with the global epistasis model, both due to specific pairwise interactions, such as the interaction between the +5 and +6 positions, but also due to more complex interactions associated with changes in the physical geometry of U1 snRNA binding [76].

## Discussion

In this paper, we address the problem of how to model the complex genetic interactions observed in high-throughput mutagenesis experiments in order to predict phenotypic values for unmeasured genotypes. Our method is based on the simple idea that the type and extent of epistasis that we predict outside our observed data should be similar to the type and extent of epistasis observed in the data itself. We show that this information about the type and extent of epistasis can be extracted from how correlations between phenotypic values decay as one moves through sequence space. Specifically, we show that: (1) this same distance correlation function also determines the degree to which our observations of mutational effects, double-mutant epistatic coefficients, and observed interactions among three or more mutants generalize across increasingly distant genetic backgrounds; and (2) the distance correlation function can be parameterized in terms of the fraction of phenotypic variance due to each order of genetic interaction (i.e., the *ℓ* variance components, where *ℓ* is the sequence length). By estimating these variance components from the data, we can construct a prior distribution over all possible genotype-phenotype maps that is concentrated on the subset of genotype-phenotype maps where the effects of mutations generalize in the same manner as occurs in our observed data. Conducting Bayesian inference under this prior then produces phenotypic estimates that reflect the belief that the extent and types of epistasis in unobserved regions of sequence space are similar to the extent and type of epistasis in regions of sequence space that we have already observed.

One way to understand our contribution here is to see it as an integration between practical Gaussian process-based methods for analyzing genotype-phenotype maps [77] and the classical spectral theory of fitness landscapes [53, 54, 56], which provides the most sophisticated mathematical theory of genetic interactions currently available. Within this theoretical literature, so-called “random field models” identical to the family of priors we propose have been extensively studied [33, 53, 56], and we have leveraged this existing knowledge to craft priors that encode comprehensible beliefs about the structure of high-dimensional genotype-phenotype maps.

Our results here also provide some significant additions to the spectral theory of fitness landscapes that help to provide a more intuitive view of this complex area of mathematical theory. First, we suggest that higher-order epistatic interactions can be qualitatively classified into two types, corresponding to interactions that result in locally positive correlations or locally negative correlations. The idea of an anti-correlated component to a genotype-phenotype map has been discussed previously in the literature in terms of the “eggbox” component [12, 55] which is perfectly anti-correlated between adjacent genotypes (i.e., whether the phenotypic value is high or low flips with each step one takes through sequence space, similar to the alternating peaks and valleys of an egg carton). Our analysis shows that there is actually a whole set of orders of genetic interaction with a similar character, corresponding to all orders of genetic interaction higher than the average number of differences between two random sequences. However, our main interest is in the components that produce locally positive correlations (which appear more likely to arise under most conceivable physical mechanisms), with the balance between these higher-order locally correlated components controlling how precisely phenotypic correlations decay with increasing Hamming distance.

Second, we defined a summary statistic *γ*_*k*_(d) which, beyond simple phenotypic correlations, measures how mutational effects (*k* = 1) or epistatic coefficients (*k* > 1) decay as the distance d between genetic backgrounds increases. The correlation of mutational effects as a function of distance between genetic backgrounds has been previously termed *γ*(*d*), which is used to measure the ruggedness of the landscape [12, 55]. Here we generalize this measure to epistatic coefficients of any order, and show that the distance correlation of epistatic coefficients of order *k* is in fact determined solely by the components of the landscape of order larger than *k* (see *SI Appendix*, where we provide a simple formula showing the relationship between different orders). This result can also help us understand why our method out-performs pairwise and three-way epistatic models. Specifically, we show that models that include only up to *k*-th order epistatic interactions in fact make the very strong assumption that any observed *k*-th order interactions generalize across all genetic backgrounds. Incorporating higher-order interactions is then equivalent to relaxing this strong assumption and allowing these lower-order interactions to change as one moves through sequence space.

Third, in the *SI Appendix* we provide some additional results to better understand the possible geometries produced by any given order of genetic interaction. In particular, we consider the mean phenotype as a function of Hamming distance from some focal genotype, which is a classical coarse descriptor of genotype-phenotype map and fitness landscape structure [e.g., 78–80]. We show that for a pure *k*-th order interaction this mean function is in fact equal to the distance correlation function up to a multiplicative constant. As a consequence of this result, the distance mean function for a model containing up to only *k*-th order terms must be a *k*-th order polynomial, so that e.g., in a pairwise interaction model the mean fitness at a given distance from a focal genotype is always a quadratic function of distance. However from a biological perspective, we might often expect mean fitness to take more complex shapes, such as a sigmoid [45, 81] (which obviously cannot be well-approximated by a quadratic), providing an explanation for the need to incorporate higher-order interactions in order to provide qualitatively reasonable fits.

The method we propose here also has some commonalities with minimum epistasis interpolation [59], another method we recently proposed for phenotypic prediction that includes genetic interactions of all orders, but the two methods differ in their aims. Minimum epistasis interpolation aims to produce a highly conservative reconstruction of the genotype-phenotype map by making the effects of mutations as consistent as possible between adjacent genetic backgrounds. In contrast, empirical variance composnent regression aims to produce a more realistic reconstruction, where the extent and type of epistasis present in the reconstruction should be similar to the extent and type of epistasis present in the data itself. Depending on the needs of the user, both minimum epistasis interpolation and empirical variance component regression can be conducted either in a Bayesian manner or as a form of *L*_2_-regularized regression [82] (where our MAP estimate is equivalent to the *L*_2_ regularized solution, *SI Appendix*). From a regularization perspective, the main difference between these methods is that they penalize different orders of genetic interaction differently, either with a penalty that increases quadratically with the order of interaction, in the case of minimum epistasis interpolation [see also 83], or a penalty determined by the empirically estimated variance components, in the case of empirical variance component regression. In Supplemental Figure S9, we provide a comparison of model performance between these two methods in the GB1 and *SMN1* dataset. We observe that empirical variance component regression consistently outperforms minimum epistasis interpolation in both the GB1 and *SMN1* datasets. This difference in model performance is likely due to the misalignment between the quadratically increasing penalty imposed by minimum epistatsis interpolation and the actual variance components in the data. Overall, empirical variance component regression is likely the superior method if high predictive performance is desirable. On the other hand, minimum epistasis interpolation is a more conservative approach and has many simple theoretical properties [59]. Thus, it should be preferred when theoretical guarantees for model behavior are more important.

One potential limitation of our approach is our choice to select the hyperparameters based on the point estimates supplied by our training data, i.e., by kernel alignment [60]. It may well be possible to produce more accurate predictions by choosing hyperparameters by maximizing the evidence [47] or via a hierarchical Bayesian model where we integrate over our uncertainty in the values of these hyperparameters, at the cost of a much greater computational burden. However, an advantage of the empirical Bayes approach is that it provides a clear conceptual separation between the first step of estimating the type and extent of epistasis present in a set of phenotypic observations, and the second step of using these estimates to make additional phenotypic predictions.

Another limitation concerning empirical variance component regression is that it is unable to explicitly model any overall nonlinearity that may be present in the genotype-phenotype map, i.e., it does not explicitly model nonspecific or global epistasis [3, 34, 35, 45, 84–86]. Rather, empirical variance component regression must learn any such global structure based on consistent patterns in the observations themselves. For instance, whereas the global epistasis model is able to easily handle the saturation of PSI at 0 and 100%, empirical variance component regression must learn these flatter regions based on the consistently small effects of mutations in particular regions of sequence space, rather than via an overall nonlinearity that is assumed by the structure of the model. Incorporating the possibility of such global nonlinearities would be an important extension to the methods presented here.

A final limitation concerns the applicability of the method we propose to very large datasets. In our implementation, we take advantage of the isotropic property of the prior distribution (i.e. that covariance depends only on Hamming distance) and the highly symmetric graph structure of sequence space. This allows us to express the covariance matrix and its inverse as polynomials in the highly sparse matrix known as the graph Laplacian, which makes inference possible on sequence spaces containing up to low millions of sequences. However, due to the exponential growth of biological sequence space as a function of sequence length, practically this still limits us to nucleic acid sequences of length 11 or less, and amino acid sequences of length 5 or less. Using the kernel trick [87], it is possible to work with much longer sequences, but at the cost of only being able to accommodate up to low tens of thousands of observed sequences, due to the resulting dense kernel matrix. Although here we have successfully analyzed datasets that contain tens to hundreds of thousands of sequences, more work is needed to scale the methods proposed here to even larger datasets and sequence spaces.

## Methods

### Low-throughput validation of unsampled *SMN1* 5′ss

To assess the predictive accuracy of our method for the activity of truly unsampled splice sites, we selected 40 5′ss absent in the *SMN1* dataset that are evenly distributed on the predicted PSI scale. We quantified the splicing activities of the selected 5′ss in the context of an *SMN1* minigene that spans exons 6-8 with the variable 5′ss residing in intron 7. The minigene construct is the same as the one used to generate the high-throughput data [31] (minigene sequence is available at https://github.com/jbkinney/15_splicing). Specifically, minigenes containing variable 5′ss were inserted in to the pcDNA5/FRT expression vector (Invitrogen). 1 μg of minigene plasmid was then transiently transfected into HeLa cells, which were collected after 48 hr. RNA was isolated from the minigene-expressing HeLa cells using Trizol (Life Technologies) and treated with RQ1 RNase-free DNase (Promega). cDNA was made using Improm-II Reverse Transcription System (Promega), following the manufacturer′s instructions. The splicing isoforms were then amplified with minigene-specific primers (F: CTGGCTAACTAGAGAACCCACTGC; R: GGCAACTAGAAGGCACAGTCG) and ^32^P-labeled dCTP using Q5 High-Fidelity DNA Polymerase (New England Biolabs) following the manufacturer′s instructions. PCR products were separated on a 5.5% non-denaturing polyacrylamide gel and were detected using a Typhoon FLA7000 phosphorimager. Finally, we used ImageJ (NIH) to quantify isoform abundance, in the process accounting for cytosine content. All 5′ss were assessed in triplicate.

### Acquisition of human 5′ splice sites

Human 5′ss were extracted from GENCODE Release 34 (GRCh38.p13) (available at https://www.gencodegenes.org/human/).

### Code availability

A python command-line interface, vcregression, that implements the Empirical Variance Component method described here is available at https://github.com/davidmccandlish/vcregression.

## Acknowledgements

This work was funded by NIH grant R35GM133613, an Alfred P. Sloan Research Fellowship (awarded to D.M.M.), as well as funding from the Simons Center for Quantitative Biology at Cold Spring Harbor Laboratory. J.B.K. acknowledges support from NIH grant R35GM133777. M.S.W. and A.R.K. acknowledge support from NIH grant R37GM42699.

## References

[1] Patrick C Phillips. “Epistasis—the essential role of gene interactions in the structure and evolution of genetic systems”. In: Nat. Rev. Genet. 9.11 (2008), pp. 855–867.

[2] Dmitry A Kondrashov and Fyodor A Kondrashov. “Topological features of rugged fitness landscapes in sequence space”. In: Trends Genet. 31.1 (2015), pp. 24–33.

[3] Júlia Domingo, Pablo Baeza-Centurion, and Ben Lehner. “The Causes and Consequences of Genetic Interactions (Epistasis)”. In: Annu. Rev. Genomics Hum. Genet. 20 (2019).

[4] Douglas M Fowler et al. “High-resolution mapping of protein sequence-function relationships”. In: Nat. Methods 7.9 (2010), pp. 741–746.

[5] Lea M Starita et al. “Activity-enhancing mutations in an E3 ubiquitin ligase identified by high-throughput mutagenesis”. In: Proc. Natl. Acad. Sci. U.S.A. 110.14 (2013), E1263–E1272.

[6] Daniel Melamed et al. “Deep mutational scanning of an RRM domain of the Saccharomyces cerevisiae poly (A)-binding protein”. In: RNA 19.11 (2013), pp. 1537–1551.

[7] C Anders Olson, Nicholas C Wu, and Ren Sun. “A comprehensive biophysical description of pairwise epistasis throughout an entire protein domain”. In: Curr. Biol. 24.22 (2014), pp. 2643–2651.

[8] Michael B Doud, Orr Ashenberg, and Jesse D Bloom. “Site-specific amino acid preferences are mostly conserved in two closely related protein homologs”. In: Mol. Biol. Evol. 32.11 (2015), pp. 2944–2960.

[9] Anna I Podgornaia and Michael T Laub. “Pervasive degeneracy and epistasis in a protein-protein interface”. In: Science 347.6222 (2015), pp. 673–677.

[10] Karen S Sarkisyan et al. “Local fitness landscape of the green fluorescent protein”. In: Nature 533.7603 (2016), p. 397.

[11] Barrett Steinberg and Marc Ostermeier. “Shifting fitness and epistatic landscapes reflect trade-offs along an evolutionary pathway”. In: J Mol Biol. 428.13 (2016), pp. 2730–2743.

[12] Claudia Bank et al. “On the (un)predictability of a large intragenic fitness landscape”. In: Proc. Natl. Acad. Sci. U.S.A. 113.49 (2016), pp. 14085–14090.

[13] Tyler N Starr, Lora K Picton, and Joseph W Thornton. “Alternative evolutionary histories in the sequence space of an ancient protein.” In: Nature 549.7672 (Sept. 2017), pp. 409–413.

[14] Victoria O Pokusaeva et al. “An experimental assay of the interactions of amino acids from orthologous sequences shaping a complex fitness landscape”. In: PLos Genet. 15.4 (2019), e1008079.

[15] Calin Plesa et al. “Multiplexed gene synthesis in emulsions for exploring protein functional landscapes”. In: Science 359.6373 (2018), pp. 343–347.

[16] Drew S Tack et al. “The genotype-phenotype landscape of an allosteric protein”. In: Molecular systems biology 17.3 (2021), e10179.

[17] Tyler N Starr et al. “Deep mutational scanning of SARS-CoV-2 receptor binding domain reveals constraints on folding and ACE2 binding”. In: Cell 182.5 (2020), pp. 1295–1310.

[18] Louisa Gonzalez Somermeyer et al. “Heterogeneity of the GFP fitness landscape and data-driven protein design”. In: bioRxiv (2021). doi: 10.1101/2021.12.08.471728. eprint: https://www.biorxiv.org/content/early/2021/12/09/2021.12.08.471728.full.pdf. url: https://www.biorxiv.org/content/early/2021/12/09/2021.12.08.471728.

[19] J N Pitt and A R Ferré-D’Amaré. “Rapid Construction of Empirical RNA Fitness Landscapes”. In: Science 330.6002 (2010), pp. 376–379. doi: 10.1126/science.1192001. url: http://www.sciencemag.org/content/330/6002/376.abstract.

[20] José I Jiménez et al. “Comprehensive experimental fitness landscape and evolutionary network for small RNA”. In: Proc. Natl. Acad. Sci. U.S.A. 110.37 (2013), pp. 14984–14989.

[21] Olga Puchta et al. “Network of epistatic interactions within a yeast snoRNA”. In: Science 352.6287 (2016), pp. 840–844.

[22] Chuan Li et al. “The fitness landscape of a tRNA gene”. In: Science 352.6287 (2016), pp. 837–840.

[23] Júlia Domingo, Guillaume Diss, and Ben Lehner. “Pairwise and higher-order genetic interactions during the evolution of a tRNA”. In: Nature 558.7708 (2018), p. 117.

[24] Valerie WC Soo et al. “Fitness landscape of a dynamic RNA structure”. In: PLoS genetics 17.2 (2021), e1009353.

[25] Rupali P Patwardhan et al. “High-resolution analysis of DNA regulatory elements by synthetic saturation mutagenesis”. In: Nature Biotechnology 27.12 (2009), pp. 1173–1175.

[26] Justin B Kinney et al. “Using deep sequencing to characterize the biophysical mechanism of a transcriptional regulatory sequence”. In: Proc. Natl. Acad. Sci. U.S.A. 107.20 (2010), pp. 9158–9163.

[27] Shengdong Ke et al. “Quantitative evaluation of all hexamers as exonic splicing elements”. In: Genome research 21.8 (2011), pp. 1360–1374.

[28] Alexander B Rosenberg et al. “Learning the sequence determinants of alternative splicing from millions of random sequences”. In: Cell 163.3 (2015), pp. 698–711.

[29] Philippe Julien et al. “The complete local genotype–phenotype landscape for the alternative splicing of a human exon”. In: Nat. Commun. 7 (2016), p. 11558.

[30] Shengdong Ke et al. “Saturation mutagenesis reveals manifold determinants of exon definition”. In: Genome Res. 28.1 (2018), pp. 11–24.

[31] Mandy S Wong, Justin B Kinney, and Adrian R Krainer. “Quantitative Activity Profile and Context Dependence of All Human 5′ Splice Sites”. In: Mol. Cell (2018).

[32] Daniel M Weinreich et al. “Should evolutionary geneticists worry about higher-order epistasis?” In: Curr. Opin. Genet. Dev. 23.6 (2013), pp. 700–707.

[33] Johannes Neidhart, Ivan G Szendro, and Joachim Krug. “Exact results for amplitude spectra of fitness landscapes”. In: J. Theor. Biol. 332 (2013), pp. 218–227.

[34] Tyler N Starr and Joseph W Thornton. “Epistasis in protein evolution”. In: Protein Sci. 25.7 (2016), pp. 1204–1218.

[35] Zachary R Sailer and Michael J Harms. “Detecting High-Order Epistasis in Nonlinear Genotype-Phenotype Maps”. In: Genetics 205.3 (2017), pp. 1079–1088.

[36] Zachary R Sailer and Michael J Harms. “High-order epistasis shapes evolutionary trajectories”. In: PLoS Comput. Biol. 13.5 (2017), e1005541.

[37] Nicholas Wu et al. “Adaptation in protein fitness landscapes is facilitated by indirect paths”. In: eLife 5 (2016), e16965.

[38] Julian Echave and Claus O Wilke. “Biophysical models of protein evolution: understanding the patterns of evolutionary sequence divergence”. In: Annu. Rev. Biophys. 46 (2017), pp. 85–103.

[39] Frank J Poelwijk, Michael Socolich, and Rama Ranganathan. “Learning the pattern of epistasis linking genotype and phenotype in a protein”. In: Nat. Commun. 10.1 (2019), pp. 1–11.

[40] Aneth S Canale et al. “Evolutionary mechanisms studied through protein fitness landscapes”. In: Curr. Opin. Struct. Biol. 48 (2018), pp. 141–148.

[41] Daniel M. Weinreich et al. “The Influence of Higher-Order Epistasis on Biological Fitness Landscape Topography”. In: J. Stat. Phys. 172.1 (2018), pp. 208–225. issn: 00224715. doi: 10.1007/s10955-018-1975-3. url: https://doi.org/10.1007/s10955-018-1975-3.

[42] Jay F Storz. “Compensatory mutations and epistasis for protein function”. In: Curr. Opin. Struct. Biol. 50 (2018), pp. 18–25.

[43] Charlotte M Miton, Karol Buda, and Nobuhiko Tokuriki. “Epistasis and intramolecular networks in protein evolution”. In: Current opinion in structural biology 69 (2021), pp. 160–168.

[44] Gloria Yang et al. “Higher-order epistasis shapes the fitness landscape of a xenobiotic-degrading enzyme”. In: Nature Chemical Biology 15.11 (2019), pp. 1120–1128.

[45] Jakub Otwinowski, David Martin McCandlish, and Joshua B Plotkin. “Inferring the shape of global epistasis”. In: Proceedings of the National Academy of Sciences 115.32 (2018), E7550–E7558.

[46] Gautam Reddy and Michael M Desai. “Global epistasis emerges from a generic model of a complex trait”. In: Elife 10 (2021), e64740.

[47] Carl Edward Rasmussen and Christopher K I Williams. Gaussian processes for machine learning. Vol. 1. MIT press Cambridge, 2006.

[48] Bradley P Carlin and Thomas A Louis. Bayes and empirical Bayes methods for data analysis. Vol. 88. Chapman & Hall/CRC Boca Raton, 2000.

[49] Radford M Neal. “MCMC using Hamiltonian dynamics”. In: Handbook of Markov chain Monte Carlo. Ed. by Steve Brooks et al. 2011, pp. 113–162.

[50] R A Fisher. “The Correlation Between Relatives on the Supposition of Mendelian Inheritance”. In: Trans R Soc Edinburgh 52.02 (1918), pp. 399–433.

[51] Trevor Hinkley et al. “A systems analysis of mutational effects in HIV-1 protease and reverse transcriptase”. In: Nature Genetics 43.5 (2011), pp. 487–489.

[52] Manfred Eigen, John McCaskill, and Peter Schuster. “The molecular quasi-species”. In: Adv. Chem. Phys 75 (1989), pp. 149–263.

[53] Peter F Stadler and Robert Happel. “Random field models for fitness landscapes”. In: J. Math. Biol. 38.5 (1999), pp. 435–478.

[54] Edward D Weinberger. “Fourier and Taylor series on fitness landscapes”. In: Biol Cybern 65.5 (1991), pp. 321–330.

[55] Luca Ferretti et al. “Measuring epistasis in fitness landscapes: The correlation of fitness effects of mutations”. In: Journal of Theoretical Biology 396 (2016), pp. 132–143. issn: 10958541. doi: 10.1016/j.jtbi.2016.01.037.

[56] Robert Happel and Peter F Stadler. “Canonical approximation of fitness landscapes”. In: Complexity 2.1 (1996), pp. 53–58.

[57] Peter F Stadler. “Landscapes and their correlation functions”. In: J. Math. Chem. 20.1 (1996), pp. 1–45.

[58] Peter F Stadler. “Fitness landscapes”. In: Biological Evolution and Statistical Physics. Springer, 2002, pp. 183–204.

[59] Juannan Zhou and David M McCandlish. “Minimum epistasis interpolation for sequence-function relationships”. In: Nature communications 11.1 (2020), pp. 1–14.

[60] Tinghua Wang, Dongyan Zhao, and Shengfeng Tian. “An overview of kernel alignment and its applications”. In: Artificial Intelligence Review 43.2 (2015), pp. 179–192.

[61] John P Cunningham, Krishna V Shenoy, and Maneesh Sahani. “Fast Gaussian process methods for point process intensity estimation”. In: Proceedings of the 25th international conference on Machine learning. 2008, pp. 192–199.

[62] Peter F Stadler. “Random walks and orthogonal functions associated with highly symmetric graphs”. In: Discrete mathematics 145.1-3 (1995), pp. 229–237.

[63] DG Higman. “Intersection matrices for finite permutation groups”. In: Journal of Algebra 6 (1967), pp. 22–42.

[64] Daniel M Weinreich et al. “The influence of higher-order epistasis on biological fitness landscape topography”. In: Journal of Statistical Physics 172.1 (2018), pp. 208–225.

[65] Amirali Aghazadeh et al. “Epistatic Net allows the sparse spectral regularization of deep neural networks for inferring fitness functions”. In: Nature communications 12.1 (2021), pp. 1–10.

[66] David H Brookes, Amirali Aghazadeh, and Jennifer Listgarten. “On the sparsity of fitness functions and implications for learning”. In: Proceedings of the National Academy of Sciences 119.1 (2022).

[67] Hui Zou and Trevor Hastie. “Regularization and variable selection via the elastic net”. In: Journal of the royal statistica 67.2 (2005), pp. 301–320.

[68] Yasushi Kondo et al. “Crystal structure of human U1 snRNP, a small nuclear ribonucleoprotein particle, reveals the mechanism of 5′splice site recognition”. In: Elife 4 (2015), e04986.

[69] David M McCandlish. “Visualizing fitness landscapes”. In: Evolution 65.6 (2011), pp. 1544–1558.

[70] Ronald R Coifman and Stéphane Lafon. “Diffusion maps”. In: Appl. and Comp. Harmonic Anal. 21.1 (2006), pp. 5–30.

[71] David M McCandlish. “Long-term evolution on complex fitness landscapes when mutation is weak”. In: Heredity 121.5 (2018), pp. 449–465.

[72] Xavier Roca et al. “Features of 5′-splice-site efficiency derived from disease-causing mutations and comparative genomics”. In: Genome research 18.1 (2008), pp. 77–87.

[73] Ido Carmel et al. “Comparative analysis detects dependencies among the 5′splice-site positions”. In: RNA 10.5 (2004), pp. 828–840.

[74] Chris Burge and Samuel Karlin. “Prediction of complete gene structures in human genomic DNA”. In: Journal of Molecular Biology 268.1 (1997), pp. 78–94.

[75] Gene Yeo and Christopher B Burge. “Maximum entropy modeling of short sequence motifs with applications to RNA splicing signals”. In: Journal of computational biology 11.2-3 (2004), pp. 377–394.

[76] Jiazi Tan et al. “Noncanonical registers and base pairs in human 5′splice-site selection”. In: Nucleic acids research 44.8 (2016), pp. 3908–3921.

[77] Philip A Romero, Andreas Krause, and Frances H Arnold. “Navigating the protein fitness landscape with Gaussian processes”. In: Proc. Natl. Acad. Sci. U.S.A. 110.3 (2013),E193–E201.

[78] Alexey S Kondrashov. “Selection against harmful mutations in large sexual and asexual populations”. In: Genetics Research 40.3 (1982), pp. 325–332.

[79] Santiago F Elena and Richard E Lenski. “Test of synergistic interactions among deleterious mutations in bacteria”. In: Nature 390.6658 (1997), pp. 395–398.

[80] Sebastian Bonhoeffer et al. “Evidence for positive epistasis in HIV-1”. In: Science 306.5701 (2004), pp. 1547–1550.

[81] Shimon Bershtein et al. “Robustness–epistasis link shapes the fitness landscape of a randomly drifting protein”. In: Nature 444.7121 (2006), pp. 929–932.

[82] Alexander J Smola and Risi Kondor. “Kernels and regularization on graphs”. In: COLT. Vol. 2777. Springer. 2003, pp. 144–158.

[83] Wei-Chia Chen et al. “Field-theoretic density estimation for biological sequence space with applications to 5 splice site diversity and aneuploidy in cancer”. In: Proceedings of the National Academy of Sciences 118.40 (2021).

[84] Justin B Kinney and David M McCandlish. “Massively Parallel Assays and Quantitative Sequence– Function Relationships”. In: Annu Rev Genomics Hum Genet 20 (2019).

[85] Karen S Sarkisyan et al. “Local fitness landscape of the green fluorescent protein”. In: Nature 533.7603 (2016), pp. 397–401.

[86] Ammar Tareen et al. “MAVE-NN: Quantitative Modeling of Genotype-Phenotype Maps as Information Bottlenecks”. In: BioRxiv (2020).

[87] Bernhard Schölkopf, Ralf Herbrich, and Alex J Smola. “A generalized representer theorem”. In: International conference on computational learning theory. Springer. 2001, pp. 416–426.

