## Supplementary material for "Higher-order epistasis and phenotypic prediction": SI Appendix

### SI Appendix for "Higher-order epistasis and phenotypic prediction"

#### Contents

**1 Introduction** **2**

**2 Distance mean function of pure $k$th-order interactions** **8**

**3 Summary statistics for empirical genotype-phenotype maps** **10**

**4 Gaussian random field landscapes** **11**

**5 Definition of the epistatic coefficient $\epsilon_C(\mathbf{f})$** **13**

**6 Distance covariance of $k$-th order epistatic coefficients for random fields** **15**

**7 Distance covariance of $k$-th order epistatic coefficients in an empirical landscape** **16**

**8 Relation between the $\lambda_k$ and the distance correlation of $k$-th order epistatic coefficients $\Gamma_k$** **18**

**9 Connection with $L_2$-regularized regression and Minimum Epistasis Interpolation** **19**

**10 Inference of hyperparameters for the prior distribution** **20**

**11 Posterior sampling using Hamiltonian Monte Carlo** **22**

**12 Implementation of regularized pairwise and three-way interaction models** **23**

**13 Symmetry properties of the Krawtchouk Polynomial** **24**

**14 Proof for the classification of interactions into locally correlated and anticorrelated groups** **25**

[2]Current address: Skyhawk Therapeutics, Inc. 35 Gatehouse Drive Waltham, MA 02451
[3]Current address: Department of Physics, National Chung Cheng University, Chiayi 62102, Taiwan (R.O.C.)

**15 Proof for results used in Proposition 1** **26**

**16 Processing of the *SMN1* splicing dataset** **27**

**17 Visualization of the *SMN1* splicing landscape** **27**

### 1 Introduction

The main questions this paper attempts to address are how to think about higher-order epistasis in an intuitive manner and how to use this understanding to build better and more principled inference procedures. In this section, we provide an overall introduction to our mathematical framework for studying genetic interaction (epistasis) and how these results relate to Empirical Variance Component Regression. Specifically, we focus on the logical relationship between several key results that relate epistasis among different number of positions to the geometry of the genotype-phenotype maps these interactions induce. The subsequent sections of the *SI Appendix* contain proofs for claims made in this first section as well as additional details and explanation for results presented in the main text.

##### Epistatic interactions of different orders

We first show that any arbitrary genotype-phenotype map can be expressed as a sum of components, each corresponding to a given order of epistatic interaction. More precisely, let $A$ be an alphabet of $\alpha$ alleles. We use $A^\ell$ to denote the sequence space, i.e. the set of all possible combinations of alleles in $A$ of length $\ell$. Given two sequences $x, x'$ in $A^\ell$, their Hamming distance $D(x, x')$ (number of positions the two sequences differ) provides a straightforward measure of distance over $A^\ell$. Given the data, we would like to construct a model for the genotype-phenotype map. A standard strategy is to expand around a wild type sequence (WT) and model the genotype-phenotype map as a linear combination of the effects of individual mutations [1, 2]:

$$f(x) = \beta_0 + \sum_i \beta_i s_i(x), \tag{1}$$

where $\beta_0$ represents the fitness of the wild type sequence. The sum $\sum_i \beta_i s_i(x)$ captures the total additive effects of mutations, with $\beta_i$ representing the effect of the $i$-th mutation, while $s_i(x)$ is a function that returns 1 if the sequence $x$ contains the mutation and 0 otherwise. In many cases, we need to accommodate not only the additive effects of individual mutations, but also epistatic interactions among mutations on different sites. Thus a possible extension of Eq. 1 is to incorporate interaction between pairs of loci [3]:

$$f(x) = \beta_0 + \sum_i \beta_i s_i(x) + \sum_{i,j} \beta_{ij} s_{ij}(x). \tag{2}$$

Analogously, $\sum_{i,j} \beta_{ij} s_{ij}(x)$ models the effects of all possible pairwise interactions among mutations, in that $\beta_{ij}$ models the effect of epistasis between mutations $i$ and $j$, $s_{ij}(x)$ encodes the presence or absence of $i$ and $j$ combination on a pair of loci. We can further extend this model to allow higher-order interactions:

$$f(x) = \beta_0 + \sum_i \beta_i s_i(x) + \sum_{i,j} \beta_{ij} s_{ij}(x) + \cdots . \tag{3}$$

The ellipses in Eq. 3 represent interactions among three or higher number of mutations. And in practice, one may choose a cutoff number $m \leq \ell$, so that Eq. 3 contains only interactions among up to $m$ mutations.

Mathematically, for any given $m$, by allowing the coefficients $\beta$ to assume arbitrary values, we can define a vector space of genotype-phenotype maps that form a

the expression in Eq. 3 defines a vector space, namely $Y_m$, of genotype-phenotype maps with coefficients $\beta$ assuming arbitrary values.

Mathematically, for any given $m$, the expression in Eq. 3 the set of genotype-phenotype maps defined by allowing the coefficients $\beta$ to assume arbitrary values forms a vector space, which we call $Y_m$. Importantly, changing the choice of wild-type genotype or choosing a one-hot encoding for the genotype-phenotype map (i.e. using indicator functions for alleles and combinations of up to $m$ alleles instead of mutations and combinations of up to $m$ mutations) will still result in this same space of possible genotype-phenotype maps . Thus we can consider $Y_m$ as capturing the expressivity of $m$-th order interaction models independent of any specific parametrization.

Now given an $m$-th order model, we can increase its expressivity by adding order $m+1$ interaction terms, similar to how we extend the additive model to pairwise epistasis model in Eq. 2. Adding these terms results in a larger class of models that corresponds to a bigger vector space $Y_{m+1}$ that contains $Y_m$ as a subspace.

For this paper, we are particularly interested in the extra piece of $Y_{m+1}$ that confers the additional flexibility compared with the space of $m$-th order models, $Y_m$. That is, we want to understand how including interactions between $m+1$ mutations increases the space of possible genotype-phenotype maps beyond what can be expressed using only $m$-th order interactions. Specifically, given a function $f \in Y_{m+1}$, we can regress out its component in the subspace $Y_m$, i.e. we can express $f$ as

$$f = f_m + \phi_{m+1} \tag{4}$$

where $f_m$ is an $m$th-order model (i.e. in $Y_m$) and the residual vector $\phi_{m+1}$ is in $Y_{m+1}$ and is orthogonal to the subspace $Y_m$. This residual $\phi_{m+1}$ represents the part of the model that cannot be captured by epistatic interactions of order up to $m$. Thus, while the function $f$ may contain $(m+1)$-th order interactions as well as lower order interactions, $\phi_{m+1}$ corresponds to a "pure" $(m+1)$-th order interaction model, as it contains no component that can be expressed by a model with fewer than $(m+1)$-th order interactions.

The set of all such pure $m+1$-th order functions also form a vector space, which we write as $V_{m+1}$. Moreover, any function $f$ may be uniquely decomposed as a linear combination:

$$f = \phi_0 + \phi_1 + \cdots + \phi_\ell, \tag{5}$$

where $\phi_k \in V_k$ is the pure $k$-th order component. Importantly, these $\phi_k$ will not typically correspond directly to the contributions of the $k$-th order terms in any given parametrization such as Eq. 3. For example, an epistatic interaction that has a strong positive effect when a particular combination of $k$ mutations co-occur can be approximated in part by an additive model that assigns a positive effect to each of these $k$ mutations. Instead, $\phi_k$ consists of only that component of $f$ that cannot be approximated by any model of order less than $k$.

##### The geometry of $k$-th order interaction models

The first main result of this document states that all models of pure $k$-th order interactions have an identical geometry in terms of how the average phenotype changes as we consider sequences farther and farther away from any given sequence of interest. Given a function $f \in V_k$, for any genotype $x \in A^\ell$, we can calculate the mean value of sequences that are $d$ mutations away from $x$, where we can set $d = 0, 1, \cdots, \ell$. The significance of the mean value function is that it allows us to study the geometry of a genotype-phenotype map centered at a focal genotype using a one dimensional curve, thus greatly reducing the complexity induced by the high dimensionality of the sequence space.

For example, we can express the indicator function of the wild type allele as 1 minus the sum of the indicator functions for the $\alpha - 1$ mutations on the same site. Similar procedure can be done for indicator functions encoding interactions among mutations. This means that the $\alpha^k$ indicator functions for any combination of $k$ sites are not linearly independent, so adding the indicator functions involving wild type alleles or switching the wild type genotype in Eq. 3 does not change its linear span.

We first establish that for a model of order $k$ (i.e., a model in $Y_k$) its distance mean function is a $k$th degree polynomial in $d$. First, consider a model of the form

$$f(x) = \sum_{i_1,\cdots,i_k} \beta_{i_1,\cdots,i_k} s_{i_1,\cdots,i_k}(x), \tag{6}$$

where $s_{i_1,\cdots,i_k}(x)$ is an indicator function encoding the presence of combinations of mutations relative to the WT. Thus for a sequence $x$ that is $d$ mutations away from the WT, we can express its value as

$$f(x) = \overline{\beta}_{i_1,\cdots,i_k}(x)\binom{d}{k}, \tag{7}$$

where $\overline{\beta}_{i_1,\cdots,i_k}(x)$ denotes the mean of the coefficients for allelic combinations present in $x$. Thus, the mean value of sequences that are $d$ mutations away from the WT is simply

$$\mu_{f,\mathrm{WT}}(d) = \langle f(x)\rangle_{D(x,WT)=d} = \overline{\beta}_{i_1,\cdots,i_k}\binom{d}{k}, \tag{8}$$

where $\mu_{f,\mathrm{WT}}(d)$ denotes the mean at distance $d$ for the function $f$ centered at the WT, $\overline{\beta}_{i_1,\cdots,i_k}$ is the mean of all coefficients in Eq. 6 and $\langle\cdot\rangle_{D(x,WT)=d}$ denotes the mean taken over all sequences that are $d$ mutations away from the WT. This is true because every coefficient occurs in the same number of sequences that are $d$ mutations away from the WT. However, the binomial coefficient $\binom{d}{k}$ can be written as a $k$-th order polynomial in $d$, which shows that the distance mean function is also a $k$-th order polynomial in $d$.

So far we have shown that the distance mean function is a polynomial of order $k$ when we consider a model where only combinations of $k$ mutations have an effect (Eq. 6). But a general model of the form of Eq. 3 with up to $k$-th order terms is a sum of such models, where we have terms of orders $0, \ldots, k$, and so the mean function is a sum of polynomials of degree $0, \ldots, k$, and hence itself a polynomial of degree $k$. Therefore, we have established that the distance mean function of any model in $Y_k$ is a $k$th-degree polynomial in $d$.

From a more biological perspective, the above result highlights the limited expressivity of low-order interaction models. For example, the mean value function for an additive model is a straight line, and the mean function of a pairwise model is quadratic regardless of the particular model and choice of WT. Because we would like to fit models that are flexible enough to have any shape of mean value function (e.g. a sigmoid function), it is therefore critical to use methods that allow all orders of interaction.

Interestingly, an even stronger result holds for pure interactions of order $k$ in that not only is the distance function for a pure $k$-th order function a polynomial of order $k$, but for fixed $A$ and $\ell$ it is always the same polynomial up to a multiplicative constant given by the phenotype at the WT. In particular, in Section 2 we show that the shape of the mean value function is the same for all genotypes $x$ in $A^\ell$ and pure $k$-th order function $\phi \in V_k$ up to a multiplicative constant:

$$\mu_{\phi,x}(d) = \phi(x)\mu_k(d), \tag{9}$$

where we use $\mu_{\phi,x}(d)$ to denote the mean at distance $d$ for the function $\phi$ centered at sequence $x$, $\mu_k(d) = \frac{\alpha^\ell}{m_k} w_k(d)$, and $m_k = \binom{\ell}{k}(\alpha-1)^k$ is a constant . Specifically,

$$w_k(d) = \alpha^{-\ell}\sum_{q=0}^{k}(-1)^q(\alpha-1)^{k-q}\binom{d}{q}\binom{\ell-d}{k-q} \tag{10}$$

is known as the Krawtchouk polynomial [4–6] and appears many times throughout the development of our theory, and it can be shown that $w_k(d)$ is a polynomial in $d$ of degree $k$.

Note that given a $k$-th order function $\phi(x)$, the mean function can be calculated with any sequence chosen as the WT. And two mean functions centered at different sequences will only differ by a multiplicative constant, according to Eq. 9.

##### Distance covariance of phenotypes, mutational effects, and local interactions

We now move from the sequence-centric view of the mean value function $\mu$ to the global structure of $k$-th order interaction models. In particular, we focus on the covariance structure in phenotypic values, local mutational effects and local epistatic interactions. The distance covariance function of these quantities dictate how far an observed local effect can be generalized when it occurs in increasingly distant backgrounds. Intuitively, accurately capturing this decay is important for making phenotypic predictions. Our main contribution here is to develop general formulas for the distance covariance functions for mutational effects, and local epistatic interactions of any order, and to relate them to the type and extent of epistasis in the genotype-phenotype map.

Specifically, let $C_f(d)$ be the covariance in phenotypes between sequences that are $d$ mutations away for a genotype-phenotype map $f$, so that [7]

$$C_f(d) = \langle (f(x) - \overline{f}) \left( f(x') - \overline{f} \right) \rangle_{D(x,x')=d}. \tag{11}$$

Then based on the mean function in Eq. 9, we find for a $k$-th order ($k > 0$) model $\phi \in V_k$

$$C_\phi(d) = \langle \phi(x)\phi(x') \rangle_{D(x,x')=d} = \frac{\|\phi\|^2}{\alpha^\ell} \mu_k(d). \tag{12}$$

Note here we calculate the product $\phi(x)\phi(x')$ without explicitly taking out the mean. This is because $\phi$ is orthogonal to the constant subspace $V_0$ and so $\phi$ already has a mean of zero. Also note that this function is constant for any $\phi \in V_k$ up to a multiplicative constant given by the squared norm of $\phi$, which is itself simply the variance of $\phi$ since $\phi$ has a mean of zero.

According to Eq. 5, we can express any genotype-phenotype map $f$ as a sum of its components, $\phi_k$ of pure interactions of different orders $k = 0, 1, \cdots \ell$. And it turns out that the distance covariance of $f$, $C_f(d) = \langle (f(x) - \overline{f})(f(x') - \overline{f}) \rangle_{D(x,x')=d}$ is simply the sum of the covariance functions $C_{\phi_k}(d)$ for each component $\phi_k$ [7]:

$$C_f(d) = \sum_{k=1}^{\ell} C_{\phi_k}(d) = \sum_{k=1}^{\ell} \frac{\|\phi_k\|^2}{\alpha^\ell} \mu_k(d) = \sum_{k=1}^{\ell} \frac{\|\phi_k\|^2}{m_k} w_k(d), \tag{13}$$

where we have used $\mu_k(d) = \frac{\alpha^\ell}{m_k} w_k(d)$. This equation allows us to reduce the empirical distance covariance in $f$ to a simple linear combination of $\ell$ fixed curves $w_k(d)$. Moreover, if we expand the component $\phi_k$ with respect to any orthonormal basis of $V_k$, then the quantity

$$\lambda_k = \frac{\|\phi_k\|^2}{m_k} \quad k = 1, \cdots, \ell \tag{14}$$

can be shown to be equal to the mean squared value of the corresponding regression coefficients (see Section 4). So here we have established a relation between the distance covariance function $C_f(d)$ to the type and extent of epistasis in $f$. Furthermore, since $\phi_k$ is the orthogonal projection of $f$ onto $V_k$, the space of pure $k$-th order interactions, the quantity

$$v_k = \frac{\|\phi_k\|^2}{\|f - \overline{f}\|^2} \quad k = 1, \cdots, \ell, \tag{15}$$

corresponds to the fraction of variance in the model $f$ due to interaction order $k$. So the list $v_k$, $k = 1, \cdots, \ell$ are known as the variance components or amplitude spectra of the genotype-phenotype map $f$ [7–9]. And we can see that one can transform between $\lambda_k$ and $v_k$ using Eqs 14 and 15.

As a result, knowing the mean squared regression coefficients $\lambda_k$, or equivalently the variance components $v_k$ together with the variance of the full landscape, allows us to write down the autocovariance function $C_f(d)$. Conversely, Eq. 13 also allows us to solve for the $\lambda_k$ for $k > 0$ if we are given the autocovariance function.

If we normalize Eq. 13 by the variance in $f$, we get the distance correlation function

$$\rho_f(d) = \sum_{k=1}^{\ell} \frac{v_k}{m_k} w_k(d) = \sum_{k=1}^{\ell} v_k \rho_k(d), \tag{16}$$

where $\rho_k(d) = w_k(d)/m_k$ is the distance correlation of a pure $k$-th order function. This allows us to draw Figure 2 in the main text.

Using a similar argument, we can generalize the phenotypic covariance $C(d)$ to the covariance in mutational effects [10]. This allows us to assess the average predictability of the effect of a mutation in a genetic background that is a given number of mutations away. One of the main contributions of this paper is the generalization of the covariance in phenotypes and mutational effects to the distance covariance in local pairwise, three-way, and higher-order interactions.

Furthermore, it turns out that the distance covariance $\Gamma_k(d)$ of mutational effects ($k = 1$) and local interactions ($k \geq 2$) is completely determined by the phenotypic covariance function $C(d)$

$$\Gamma_k(d) = 2^k \sum_{q=0}^{k} (-1)^q \binom{k}{q} (C(d+q) + \overline{f}^2). \tag{17}$$

Importantly, we also find a strong parallelism between $\Gamma_k(d)$ and $C(d)$ in their relation to the mean squared regression coefficients $\lambda_k$ (see Section 8):

$$\Gamma_k(d) = 2^k \sum_{j=0}^{\ell-k} \lambda_{k+j} w_j^{\ell-k}(d), \tag{18}$$

where $w_j^{\ell-k}$ is the Krawtchouk polynomial of order $j$ for a sequence space of reduced length $\ell - k$.

We can also normalize $\Gamma_k(d)$ by dividing it with $\Gamma_k(0)$, i.e. the variance in $k$-th order interaction effects, to derive the distance autocorrelation [11]:

$$\gamma_k(d) = \frac{\Gamma_k(d)}{\Gamma_k(0)} = \sum_{j=0}^{\ell-k} v_{k+j}^{\ell-k} \rho_j^{\ell-k}(d). \tag{19}$$

Here $v_{k+j}^{\ell-k}$ is the renormalized variance component

$$v_{k+j}^{\ell-k} = m_j^{\ell-k} \lambda_{k+j} / \left( \sum_{i=1}^{\ell-k} m_i^{l-k} \lambda_{i+j} \right) \tag{20}$$

where $m_j^{\ell-k} = \binom{\ell-k}{j} (\alpha - 1)^j$ is the dimensionality of the vector space of pure $j$-th order interactions for a sequence of length $\ell - k$, and $\rho_j^{\ell-k}(d)$ is the distance autocorrelation in $j$-th order interaction in a sequence space of length $\ell - k$. This allows us to make Figure 1. Notice that to calculate the distance correlation function of local $k$-way interactions, we simply drop $\lambda_\ell$, $l < k$, and construct $\gamma_k$ as one would for a smaller sequence space of length $\ell - k$, using the corresponding order-specific distance correlation functions, $\rho_j^{\ell-k}, j = k, \cdots, \ell$.

**Bayesian inference**

We now proceed in a Empirical Bayes framework by using the information that we extract from the data to build an informative prior for inference. Specifically, we want to build a prior distribution over

$X$, the space of all genotype-phenotype maps. We further specify that our prior should be Gaussian and isotropic, that is, the covariance between sequences is a function of their Hamming distance, and we set the mean of the prior to be the constant function with zeros for all sequences. So the essential part of the prior is to choose the covariance, or the kernel function $K$ that specifies the similarity between sequences as a function of their Hamming distance. Importantly, because we are assuming that the prior is isotropic, the covariance in our prior takes on only $\ell+1$ distinct values: $K(d), d = 0, 1, \cdots, \ell$ (typically the posterior will be highly anisotropic, because its covariance structure will depend on which specific sequences were observed). In Section 4, we show that under the isotropic condition, all possible kernels describing such a prior belong to a family parametrized by the variance in regression coefficients $\lambda_k$, $0 \le k \le \ell$:

$$K(d) = \sum_{k=0}^{\ell} \lambda_k w_k(d). \tag{21}$$

Note that this function is very similar to Eq. 13. The difference is that Eq. 13 concerns the covariance between sequences in a realized genotype-phenotype map $f$, while the equation above specifies the covariance in a distribution over all possible genotype-phenotype maps that we use as a prior for Bayesian inference. In particular, here the $\lambda_k$ are hyperparameters that control the average magnitude of regression coefficients for different orders of genetic interaction. To achieve good prediction, we would like to choose an informative prior that reflect some important large scale features of the data. To do this, we first note that Eq. 13 allows us to estimate the $\lambda_k$'s and the variance components from the distance covariance function $C(d)$, even when the data represents a partial sampling of $f$, since we can extract the empirical distance covariance function $\widehat{C}(d)$ by enumerating through all pairs of sequences in the data

$$\widehat{C}(d) = \langle (f(x) - \overline{f})(f(x') - \overline{f}) \rangle_{D(x,x')=d \text{ and } x,x' \text{ in the data}}, \tag{22}$$

from which we can get an estimate for the variance in regression coefficients for all orders $k$, $\widehat{\lambda}_k$ (or equivalently the variance components) by fitting $\widehat{C}(d)$ using the parametric form $\sum_{k=0}^{\ell} \widehat{\lambda}_k w_k(d)$ under the constraint $\widehat{\lambda}_k > 0$ for all $k$ (see Section 10 for detail of the inference procedure).

To encode the information about epistasis in the data into our prior, we simply set our kernel to

$$K(d) = \sum_{k=0}^{\ell} \widehat{\lambda}_k w_k(d). \tag{23}$$

So a typical sample from the prior under this kernel function will have similar distance covariance as the data $\widehat{C}(d)$. And based on Eqs. 13 and 17, the sample will also have similar variance components and covariance in mutational effect and local epistatic interaction as the data.

For inference, we use a Gaussian process regression framework [12], with prior distribution

$$\mathbf{f} \sim \mathcal{N}(\mu, \mathbf{K}). \tag{24}$$

Here the covariance matrix $\mathbf{K}$ contains pairwise covariance between all possible pairs of sequences and we can use our kernel function $K$ to populate $\mathbf{K}$ so that $\mathbf{K}_{x,x'} = K(D(x, x'))$. Furthermore, Eq. 23 allows us to expand $\mathbf{K}$ as a linear combination

$$\mathbf{K} = \sum_{k=0}^{\ell} \widehat{\lambda}_k \mathbf{W}_k, \tag{25}$$

where $\mathbf{W}_k$ is determined by the Krawtchouk polynomial $w_k(d)$.

The joint distribution of a full-length genotype-phenotype map $\mathbf{f} \in \mathbb{R}^{\alpha^\ell}$, and the data $\mathbf{y} \in \mathbb{R}^m$ is [12]

$$\begin{bmatrix} \mathbf{f} \\ \mathbf{y} \end{bmatrix} \sim \mathcal{N}(\begin{bmatrix} \mathbf{0}_{\alpha^\ell} \\ \mathbf{0}_m \end{bmatrix}, \begin{bmatrix} \mathbf{K} & \mathbf{K}_{\cdot B} \\ \mathbf{K}_{\cdot B} & \mathbf{K}_{BB} + \mathbf{E} \end{bmatrix}), \tag{26}$$

where $B$ denotes the subset of all sequences in $A^\ell$ that is in the data and $\mathbf{E}$ is a diagonal covariance matrix whose diagonal entries are set to estimates of the experimental noise variance for each sequence, corresponding to independent experimental noise for each observed sequence. Specifically, the noise distribution is also set to be Gaussian with $\mathbf{e} \sim \mathcal{N}(\mathbf{0}, \mathbf{E})$, and $\mathbf{y} = \mathbf{f}_B + \mathbf{e}$. From this we can calculate the posterior distribution for $\mathbf{f}$ [12]

$$\mathbf{f}|\mathbf{y} \sim \mathcal{N}(\mathbf{K}_{\cdot B}(\mathbf{K}_{BB} + \mathbf{E})^{-1}\mathbf{y}, \mathbf{K} - \mathbf{K}_{\cdot B}(\mathbf{K}_{BB} + \mathbf{E})^{-1}\mathbf{K}_{B\cdot}). \tag{27}$$

In Section 9, we make the connection between this Bayesian inference procedure and regularized regression. And in Section 11, we introduce a method for efficiently sampling from the posterior in Eq. 27.

### 2 Distance mean function of pure $k$th-order interactions

Our aim for this section is to derive the distance mean function for any genotype-phenotype map consisting of pure $k$-th order interactions. To do this, we need some basic facts in spectral fitness landscape theory. Throughout this section, we will use known facts about the graph structure of the sequence space without proof. For a thorough introduction to spectral fitness landscape theory, see refs. [5, 9].

Briefly, the sequence space $A^\ell$ and the space of functions defined over $A^\ell$ can be more easily studied by introducing a graph structure over $A^\ell$. Specifically, define a graph with all the sequences in $A^\ell$ as vertices, and two sequences $x$ and $x'$ are adjacent on this graph if and only if they differ by a point mutation, i.e. $D(x, x') = 1$. The structure of this so-called Hamming graph can be represented by a $\alpha^\ell \times \alpha^\ell$ matrix known as the graph Laplacian with entries indexed by sequences in $A^\ell$:

$$\mathbf{L}_{x,x'} = \begin{cases} (\alpha-1)\ell & D(x,x') = 0 \\ -1 & D(x,x') = 1 \\ 0 & \text{otherwise,} \end{cases} \tag{28}$$

where $(\alpha - 1)\ell$ is the degree (number of neighbors) of a vertex which is equal to the number of distance 1 neighbors.

We are interested in the eigenvalues and eigenvectors of $\mathbf{L}$, since they are directly related to different orders of genetic interactions. To derive the eigenvalues of $\mathbf{L}$ for the Hamming graph of sequence space of arbitrary length, we start out with the simplest Hamming graph with $\ell = 1$. In this case, a sequence reduces to a letter in the alphabet and is connected to every other sequence. In other words, for $\ell = 1$, the Hamming graph coincides with the complete graph. It is well known that the Laplacian of a complete graph with $\alpha$ vertices has eigenvalues 0 and $\alpha$, with multiplicity 1 and $\alpha - 1$, respectively. To derive the eigenvalues of $\mathbf{L}$ for general sequence spaces of length $\ell \geq 2$, we note that the Hamming graph for sequence space of length $\ell$ can be constructed as an $\ell$-fold Cartesian product of the complete graph. A direct consequence of this construction is that the graph Laplacian $\mathbf{L}$ for $\ell \geq 2$ has only $\ell + 1$ distinct eigenvalues $\lambda_k = \alpha k$, $k = 0, \ldots, \ell$ with multiplicity $m_k = \binom{\ell}{k}(\alpha - 1)^k$ [5, 13]. Now we can directly relate the eigensystem of $\mathbf{L}$ to different orders of genetic interactions. Specifically, we find that $V_k$, i.e. the space of pure $k$-th order interactions constructed in Section 1, coincides with the eigenspace of $\mathbf{L}$ associated with eigenvalue $\lambda_k$ [14]:

$$V_k = \{\mathbf{v} : \mathbf{L}\mathbf{v} = \lambda_k \mathbf{v}\}. \tag{29}$$

Since the multiplicity of eigenvalue $\lambda_k$ is $m_k$, we have $\dim(V_k) = m_k$.

Now that we have established some basic facts about the eigensystem of the graph Laplacian $\mathbf{L}$ and its connection to $k$-th order interactions, we can turn to the distance mean function of $k$-th order interactions. Let $\phi$ be a genotype-phenotype map with pure $k$-th order interaction. Thus, $\phi \in V_k$, and $\mathbf{L}\boldsymbol{\phi} = \lambda_k \boldsymbol{\phi}$.

Here we choose to work with the adjacency matrix $\mathbf{A} = \mathbf{D} - \mathbf{L}$, where $\mathbf{D} = (\alpha - 1)\ell$ is a diagonal matrix. It is easy to verify that

$$\mathbf{A}\boldsymbol{\phi} = ((\alpha - 1)\ell - \lambda_k)\boldsymbol{\phi} \equiv \eta_k \boldsymbol{\phi}. \tag{30}$$

Given a focal genotype $x$, we can divide the sequence space $A^\ell$ into disjoint sets of sequences that are certain distances away from $x$, $P_d^x = \{x' \in A^\ell : D(x, x') = d\}$, so that,

$$A^\ell = \cup_{d=0}^{\ell} P_d^x. \tag{31}$$

It is easy to verify that $|P_d^x| = m_d = \binom{\ell}{d}(\alpha - 1)^\ell$. Since the value of $\phi(x')$ can be expressed as a sum of the values of genotypes adjacent to $x'$, that is, $\eta_k \phi(x') = \sum_{y \sim x'} \phi(y)$ according to Eq. 30, by summing through all sequences $x'$ in $P_d^x$, we find

$$\eta_k \sum_{x' \in P_d^x} \phi(x') = \sum_{x' \in P_d^x} \sum_{y \sim x'} \phi(y) \tag{32}$$

$$= \sum_{d'} \sum_{y \in P_{d'}^x} \left( \sum_{x' \in P_d^x, x' \sim y} A_{x'y} \right) \phi(y) \tag{33}$$

$$= \sum_{d'} \sum_{y \in P_{d'}^x} \widehat{A}_{d,d'} \phi(y) \tag{34}$$

$$= \sum_{d'} \widehat{A}_{d,d'} \sum_{y \in P_{d'}^x} \phi(y). \tag{35}$$

Here we define $\widehat{A}_{d,d'} \equiv \sum_{x' \in P_d^x, x' \sim y} A_{x'y}$, which is equal to the number of sequences in the distance class $P_d^x$ that are adjacent to a sequence $y$ in the distance class $P_{d'}^x$. It is easy to verify that this number does not change with respect to the choice of $x$ and $y$, and thus only depends on $d$ and $d'$.

Furthermore, we can arrange the values of $\widehat{A}_{d,d'}$ for $0 \le d, d' \le \ell$ on an $(\ell + 1) \times (\ell + 1)$ matrix with row and columns indexed by $d$ and $d'$. This matrix $\widehat{\mathbf{A}}$ is known as the collapsed adjacency matrix of the Hamming graph [9]. We then find the sum at distance $d$,

$$\sigma_{\phi,x}(d) \equiv \sum_{y \in P_d^x} \phi(y), \tag{36}$$

for $0 \le d \le \ell$ is a right eigenvector of $\widehat{\mathbf{A}}$ with eigenvalue $\eta_k$. We can further verify that $\widehat{\mathbf{A}}$ has the same distinct eigenvalues as $\mathbf{A}$. Moreover, the right eigenvector associated with eigenvalue $\eta_k$ is given by [5, 7]

$$\zeta_k(d) = \alpha^\ell \frac{m_d}{m_k} w_k(d) \quad 0 \le d \le \ell, \tag{37}$$

where $w_k(d)$ is the Krawtchouk polynomial. Since, $\zeta_k$ and $\sigma_{\phi,x}$ are both eigenvectors of $\widehat{\mathbf{A}}$ with eigenvalue $\eta_k$, and $\eta_k$ has multiplicity 1, we find $\zeta_k(d)$ and $\sigma_{\phi,x}(d)$ differ by at most a multiplicative constant. And since $\zeta_k(0) = 1$ according to the definition of the $w_k(d)$, and $\sigma_{\phi,x}(0) = \phi(x)$, we find

$$\sigma_{\phi,x}(d) = \phi(x)\zeta_k(d). \tag{38}$$

Lastly, to calculate the mean value for sequences in $P_d^x$, we simply divide Eq. 38 by $|P_d^x| = m_d$,

$$\mu_{\phi,x}(d) = \frac{1}{m_d}\phi(x)\zeta_k(d) = \phi(x)\frac{\alpha^\ell}{m_k} w_k(d) \equiv \phi(x)\mu_k(d). \tag{39}$$

This gives us Eq. 9 in the Introduction.

#### 3 Summary statistics for empirical genotype-phenotype maps

This section defines various summary statistics for an empirical genotype-phenotype map used in the main text, including the quantities plotted in Figure 1 in the main text. We also show how different quantities can be transformed from one to another. Importantly, we elucidate how the distance covariance function in phenotype is connected to the mean squared regression coefficients of different orders of interactions.

Recall the distance covariance function of a genotype-phenotype map $f$ defined in the Introduction.

$$C_f(d) = \langle (f(x) - \overline{f})(f(x') - \overline{f}) \rangle_{D(x,x')=d} = \frac{1}{N_d} \sum_{x,x':D(x,x')=d} (f(x) - \overline{f})(f(x') - \overline{f}), \tag{40}$$

where $N_d = \alpha^\ell \binom{\ell}{d}(\alpha - 1)^d$ is the number of pairs of sequences at Hamming distance $d$ and $\overline{f}$ is the mean phenotypic value. We can also define the autocorrelation function by normalizing $C(d)$ with the empirical variance:

$$\rho_f(d) = \frac{C_f(d)}{C_f(0)}. \tag{41}$$

Also recall that we have the analytical formula for the distance covariance function for a pure $k$-th order interaction model $\phi_k$:

$$C_{\phi_k}(d) = \langle \phi_k(x)\phi_k(x') \rangle_{D(x,x')=d} = \frac{1}{\alpha^\ell} \sum_x \phi_k(x) \left( \frac{1}{m_d} \sum_{x':D(x,x')=d} \phi_k(x') \right) = \frac{\|\boldsymbol{\phi}_k\|^2}{\alpha^\ell} \mu_k(d), \tag{42}$$

Furthermore, for $f = \sum_{k=0} f_k$, we can calculate its distance covariance similarly:

$$\begin{aligned} C_f(d) &= \frac{1}{\alpha^\ell} \sum_x (f(x) - \overline{f}) \sum_{k=1}^{\ell} \phi_k(x)\mu_k(d) && (43) \\ &= \frac{1}{\alpha^\ell} \sum_{k=1}^{\ell} \left( \sum_x (f(x) - \overline{f})\phi_k(x) \right) \mu_k(d) && (44) \\ &= \frac{1}{\alpha^\ell} \sum_{k=1}^{\ell} \|\boldsymbol{\phi}_k\|^2 \mu_k(d) && (45) \\ &= \sum_{k=1}^{\ell} \frac{\|\boldsymbol{\phi}_k\|^2}{m_k} w_k(d). && (46) \end{aligned}$$

Here we have used the fact that $\sum_x (f(x) - \overline{f})\phi_k(x) = (\mathbf{f} - \overline{f}\mathbf{1})^T \boldsymbol{\phi}_k = \|\boldsymbol{\phi}_k\|^2$, for $k \geq 1$.

We now show that the quantity $\lambda_k = \frac{\|\boldsymbol{\phi}_k\|^2}{m_k}$ is the mean squared regression coefficient for $k$-th order interactions by expanding $\phi_k$ with respect to an orthonormal basis. Recall $\phi_k$ is the projection of $f$ onto the eigenspace $V_k$. We can now define an arbitrary orthonormal basis $\{u_{k,i}\}_{1 \leq i \leq m_k}$ of $V_k$. There are $m_k$ vectors in the basis since the dimension of $V_k$ is $m_k$. This allows us to express $\phi_k = \sum_{i=1}^{m_k} a_{k,i} u_{k,i}$. Therefore, the quantities $a_{k,i}$ are the regression coefficients of $\phi_k$ under this particular choice of basis. Given that the vectors $\mathbf{u}_{k,i}$ are orthonormal, we find

$$\lambda_k = \frac{\|\boldsymbol{\phi}_k\|^2}{m_k} = \frac{\| \sum_{i=1}^{m_k} a_{k,i} \mathbf{u}_{k,i} \|^2}{m_k} = \frac{\sum_{i=1}^{m_k} a_{k,i}^2}{m_k}, \tag{47}$$

which is the mean squared regression coefficient of order $k$.

In the special case when $k = 0$, $\lambda_0 = \frac{\|\phi_0\|^2}{m_0} = \|\phi_0\|^2$, since $m_0$ is equal to 1. Here $\phi_0$ is the projection onto $V_0$, the constant subspace, which is spanned by the unit vector $\mathbf{u} = \alpha^{-\frac{\ell}{2}}\mathbf{1}$, where $\mathbf{1}$ is the vector of all ones. Therefore, we find

$$\lambda_0 = \|\phi_0\|^2 = \|(\mathbf{f}^T\mathbf{u})\mathbf{u}\|^2 = (\mathbf{f}^T\mathbf{u})^2 = (\alpha^{-\frac{\ell}{2}}\sum_x f(x))^2 = \alpha^\ell \overline{f}^2. \tag{48}$$

Lastly, suppose we only have noisy observations $\mathbf{y} = \mathbf{f}_B + \mathbf{e}$ on a subset of sequences $B \subset A^\ell$. Here $\mathbf{e}$ is the noise vector which we assume is drawn from a normal distribution: $\mathbf{e} \sim \mathcal{N}(\mathbf{0}, \mathbf{E})$, with $\mathbf{E}$ being a diagonal matrix. We can extract the empirical covariance function by averaging over pairs of sequences in $B$ for different distance classes. Specifically, let $\mathcal{D}(B) = (\mathcal{D}_0, \mathcal{D}_1, \ldots, \mathcal{D}_\ell)$ be the distance distribution of the set $B$, where $\mathcal{D}_i$ is the number of pairs of ordered sequences that are at Hamming distance $i$. Define the empirical autocovariance function

$$c(d) = \begin{cases} \frac{1}{|B|}\sum_{x\in B}(y(x) - \overline{y})^2 - \overline{\sigma^2} & d = 0 \\ \frac{1}{\mathcal{D}_d}\sum_{\{x,x'\in B: D(x,x')=d\}}(y(x) - \overline{y})(y(x') - \overline{y}) & d = 1, \cdots, \ell. \end{cases} \tag{49}$$

For $d = 0$, we substract the mean variance of the noise components $\overline{\sigma^2}$ from the raw empirical variance $\frac{1}{|B|}\sum_{x\in B}(y(x) - \overline{y})^2$ so that $c(0)$ is not inflated by the observation noise. The noise component is not accounted for when $d > 0$ since we assume the noise distribution is independent with mean zero, making the contribution from noise to $c(d)$ for $d > 0$ negligible. Note that $C(d)$ and $c(d)$ coincide if we have data for all sequences and the data is noise-free.

### 4 Gaussian random field landscapes

So far we have focused on the structure of an empirical landscape $\mathbf{f}$ . In this section, we extend our theory to a probabilistic framework that allows us to perform Bayesian inference for genotype-phenotype maps. More specifically, we will first define a covariance structure for a Gaussian prior. We then describe our inference procedure using a Gaussian process regression framework.

Given noisy observations on a subset of all possible sequences, our aim is to reconstruct the full landscape $\mathbf{f}$ so that the reconstructed landscape reflects the statistical features of the observed data. Since the underlying landscape $\mathbf{f}$ is unknown, we use a Bayesian strategy by treating it as a random function that is drawn from a Gaussian prior, that is

$$\mathbf{f} \sim \mathcal{N}(\boldsymbol{\mu}, \mathbf{K}), \tag{50}$$

where $\boldsymbol{\mu} \in \mathbb{R}^{\alpha^\ell \times 1}$ and $\mathbf{K} \in \mathbb{R}^{\alpha^\ell \times \alpha^\ell}$ are the mean vector and covariance matrix, respectively. Throughout this paper, we assume the prior distribution has mean zero, i.e. $\boldsymbol{\mu} = \mathbf{0}$.

In section 1, we pick the matrix $\mathbf{K}$ by assuming that the distribution is isotropic and stated that all isotropic matrices can be populated by a kernel function $K(d) = \sum_{k=0}^{\ell}\lambda_k w_k(d)$. Here we provide a constructive proof for this result, using arbitrary orthonormal bases for the subspaces of different orders of genetic interaction $V_k$ and independent, Gaussian-distributed regression coefficients with respect to these bases.

We start out by defining simple distributions for the regression coefficients of different orders $k$. Specifically, we assume the regression coefficients are independent and Gaussian with mean 0 and identical

Throughout this paper, we use the boldface $\mathbf{f} \in \mathbb{R}^{\alpha^\ell}$ to denote the $\alpha^\ell$-dimensional column vector indexed by sequences in $A^\ell$ and $f$ to denote the function which allows us to evaluate the phenotype of a sequence $f(x)$ such that $f(x)$ is the $x$-th entry of $\mathbf{f}$.

variance within each order $k$. Let $\mathbf{a}_k \in \mathbb{R}^{m_k}$ be the random vector containing all $k$-th order regression coefficients, then

$$\mathbf{a}_k \sim \mathcal{N}(\mathbf{0}, \lambda_k \mathbf{I}_{m_k}). \tag{51}$$

Let $\mathbf{Q}_k$ be the matrix with the orthonormal vectors $\{\mathbf{u}_{k,i}\}_{1\leq i\leq m_k}$ as columns. It is easy to check that the random vector $\mathbf{f} = \sum_{k=0}^{\ell} \mathbf{Q}_k \mathbf{a}_k$ is also Gaussian and has mean 0. Furthermore, its covariance matrix is

$$\mathbf{K} = \mathbb{E}_{\mathbf{f}}\left[\mathbf{f}\mathbf{f}^T\right] = \mathbb{E}_{\mathbf{a}_0,\mathbf{a}_1,\cdots,\mathbf{a}_\ell}\left[\sum_{j=0}^{\ell} \mathbf{Q}_j \mathbf{a}_j \left(\sum_{k=0}^{\ell} \mathbf{Q}_k \mathbf{a}_k\right)^T\right] = \sum_{j,k} \mathbf{Q}_j \mathbb{E}_{\mathbf{a}_j,\mathbf{a}_k}\left[\mathbf{a}_j \mathbf{a}_k^T\right] \mathbf{Q}_k^T \tag{52}$$

$$= \sum_{k=0}^{\ell} \lambda_k \mathbf{Q}_k \mathbf{Q}_k^T = \sum_{k=0}^{\ell} \lambda_k \mathbf{W}_k. \tag{53}$$

Here $\mathbf{W}_k = \mathbf{Q}_k \mathbf{Q}_k^T$ is a $\alpha^\ell \times \alpha^\ell$ matrix whose entries are given by the Krawtchouk polynomial [4, 9], where these entries only depend on the Hamming distance between sequences:

$$\mathbf{W}_k(x, x') = w_k^\ell(d(x, x')). \tag{54}$$

So far we have defined a family of Gaussian prior distributions for $\mathbf{f}$ with covariance matrix $\mathbf{K} = \sum_{k=0}^{\ell} \lambda_k \mathbf{W}_k$, where the $\lambda_k > 0$ serve as hyperparameters of the prior distribution, which can be specified *a priori* or inferred from the data. Furthermore, since the columns of $\mathbf{Q}_k$ are orthonormal, $\mathbf{W}_k = \mathbf{Q}_k \mathbf{Q}_k^T$ is the projection matrix to the space of $k$-th order interactions. As a result, the matrix $\mathbf{K}$ defined above is guaranteed to be positive-definite if $\lambda_k > 0$ for all $\ell \geq k \geq 0$, therefore is a proper covariance matrix.

Because the $\mathbf{W}_k(x, x')$ only depend on the Hamming distance between pairs of sequences, the covariance of this prior distribution likewise is a function of the Hamming distance between sequences. In other words, we have defined a Gaussian isotropic random field [5, 8, 9]. This allows us to summarize the covariance structure of our prior distribution by the following kernel function

$$K(d) = \sum_{k=0}^{\ell} \lambda_k w_k^\ell(d). \tag{55}$$

Note that Eq. 55 is very similar to Eq. 13. The main difference is that here $\lambda_k > 0$ are hyperparameters for the prior that specify the variance of any coefficient of order $k$ in our prior. In contrast, in Eq. 13, $\lambda_k$ is the mean squared $k$-th order regression coefficient of a specific landscape. Note that we also include the 0th order term $\lambda_0$ in Eq. 55 because we would like to include in our prior all functions, instead of just functions that have the same mean as $\boldsymbol{\mu}$. In particular, the mean of a draw from the prior is normally distributed with variance $\alpha^{-\ell}\lambda_0$. As a result, the expected empirical autocovariance function differs from the kernel function by a constant:

$$\mathbb{E}_{\mathbf{f}}\left[C_{\mathbf{f}}(d)\right] = K(d) - \alpha^{-\ell}\lambda_0. \tag{56}$$

We write $I = A^\ell \setminus B$ as the set of all missing sequences. Throughout this paper, we also use $B$ and $I$ to denote columns and rows of matrices that are indexed by $A^\ell$. For example, $\mathbf{K}_{BB}$ is the $m \times m$ submatrix of $\mathbf{K}$ generated by selecting rows and columns corresponding to $B$, while $\mathbf{K}_{\cdot B}$ denotes the $\alpha^\ell \times m$ matrix whose columns correspond to sequences in $B$.

Recall that $\mathbf{y} = \mathbf{f}_B + \mathbf{e} \in \mathbb{R}^m$ is the vector of observations for the subset $B$, where $\mathbf{e}$ is a vector of observation noise so that $\mathbf{e} \sim \mathcal{N}(\mathbf{0}, \mathbf{E})$ with $\mathbf{E}$ being a diagonal matrix. The distribution of $\mathbf{y}$ is

$$\mathbf{y} \sim \mathcal{N}(\mathbf{0}, \mathbf{K}_{BB} + \mathbf{E}). \tag{57}$$

Without loss of generality, we will order our sequences so that the $m$ sequences in $B$ whose phenotypes are known come first. The joint distribution of the full landscape $\mathbf{f}$ and $\mathbf{y}$ is then

$$\begin{bmatrix}\mathbf{f}\\ \mathbf{y}\end{bmatrix} \sim \mathcal{N}(\begin{bmatrix}\mathbf{0}_{\alpha^\ell}\\ \mathbf{0}_m\end{bmatrix}, \begin{bmatrix}\mathbf{K} & \mathbf{K}_{\cdot B}\\ \mathbf{K}_{\cdot B} & \mathbf{K}_{BB}+\mathbf{E}\end{bmatrix}). \tag{58}$$

The posterior distribution for $\mathbf{f}$ is also Gaussian and is given by well-known formula for Gaussian process regression [12]

$$\mathbf{f}|\mathbf{y} \sim \mathcal{N}(\mathbf{K}_{\cdot B}(\mathbf{K}_{BB}+\mathbf{E})^{-1}\mathbf{y}, \mathbf{K}-\mathbf{K}_{\cdot B}(\mathbf{K}_{BB}+\mathbf{E})^{-1}\mathbf{K}_{B\cdot}), \tag{59}$$

where $\mathbf{K}_{\cdot B}(\mathbf{K}_{BB}+\mathbf{E})^{-1}\mathbf{y} = \widehat{\mathbf{f}}$ is know as the maximum a posteriori (MAP) estimate. The posterior variance for a single sequence $x$ can be calculated as

$$\sigma_x^2 = \mathbf{K}_{xx} - \mathbf{K}_{xB}(\mathbf{K}_{BB}+\mathbf{E})^{-1}\mathbf{K}_{Bx}\ , \tag{60}$$

where $\mathbf{K}_{Bx} = \mathbf{K}_{xB}^T \in \mathbb{R}^m$ is the column vector containing the covariance between the genotype $x$ and every genotype in the training data $B$.

### 5 Definition of the epistatic coefficient $\epsilon_C(\mathbf{f})$

In this section, we define a natural generalization of the familiar epistatic coefficient, which is defined between a pair of sites, and our generalization extends its definition to $k \geq 2$ sites. We show that the $k$th-order epistatic coefficient can be viewed as the deviation from the linear prediction based on all lower order epistatic coefficients. We then dedicate the next two sections to study the distance correlation of $k$th-order epistatic coefficients for a random field, and then also for empirical landscapes.

First, given a reference sequence $x$, let $S = \{S_1, \cdots, S_k\} \subset \{1, \cdots, \ell\}$ denote a subset of $k$ sites, so that $x_{S_1}$ returns the allele of $x$ at site $S_1$. Let $\psi \in A \setminus \{x_{S_1}\} \times \cdots \times A \setminus \{x_{S_k}\}$ be a list of length $k$ that specifies a "mutant" allele for each of the positions in $S$. Next, let $x^{s,\psi_s}$ denote the sequence with the mutant alleles prescribed by $\psi$ on a subset $s$ of $S$. Let $C_x^{S,\psi}$ denote the set of all sequences with different combinations of wild type and mutant alleles on $S$, i.e. $C_x^{S,\psi} = \{x^{s,\psi_s} : s \in 2^S\}$, where $2^S$ denotes the power set of $S$. We find that in the case where $k = 1$, $C_x^{S,\psi}$ is an "edge" formed by the "wild type" $x$ and a single mutant. And when $k = 2$, we have four sequences including $x$, the two single mutants, and the double mutant, which form a "square". For $k \geq 3$, $C_x^{S,\psi}$ defines a binary sequence space or, geometrically, a $k$-dimensional hypercube that is embedded in the larger space $A^\ell$.

Note that the set $C_x^{S,\psi}$ can also be built by picking $k$ variable positions, for which we further specify two alleles per positions, and a choice of "background" sequence on the remaining $\ell - k$ positions . Thus, using simple combinatorial arguments, we find the total number of $k$-dimensional cubes one can build is

$$\binom{\ell}{k}\binom{\alpha}{2}^k \alpha^{\ell-k}. \tag{61}$$

Let $\mathbf{f}$ be a genotype-phenotype map. For any hypercube $C_x^{S,\psi}$ of size $2^k$, define the generalized $k$-th order epistatic coefficient associated with the cube $C_x^{S,\psi}$ to be

$$\epsilon_x^{S,\psi}(\mathbf{f}) = (-1)^k \sum_{s\in 2^S} (-1)^{|s|} f(x^{s,\psi_s}) \tag{62}$$

$$= (-1)^k \sum_{x' \in C_x^{S,\psi}} (-1)^{h(x')} f(x'), \tag{63}$$

Note that $C_x^{S,\psi}$ is simply a set and can be built using different choices of "wild type" sequences where $h(x')$ denotes the Hamming distance between $x$ and $x'$. For $k=1$, we call $\epsilon_x^{S,\psi}(\mathbf{f})$ the local additive coefficient (which corresponds to an edge of the Hamming graph), and for $k=2$ we call it a local pairwise epistatic coefficient (which is evaluated on a single square face of the larger sequence space).

Now we justify this definition of the generalized epistatic coefficient as the deviation from the linear predictions made based on lower-order local epistatic interactions. Specifically, in the cube $C_x^{S,\psi}$, consider $y$ the sequence with all mutant alleles on $S$, that is, the sequence in $C_x^{S,\psi}$ that is $k$ mutations away from $x$. Now consider the problem of predicting the values for $y$, given the values for all other sequences in $C_x^{S,\psi}$. We note that $y$ may contain the local additive effects of mutations relative to $x$, which can be calculated by taking the difference between the values of the single mutants and $x$. Furthermore, $y$ may also contain local epistatic coefficients of various orders, which can be calculated using the values of sequences in $C_x^{S,\psi}$. Thus, a reasonable prediction for the value of $y$ is:

$$\sum_{S' \subset S} \epsilon_x^{S',\psi_{S'}}(\mathbf{f}), \tag{64}$$

where $S'$ is a subset of $S$, and $\psi_{S'}$ is the corresponding list of mutant alleles, so that $\epsilon_x^{S',\psi_{S'}}(\mathbf{f})$ defines the local epistatic coefficient on a smaller hypercube anchored at $x$ and contained in $C_x^{S,\psi}$. Thus $\sum_{S' \subset S} \epsilon_x^{S',\psi_{S'}}(\mathbf{f})$ contains the local additive effects, pairwise, and higher-order epistatic interactions. It turns out the local epistatic coefficient is equal to the deviation of the true value of $y$ from this local epistatic prediction:

$$f(y) - \sum_{S' \subset S} \epsilon_x^{S',\psi_{S'}}(\mathbf{f}) = \epsilon_x^{S,\psi}(\mathbf{f}). \tag{65}$$

*Proof.* We begin by converting the enumeration of $S'$ in the sum $\sum_{S' \subset S} \epsilon_x^{S',\psi'_{S'}}(\mathbf{f})$ to the enumeration of $x'$. Using Eq. 63, we find

$$\begin{aligned}
\sum_{S' \subset S} \epsilon_x^{S',\psi_{S'}}(\mathbf{f}) &= \sum_{\{x' \in C_x^{S,\psi} : x' \neq y\}} f(x') \sum_{q=0}^{k-h(x')-1} (-1)^{h(x')+q} (-1)^{h(x')} \binom{k-h(x')}{q} && (66)\\
&= \sum_{\{x' \in C_x^{S,\psi} : x' \neq y\}} f(x') \sum_{q=0}^{k-h(x')-1} (-1)^{q} \binom{k-h(x')}{q} && (67)\\
&= \sum_{\{x' \in C_x^{S,\psi} : x' \neq y\}} (-1)^{k-h(x')+1} f(x') && (68)\\
&= (-1)^{k+1} \sum_{\{x' \in C_x^{S,\psi} : x' \neq y\}} (-1)^{h(x')} f(x'), && (69)
\end{aligned}$$

where we have used the fact that $\sum_{q=0}^{k-h(x')} (-1)^q \binom{k-h(x')}{q} = 0$. Therefore, we find

$$\begin{aligned}
f(y) - \sum_{S' \subset S} \epsilon_x^{S',\psi'_{S'}}(\mathbf{f}) &= (-1)^{2k} f(y) + (-1)^k \sum_{\{x' \in C_x^{S,\psi} : x' \neq y\}} (-1)^{h(x')} f(x') && (70)\\
&= (-1)^k \sum_{\{x' \in C_x^{S,\psi} : x' \neq y\}} (-1)^{h(x)} f(x') && (71)\\
&= \epsilon_x^{S,\psi}(\mathbf{f}). && (72)
\end{aligned}$$

□

In the next two sections, we derive the distance covariance function for $k$-th order epistatic coefficients for random field landscapes, and complete empirical landscapes. The statistics for both cases are identical and the derivation for the empirical landscape automatically implies the result for the random field case. However, the derivation for the random field case can be greatly simplified due to the isotropic property of the distribution. Therefore, here we present the derivations for both cases separately.

### 6 Distance covariance of $k$-th order epistatic coefficients for random fields

In the last section, we defined a measure of local epistatic interaction $\epsilon^{S,\psi}$ over the $p$-dimensional hypercube $C_x^{S,\psi}$. In this section, we derive the covariance of $\epsilon^{S,\psi}$ as a function of Hamming distance under the Gaussian random field model $\mathbf{f} \sim \mathcal{N}(\mathbf{0}, \sum_{k=0}^{\ell} \lambda_k \mathbf{W}_k)$ defined in Section 4.

First, recall the hypercube $C_x^{S,\psi}$ that has the sequence $x$ as the "wild type" and variable positions and mutant alleles specified by $S$ and $\psi$. Let $x^{T,\tau}$ be a sequence that differs from $x$ at positions $T = \{T_1, \cdots, T_d\} \subset \{1, \cdots, \ell\} \setminus S$, with mutant alleles specified by $\tau \in A \setminus \{x_{T_1}\} \times \cdots \times A \setminus \{x_{T_d}\}$. We can construct a new cube $C_{x^{T,\tau}}^{S,\psi}$, with the same set of mutations at positions $S$. Geometrically, we can construct $C_{x^{T,\tau}}^{S,\psi}$ by moving $C_x^{S,\psi}$ in $d$ different direction, so that that new cube $C_{x^{T,\tau}}^{S,\psi}$ is anchored at the sequence $x^{T,\tau}$. Our random field model specifies the statistical similarity in values between sequences that are a certain number of mutations away. This notion of distance can be naturally generalized to geometric objects like $C_x^{S,\psi}$, since we can establish a one-to-one correspondence between sequences on $C_x^{S,\psi}$ and $C_{x^{T,\tau}}^{S,\psi}$, so that their Hamming distance is uniformly $d$. We now proceed to derive the covariance of the local epistatic coefficients defined over the two cubes.

Using $\mathbb{E}_{\mathbf{f}}$ to denote expectation over the distribution of $\mathbf{f}$, it is easy to verify that $\mathbb{E}_{\mathbf{f}}\left[\epsilon_x^{S,\psi}(\mathbf{f})\right] = 0$, given that the random vector $\mathbf{f}$ has mean $\mathbf{0}$. Next, to simplify notation, let $x_s = x^{s,\psi_s}$ denote the sequence that have the mutant alleles specified by $\psi$ at positions $s \subset S$ in the $x$ background, and $x'_s$ similarly denote the mutant sequence in the $x^{T,\tau}$ background (i.e. $x'_s = (x^{T,\tau})^{s,\psi_s}$). The covariance between the $k$-th order epistatic coefficients defined over these two cubes is then

$$\mathbb{E}_{\mathbf{f}}\left[\epsilon_x^{S,\psi}(\mathbf{f})\epsilon_{x^{T,\tau}}^{S,\psi}(\mathbf{f})\right] = \mathbb{E}_{\mathbf{f}}\left[(\sum_{s\subset S}(-1)^{|s|}f(x_s))(\sum_{t\subset S}(-1)^{|t|}f(x'_t))\right] \tag{73}$$

$$= \mathbb{E}_{\mathbf{f}}\left[\sum_{s,t\subset S}(-1)^{|s|+|t|}f(x_s)f(x'_t)\right] \tag{74}$$

$$= \mathbb{E}_{\mathbf{f}}\left[\sum_{s\subset S}(-1)^{|s|}\sum_{t\subset S}(-1)^{|t|}f(x_s)f(x'_t)\right] \tag{75}$$

$$= \sum_{s\subset S}(-1)^{|s|}\sum_{t\subset S}(-1)^{|t|}\mathbb{E}_{\mathbf{f}}\left[f(x_s)f(x'_t)\right] \tag{76}$$

$$= \sum_{s\subset S}\sum_{q=0}^{k}(-1)^q\binom{k}{q}K(d+q) \tag{77}$$

$$= 2^k\sum_{q=0}^{k}(-1)^q\binom{k}{q}K(d+q) \equiv \Gamma_k(d). \tag{78}$$

The quantity $\Gamma_k(d)$ is defined for $0 \leq d \leq \ell - k$, where the upper bound is limited by the number of sites to mutate without changing the subcube configuration. Moreover, the vector $\left[\Gamma_k(d)\right]_{0\leq d\leq \ell-k}$ is a simple linear transformation of the autocovariance function $K$ of the random field and measures how far we can generalize an observed epistatic coefficient as we traverse the sequence space.

### 7 Distance covariance of $k$-th order epistatic coefficients in an empirical landscape

In this section we derive the autocovariance function of $k$th-order epistatic coefficients for a definite empirical landscape $\mathbf{f}$, similar to $\Gamma_k$ for the case of a random field in the previous section.

We first find that the sum of $k$th-order epistatic coefficients over all anchor sequences and $k$-cubes is 0:

$$\sum_x \sum_S \sum_\psi \epsilon_x^{S,\psi}(\mathbf{f}) = (-1)^k \sum_x f(x) \sum_{q=0}^{k} (-1)^q \binom{\ell}{q} (\alpha-1)^q \binom{l-q}{k-q} (\alpha-1)^{k-q} \tag{79}$$

$$= (\alpha-1)^k \sum_x f(x) \sum_{q=0}^{k} (-1)^q \binom{\ell}{q} \binom{l-q}{k-q} \tag{80}$$

$$= (\alpha-1)^k \sum_x f(x) \binom{\ell}{k} \sum_{q=0}^{k} (-1)^q \binom{k}{q} = 0 \tag{81}$$

Note that, following [15], in the above expression we are averaging over all sequence backgrounds and polarities of mutation which by symmetry naturally sets the mean $k$th-order epistatic coefficient to 0.

Now we define the empirical covariance of $k$-th order epistatic coefficient for a landscape $f$ as

$$\Gamma_k(d) = \frac{1}{p_k^d} \sum_x \sum_S \sum_\psi \sum_T \sum_\tau \epsilon_x^{S,\psi}(\mathbf{f}) \epsilon_{x^{T,\tau}}^{S,\psi}(\mathbf{f}), \tag{82}$$

where $T = \{T_1, \cdots, T_d\} \subset \{1, \cdots, \ell\} \setminus S$ is a subset of sites with mutations specified by $\tau \in A \setminus \{x_{T_1}\} \times \cdots \times A \setminus \{x_{T_d}\}$. We find that the normalizing constant (the number of pairs of $k$-cubes enumerated in the sum) is given by $p_k^d = \alpha^\ell \binom{\ell}{k} \binom{\ell-k}{d} (\alpha-1)^{k+d}$. Note that we have used $\Gamma_k$ to denote the autocovariance function of epistatic coefficients of both random fields and empirical landscapes for convenience. The meaning of $\Gamma_k$ should be obvious from the context or explicitly specified throughout the paper. For the special case of $k=1$, $\Gamma_k$ reduces to the distance covariance function of mutational effects of an empirical landscape, and was first derived in ref. [15], and applied to an empirical data set in ref. [16]. Here we have generalized this quantity to epistatic coefficients of all orders.

Suppose two sequences differ at $d+q$ sites, then the number of pairs of $k$-cubes at distance $d$ where these two sequences occur respectively in the sum of Eq. 82 is

$$\pi_{k,d}^{d+q} = 2^k \binom{d+q}{d} \binom{\ell-d-q}{k-q} (\alpha-1)^{k-q}. \tag{83}$$

Continuing from Eq. 82 and changing the summation to be over pairs of sequences at different distances, we get

$$\Gamma_k(d) = \frac{1}{p_k^d} \sum_{q=0}^{k} \sum_{D(x,x')=d+q} \pi_{k,d}^{d+q} (-1)^q f(x) f(x') \tag{84}$$

$$= \sum_{q=0}^{k} (-1)^q \frac{\pi_{k,d}^{d+q}}{p_k^d} \sum_{D(x,x')=d+q} f(x) f(x') \tag{85}$$

$$= 2^k \sum_{q=0}^{k} (-1)^q \frac{\binom{k}{q}}{\alpha^\ell \binom{\ell}{d+q} (\alpha-1)^{d+q}} \sum_{D(x,x')=d+q} f(x) f(x') \tag{86}$$

From Line 85 to 86, we have used the identity

$$\frac{\pi_{k,d}^{d+q}}{p_k^d} = \frac{2^k \binom{d+q}{d} \binom{\ell-d-q}{k-q} (\alpha-1)^{k-q}}{\alpha^\ell \binom{\ell}{k} \binom{\ell-k}{d} (\alpha-1)^{k+d}} = \frac{2^k \binom{k}{q}}{\alpha^\ell \binom{\ell}{d+q} (\alpha-1)^{d+q}}. \tag{87}$$

Furthermore, since $\alpha^\ell \binom{\ell}{d+q} (\alpha-1)^{d+q}$ is equal to the number of pairs of sequences at distance $d+q$, we find

$$\frac{1}{\alpha^\ell \binom{\ell}{d+q} (\alpha-1)^{d+q}} \sum_{D(x,x')=d+q} f(x) f(x') = \langle f(x) f(x') \rangle_{D(x,x')=d+q}, \tag{88}$$

where we use $\langle \cdot \rangle_{D(x,x')=d+q}$, to denote the mean taken over all pairs of sequences satisfying $D(x,x') = d+q$. Therefore, we find

$$\Gamma_k(d) = 2^k \sum_{q=0}^{k} (-1)^q \binom{k}{q} \langle f(x) f(x') \rangle_{D(x,x')=d+q} \tag{89}$$

$$= 2^k \sum_{q=0}^{k} (-1)^q \binom{k}{q} (C(d+q) + \bar{f}^2), \tag{90}$$

based on the definition of $C(d)$ in Eq. 40. Assuming $\mathbf{f}$ has zero mean, we get a result very similar to Eq. 78 in the random field case:

$$\Gamma_k(d) = 2^k \sum_{q=0}^{k} (-1)^q \binom{k}{q} C(d+q). \tag{91}$$

The main difference is that here we replaced the kernel function $K(d)$ with the empirical autocovariance function $C(d)$.

Lastly, for the special case $k = 1$, we arrive at the distance covariance of mutational effects

$$\Gamma_1(d) = 2(C(d) - C(d+1)). \tag{92}$$

We can also calculate the corresponding distance correlation function by dividing this quantity with $\Gamma_1(0)$ (the variance of mutational effects):

$$\frac{\Gamma_1(d)}{\Gamma_1(0)} = \frac{2(C(d) - C(d+1))}{2(C(0) - C(1))} = \frac{\rho(d) - \rho(d+1)}{1 - \rho(1)}, \tag{93}$$

which is identical to the expression for $\gamma_d$ in ref. [10].

### 8 Relation between the $\lambda_k$ and the distance correlation of $k$-th order epistatic coefficients $\Gamma_k$

In the previous two sections, we have shown that the distance covariance function $\Gamma_k(d)$ of $k$-th order epistatic coefficients can be obtained from the kernel function $K(d)$ of a random field or the empirical autocovariance function $C(d)$ of a full landscape $\mathbf{f} \in \mathbb{R}^{\alpha^\ell}$. Moreover, in both cases, given the mean squared regression coefficients, $\lambda_0, \lambda_1, \cdots, \lambda_\ell$, we can calculate $\Gamma_k(d)$ using $\lambda_k, \lambda_{k+1}, \cdots, \lambda_\ell$. We summarize this result more formally with the following proposition.

**Proposition 1.** *Let $\lambda_0, \lambda_1, \cdots, \lambda_\ell$ be the mean squared regression coefficients for orders 0 through $\ell$ of a random field or an empirical landscape. The corresponding distance covariance of k-th order epistatic coefficients is*

$$\Gamma_k(d) = 2^k \sum_{j=0}^{\ell-k} \lambda_{k+j} w_j^{\ell-k}(d), \tag{94}$$

*where $w_j^{\ell-k}(d)$ is the j-th order Krawtchouk polynomial for a sequence space of length $\ell - k$.*

*Proof.* First, recall that in the case of random field landscapes the kernel function can be written as

$$K(d) = \sum_{k=0}^{\ell} \lambda_k w_k^\ell(d). \tag{95}$$

Similarly, in the case of an empirical landscape, the uncentered autocovariance function can be written as

$$C(d) + \overline{f}^2 = \sum_{k=1}^{\ell} \lambda_k w_k^\ell(d) + \alpha^\ell \, \overline{f}^2 \frac{1}{\alpha^\ell} = \sum_{k=1}^{\ell} \lambda_k w_k^\ell(d) + \lambda_0 w_0^\ell(d) = \sum_{k=0}^{\ell} \lambda_k w_k^\ell(d), \tag{96}$$

using Eqs. 10, 13, and 48.

In Section 15, Proposition 2, we will show that the Krawtchouk polynomial satisfies the identity

$$\sum_{p=0}^{m} (-1)^p \binom{m}{p} w_k^\ell(d+p) = w_{k-m}^{\ell-m}(d).$$

Using this property, we then find that:

$$\Gamma_k(d) = 2^k \sum_{q=0}^{k} (-1)^q \binom{k}{q} \sum_{k'=1}^{\ell} \lambda_{k'} w_{k'}^\ell(d+q) \tag{97}$$

$$= 2^k \sum_{k'=1}^{\ell} \lambda_{k'} \sum_{q=0}^{k} (-1)^q \binom{k}{q} w_{k'}^\ell(d+q) \tag{98}$$

$$= 2^k \sum_{j=0}^{\ell-k} \lambda_{k+j} w_j^{\ell-k}(d). \tag{99}$$

□

We can also write down the distance correlation function for $k$-th order local epistatic coefficients, namely $\gamma_k(d)$, by dividing $\Gamma_k(d)$ with $\Gamma_k(0)$, i.e. the variance of local $k$-th order epistatic coefficients. This gives us:

$$\gamma_k(d) = \frac{\Gamma_k(d)}{\Gamma_k(0)} \tag{100}$$

$$= 2^k \sum_{j=0}^{\ell-k} \frac{\lambda_{k+j}}{\sum_{n=0}^{\ell-k} \lambda_{k+n} w_n^{\ell-k}(0)} w_j^{\ell-k}(d) \tag{101}$$

$$= 2^k \sum_{j=0}^{\ell-k} \frac{\lambda_{k+j} m_j^{\ell-k}}{\sum_{n=0}^{\ell-k} \lambda_{k+n} m_n^{\ell-k}} \frac{1}{m_j^{\ell-k}} w_j^{\ell-k}(d) \tag{102}$$

$$= 2^k \sum_{j=0}^{\ell-k} v_j^{\ell-k} \rho_j^{\ell-k}(d). \tag{103}$$

Here we have used the fact that $w_n^{\ell-k}(0) = m_n^{\ell-k} = \binom{\ell-k}{n}(\alpha-1)^n$. So $w_j^{\ell-k}(d)/m_j^{\ell-k} = \rho_j^{\ell-k}(d)$ can be viewed as the distance correlation function of a pure $j$-th order function in a sequence space of length $\ell-k$. Lastly, we find that $v_j^{\ell-k} = \frac{\lambda_{k+j} m_j^{\ell-k}}{\sum_{j=0}^{\ell-k} \lambda_{k+n} m_n^{\ell-k}}$, is equal to the variance component of $j$th-order interactions in a sequence space of length $\ell - k$, where we calculate the variance components using $\lambda_k, \lambda_{k+1}, \ldots, \lambda_\ell$ and their corresponding multiplicities in this space of shorter sequences.

### 9 Connection with $L_2$-regularized regression and Minimum Epistasis Interpolation

In the main text, we claim that our method generalizes common regression models by having them as special cases or as models that arise in particular limits of our modeling framework. Here we make this connection precise. First consider the log posterior probability given the data $\mathbf{y}$:

$$-\log(p(\mathbf{f}|\mathbf{y})) = \mathbf{f}^T\mathbf{K}^{-1}\mathbf{f} + (\mathbf{f}_B - \mathbf{y})^T\mathbf{E}^{-1}(\mathbf{f}_B - \mathbf{y}) + \text{constant}, \tag{104}$$

where the first term corresponds to the prior density and the second term the data likelihood. Since $\mathbf{K} = \sum_{k=0}^{\ell} \lambda_k \mathbf{W}_k$ is also the eigendecomposition of $\mathbf{K}$, we can decompose the first term as $\mathbf{f}^T\mathbf{K}^{-1}\mathbf{f} = \sum_{k=0}^{\ell} \frac{1}{\lambda_k}\mathbf{f}^T\mathbf{W}_k\mathbf{f} = \sum_{k=0}^{\ell} \frac{1}{\lambda_k}\|\mathbf{f}_k\|^2$, where $\mathbf{f}_k$ is the component of $\mathbf{f}$ of $k$-th order interaction (see section 3) and $\|\mathbf{f}_k\|^2$ measures the sum of squared regression coefficients of order $k$ under any orthonormal basis of $V_k$. Now it becomes transparent that our method can be viewed as a particular type of $L_2$-regularized regression [17], where we penalize different components of $\mathbf{f}$ corresponding to different interaction orders according to the hyperparameters $\lambda_k$ that we specify. Since we use the negative log posterior probability as the cost function in this regularization problem, the $\mathbf{f}$ that minimizes Eq. 104 is also the mode of the posterior distribution (i.e., the MAP estimate).

For example, to get a regularized pairwise model from Eq. 104, we can set $\lambda_k = c$ for $k > 2$ and take the limit as $c \to 0$. This corresponds to assigning zero variance in the prior to these variance components or equivalently giving infinite penalties to interaction terms of order larger than 2. The remaining nonzero $\lambda_k$, $k \leq 2$, can then be set to values that correspond to different regularization strengths among the orders, where the ratios between $\lambda_0, \lambda_1, \lambda_2$ correspond to specific choices of design matrices whose columns span the space of interactions up to order 2 and their absolute scale serves as the inverse of the regularization parameter (i.e. larger $\lambda_k$ correspond to less regularization). As one example, recall that $\mathbf{W}_k = \mathbf{Q}_k\mathbf{Q}_k^T$, where $\mathbf{Q}_k$ is any matrix with orthonormal columns that spans the space of pure $k$th order interactions. So we have $\mathbf{f}^T\mathbf{W}_k\mathbf{f} = \mathbf{f}^T\mathbf{Q}_k\mathbf{Q}_k^T\mathbf{f} = \|\boldsymbol{\beta}_{\boldsymbol{k}}\|^2$. Therefore, setting the $\lambda_k$ for $k = 0, 1, 2$ to be equal turns the problem of maximizing Eq. 104 into an $L_2$-regularized regression problem that uniformly penalizes the regression coefficients $\boldsymbol{\beta}_{\boldsymbol{k}} = \mathbf{Q}_k^T\mathbf{f}$ for $k = 0, 1, 2$ under any orthonormal basis of $Y_2$. Generally speaking, we can derive the corresponding $\lambda_0, \lambda_1, \lambda_2$, for $L_2$ regularization for choices of design matrix $\mathbf{X}$ that exhibits sufficient symmetry (specifically, we require that the entries of the matrix $\mathbf{X}\mathbf{X}^T$ depend only on Hamming distance). For instance, in Supplemental Figure 2, we show the $\lambda_0, \lambda_1, \lambda_2$, that correspond to a regularized pariwise model that uniformly penalizes regression coefficients under a one-hot design matrix that contains 1st and 2nd order terms.

Eq. 104 also helps elucidate the connection between empirical variance component regression and our previously proposed method, minimum epistasis interpolation [14]. To see the connection, we first recall the regularized version of minimum epistasis interpolation with the cost function

$$\mathbf{f}^T\mathbf{C}\mathbf{f} + \omega(\mathbf{f}_B - \mathbf{y})^T\mathbf{E}^{-1}(\mathbf{f}_B - \mathbf{y}), \tag{105}$$

where $\mathbf{C} = \mathbf{L}^2 - \alpha\mathbf{L}$ is a special matrix that measures the sum of the squared epistatic coefficients over all 2-dimensional faces of the landscape $\mathbf{f}$, and $\omega$ is a positive weight that specify the relative importance of the regularization term and the squared loss term. Therefore, we can interpret Eq. 105 as a negative log posterior probability of the same form as Eq. 104, under the prior distribution specified by the precision matrix $\mathbf{C}$ and Gaussian noise with covariance matrix $\frac{1}{\omega}\mathbf{E}$.

Recall that the eigenvalue of $\mathbf{L}$ associated with the space of functions of interaction order $k$ is $\alpha k$. Given that $\mathbf{C} = \mathbf{L}^2 - \alpha\mathbf{L}$, the eigenvalues for $\mathbf{C}$ are $\alpha^2(k^2 - k)$, so that the corresponding $\lambda_k$ are given by $1/\alpha^2(k^2 - k)$ for $k > 2$. However, the constant and additive components are unregularized since $\alpha^2(k^2 - k) = 0$ for $k = 0, 1$. This is consistent with the definition of $\mathbf{C}$ as the matrix for calculating sum of squared epistatic coefficients, since a constant or additive landscape has zero epistatic coefficient everywhere. As a result, $\mathbf{C}$ in Eq. 105 actually defines an improper prior that arises as the limit when $\lambda_k \to \infty$ for $k = 0, 1$ while $\lambda_k = 1/\alpha^2(k^2 - k)$ for $k > 2$. Note that similar priors can also be constructed based on higher-order local interactions, see [18] for an application of such priors for conducting density estimation in sequence space.

Lastly, if we take the limit $\omega \to \infty$, the solution that minimizes Eq. 105 will perfectly reconstruct the data, while at the same time having the minimum total squared epistasis over all faces of the reconstructed landscape. This allows us to find the minimum epistasis interpolation solution.

### 10 Inference of hyperparameters for the prior distribution

To use Gaussian process regression, we must choose the covariance matrix $\mathbf{K}$, which in our case can be specified by a kernel function $K(d)$ for $d = 0, 1, \cdots, \ell$. According to Eq. 52, the kernel function $K(d)$ of our prior distribution must take the form

$$K(d) = \sum_{k=0}^{\ell} \lambda_k w_k^{\ell}(d). \tag{106}$$

In this paper, we take an Empirical Bayes approach by using point estimates inferred from the data to set the hyperparameters $\lambda_k$. First, recall that $c(d)$ is the empirical autocovariance function extracted from the data $\mathbf{y}$ where $c(0)$ has had the average noise variance removed (Eq. 49). We would like to choose the hyperparameters $\lambda_k$ so that the kernel function $K(d)$ aligns as well as possible with $c(d)$. A naive method for finding such $\lambda_k$ is by solving the system of linear equations

$$c(d) + \overline{y}^2 = \sum_{k=0}^{\ell} \lambda_k w_k^{\ell}(d), \quad d = 0, 1, \cdots, \ell \tag{107}$$

for the hyperparameters $\lambda_k$. Note that on the left hand side of Eq. 107, we have added $\overline{y}^2$ to $c(d)$. This is required because $c(d)$ is the (centered) distance covariance function, which does not contain any information about the mean and so $c(d)+\overline{y}^2$ is the uncentered analog (i.e. distance function of the second raw moment), which must be used here in order to infer the 0-th order hyperparameter $\lambda_0$. Note also that this naive procedure is equivalent to directly using $c(d)+\overline{y}^2$ as the kernel function $K(d)$ for our prior distribution.

An important constraint for the $\lambda_k$ is that they must be nonnegative, since they are the eigenvalues of the positive semi-definite matrix $\mathbf{K} = \sum_{k=0}^{\ell} \lambda_k \mathbf{W}_k$. While it has been shown that when $B$ is the whole sequence space and there is no measurement noise, the $\lambda_k$'s solved using the equation above must be nonnegative [5], no such guarantee exists when $B$ is a proper subset of $A^\ell$. Moreover, given only partial data it is also possible that a subset of distance classes are absent, so that Eq. 107 no longer has a unique solution. And, in addition, we would like to be able to incorporate our estimates of experimental uncertainty when constructing our prior.

In order to address these issues, we take a regularized weighted least-squares approach to optimally approximate the empirical autocovariance function. This method is similar to a machine learning technique called kernel alignment [19]. It is also similar to the Haseman-Elston Regression [20] commonly used in quantitative genetics for inferring variance components. Briefly, our strategy is to approximate the empirical second moment matrix $\mathbf{y}\mathbf{y}^T$ using a nonnegative linear combination of the basis matrices $\mathbf{W}_k$'s and the noise variance matrix $\mathbf{E}$. Mathematically, we achieve this by minimizing the squared Frobenius norm ($\|\cdot\|_F$) of the difference between the target matrix $\mathbf{y}\mathbf{y}^T$ and the submatrix $\mathbf{K}_{BB} = \sum_{k=0}^{\ell} \lambda_k \mathbf{W}_{k_{BB}}$, i.e.:

$$\|\mathbf{y}\mathbf{y}^T - (\sum_{k=0}^{\ell} \lambda_k \mathbf{W}_{k_{BB}} + \mathbf{E})\|_F^2. \tag{108}$$

It turns out this quantity is equal to the weighted sum of squared differences between the kernel function $K(d)$ and the uncentered empirical autocovariance function, up to an additive constant. To see this, define $t(d) = c(d) + \overline{y}^2$, where $c(d)$ is the empirical autocovariance function (Eq.49), so that again $t(d)$ can be viewed as the uncentered autocovariance function. We then find

$$\|\mathbf{y}\mathbf{y}^T - (\sum_{k=0}^{\ell} \lambda_k \mathbf{W}_{k_{BB}} + \mathbf{E})\|_F^2 = \sum_{i\in B} \left[(\mathbf{y}_i^2 - \mathbf{E}_{ii}^2) - K(0)\right]^2 + \sum_{d=1}^{\ell} \sum_{D(i,j)=d} \left[\mathbf{y}_i\mathbf{y}_j - K(d)\right]^2 \tag{109}$$

$$= \mathcal{D}_0 \left[t(0) - K(0)\right]^2 + \sum_{i\in B} \left[(\mathbf{y}_i^2 - \mathbf{E}_{ii}^2) - t(0)\right]^2 + \sum_{d=1}^{\ell} \mathcal{D}_d \left[t(d) - k(d)\right]^2 + \sum_{d=1}^{\ell} \sum_{D(i,j)=d} \left[\mathbf{y}_i\mathbf{y}_j - t(d)\right]^2 \tag{110}$$

$$= \sum_{d=0}^{\ell} \mathcal{D}_d \left[t(d) - \sum_{k=0}^{\ell} \lambda_k w_k(d)\right]^2 + \sum_{i\in B} \left[(\mathbf{y}_i^2 - \mathbf{E}_{ii}^2) - t(0)\right]^2 + \sum_{d=1}^{\ell} \sum_{D(i,j)=d} \left[\mathbf{y}_i\mathbf{y}_j - t(d)\right]^2. \tag{111}$$

Since the last two terms do not depend on the hyperparameters $\lambda_k$, they can be omitted in the fitting procedure. Thus, under this weighted least squares approach, we can find our hyperparameters $\lambda_k$ by minimizing the function:

$$\text{minimize}_{\boldsymbol{\lambda}} \quad \sum_{d=0}^{\ell} \mathcal{D}_d \left[t(d) - \sum_{k=0}^{\ell} \lambda_k w_k(d)\right]^2, \tag{112}$$

under the constraint that $\lambda_k \geq 0$, for all $k$.

Importantly, when we can exactly reconstruct $t(d)$ using all nonnegative $\lambda_k$'s, the least squared solution in Eq. 112 and naive solution to Eq. 107 fitted to the $t(d)$ of an incomplete landscape $f_B$ coincide. But when Eq. 107 gives negative $\lambda_k$'s, the optimal solution to Eq. 112 may include some $\lambda_k = 0$, which would result in a prior that is missing $k$-th order interactions. In addition, when not all distance classes are present in the data, the solution to Eq. 112 may not be unique. We address these issues by imposing regularization on the $\lambda_k$. Specifically, reparametrizing $\lambda_k = \exp(\eta_k)$, we solve the minimization problem:

$$\text{minimize}_{\boldsymbol{\eta}} \quad \sum_{d=0}^{\ell} \mathcal{D}_d \left[ t(d) - \sum_{k=0}^{\ell} \exp(\eta_k) w_k(d) \right]^2 + \beta \sum_{k=2}^{\ell-1} \|2\eta_k - \eta_{k-1} - \eta_{k+1}\|^2. \tag{113}$$

The first sum in Eq. 113 is identical to the function to be minimized in Eq. 112, but the problem is now an unconstraned minimization problem due to the reparametrization in terms of the $\eta_k$. The regularization term $\sum_{k=2}^{\ell-1} \|2\eta_k - \eta_{k-1} - \eta_{k+1}\|^2$ is equal to the sum of squared second order finite differences in $\boldsymbol{\eta}$. It thus ensures numerical stability when estimating $\lambda_1, \cdots, \lambda_\ell$ by encouraging them to conform to a linear function on a log scale (see [8] for a theoretical basis for using exponentially decaying $\lambda_k$ to model fitness landscapes). The value of $\beta > 0$ controls the strength of regularization. In practice, we choose the optimal $\beta$ via 10-fold cross validation, where we choose the value of $\beta$ that best fits the pattern of auto-correlation in the held-out fold. More specifically, we choose the value that minimizes the mean-squared difference between the empirical second moment matrix for the held-out fold and its predicted second moment matrix (i.e. we minimize the loss defined as Eq. 108 divided by the number of held-out sequences, where $\mathbf{y}$ are the phenotypic values for the held-out fold and the diagonal of $\mathbf{E}$ contains the corresponding noise variances).

For the analyses in the main text, we always use cross validation to estimate the $\lambda_k$, where our cross validation procedure only uses information about the distance-correlation structure of the data, in line with our empirical Bayesian approach and overall philosophy of considering how mutational effects, local epistatic coefficients, etc., change as one moves through sequence space. However, other approaches such as choosing the $\lambda_k$ by maximimizing the Bayesian evidence or conducting a hierarchical Bayesian approach would also be reasonable choices. Thus, in addition to the regularized estimation procedure, our software also supports naive estimation of the lambdas from Eq. 107, user specification of the regularization parameter $\beta$, and direct user specification of the $\lambda_k$.

### 11 Posterior sampling using Hamiltonian Monte Carlo

Eq. 60 allows us to calculate the posterior variance for individual sequences. However, since the evaluation of this function is as costly as the MAP estimate, in practice we can only acquire the posterior variance for a subset of sequences of high interest. In this section we outline an alternative method for estimating the posterior covariance matrix by directly sampling from the posterior distribution in Eq. 59. Specifically, suppose $\mathbf{f}^{(i)}$, $(i = 1, \cdots, n)$ is a set of $n$ samples drawn from the posterior distribution using a Markov chain whose stationary distribution is our posterior distribution, then we can approximate the posterior covariance matrix with the finite sum

$$\mathbf{K} - \mathbf{K}_{\cdot B}(\mathbf{K}_{BB} + \mathbf{E})^{-1}\mathbf{K}_{B\cdot} = \mathrm{E}\left[(\mathbf{f} - \widehat{\mathbf{f}})(\mathbf{f} - \widehat{\mathbf{f}})^T\right] \approx \frac{1}{n}\sum_{l=1}^{n}(\mathbf{f}^{(l)} - \overline{\mathbf{f}})(\mathbf{f}^{(l)} - \overline{\mathbf{f}})^T, \tag{114}$$

where $\overline{\mathbf{f}}$ is the mean vector taken over all samples $\mathbf{f}^{(i)}$.

A major challenge for sampling from the posterior distribution is posed by the typical high dimensionality of the sequence space. Specifically, as the dimension of the sample space increases, the region of high probability of the posterior distribution (the typical set) becomes increasingly singular and concentrated in space [21, 22]. As a consequence, the diffusive behavior of popular naive random walk algorithms such as MCMC either leads to high rejection rates or highly autocorrelated samples, both making the exploration of the probability distribution extremely slow.

In this paper, we employ the Hamiltonian Monte Carlo (HMC) sampling method [21, 22]. HMC is a gradient-based algorithm that is able to take advantage of the local geometry of the typical set, making it more suitable for sampling from high dimensional probability distributions. HMC first introduces an auxiliary momentum parameter to complement each dimension of our target probability space. The total energy (the Hamiltonian) of the system is then defined as the sum of the potential energy given by the log probability of the posterior and the kinetic energy, which is equal to the squared norm of the momentum vector in our case. The algorithm proceeds using a Markov chain consisting of alternate random updates to the momentum vector and deterministic integration of Hamiltonian dynamics that leaves the total energy unchanged. In practice, this integration is discretized and performed using the so-called leapfrog method [22]. Since the numerical errors accumulated during the leapfrog steps lead to changes in total energy at the end of the integration, a Metropolis step at the end of the numerical integration is used to keep the Markov chain reversible. Together, this sampling scheme allows the HMC algorithm to make large jumps in probability space while keeping the rejection rate small.

The HMC algorithm relies on the gradient of the log posterior probability to perform the Hamiltonian dynamics integration. To derive the gradient, first note that the precision matrix for the posterior distribution is given by

$$\widetilde{\mathbf{K}} = (\mathbf{K} - \mathbf{K}_{\cdot B}(\mathbf{K}_{BB} + \mathbf{E})^{-1}\mathbf{K}_{B\cdot}))^{-1}. \tag{115}$$

Since the log probability of a sample $\mathbf{f}$ is $\log(\mathbf{f}|\mathbf{y}) = -\frac{1}{2}(\mathbf{f} - \widehat{\mathbf{f}})^T\widetilde{\mathbf{K}}(\mathbf{f} - \widehat{\mathbf{f}}) + \text{costant}$, we find

$$\frac{1}{2}\nabla \log p(\mathbf{f}|\mathbf{y}) = -\widetilde{\mathbf{K}}\mathbf{f} + \widetilde{\mathbf{K}}\widehat{\mathbf{f}}. \tag{116}$$

Next, we can simplify the expression for $\widetilde{\mathbf{K}}$ by expanding the inverse using the Woodbury identity. This gives

$$\widetilde{\mathbf{K}} = \mathbf{K}^{-1} + \begin{bmatrix} \mathbf{E}^{-1} & 0 \\ 0 & 0 \end{bmatrix}. \tag{117}$$

Since Eq. 52 is also the eigendecomposition of $\mathbf{K}$ and all $\lambda_k$'s are constrained to be positive, the inverse of $\mathbf{K}$ exists and is equal to $\mathbf{K}^{-1} = \sum_{k=0}^{\ell} \frac{1}{\lambda_k}\mathbf{W}_k$. Therefore, the evaluation of Eq. 116 involves multiplying the sample $\mathbf{f}$ by a diagonal matrix and the matrix $\mathbf{K}^{-1}$, which can be greatly sped up using a representation of $\mathbf{K}^{-1}$ as a polynomial in the graph Laplacian $\mathbf{L}$, since the latter is a sparse matrix.

Finally, we employ the dual averaging algorithm [23] to find the optimal step size for the leapfrog integrator during an initial tuning phase, so that the average rejection rate of the Metropolis steps is near the optimal value of 0.65 [23].

### 12 Implementation of regularized pairwise and three-way interaction models

We use the following linear model to fit additive, pairwise and 3-way interaction models:

$$\widehat{f}(x) = \sum_j \beta_j \phi_j(x), \tag{118}$$

where the $\phi_j(x)$ are indicator variables encoding the presence or absence of particular alleles at certain sites in $x$ (i.e. we use a one-hot encoding). For the additive model, each $\phi_j(x)$ returns 1 if a given allele is present at a given site in genotype $x$ and 0 otherwise. For the pairwise and three-way models, $\phi_j(x)$ encodes the presence or absence of combinations of allelic states on pairs or triples of sites, respectively, taking the value 1 if all members of the combination are present and 0 otherwise. We can express Eq. (118) in matrix notation

$$\widehat{\mathbf{f}} = \mathbf{X}\boldsymbol{\beta}. \tag{119}$$

Given $m$ observations, the dimension of $\mathbf{X}$ is $m \times (1 + \ell\alpha)$ for the additive model, $m \times \sum_{k=0}^{2} \binom{\ell}{k}\alpha^k$ for the pairwise model, and $m \times \sum_{k=0}^{3} \binom{\ell}{k}\alpha^k$ for the three-way model.

We fit the additive model using ordinary least squares. The pairwise and three-way regression models were fitted using elastic net regularization [24], where the penalty of model complexity is a mixture of $L_1$ and $L_2$ norms. Specifically, we find our solution by solving the following minimization problem:

$$\min_{\boldsymbol{\beta}\in\mathbb{R}^p} \|\mathbf{y} - \mathbf{X}\boldsymbol{\beta}\|_2 + \lambda \left((1-\alpha)\|\boldsymbol{\beta}\|_2^2/2 + \alpha\|\boldsymbol{\beta}\|_1\right), \tag{120}$$

where the penalty for model complexity is controlled by $\alpha$, which represents a compromise between lasso ($\alpha = 1$) and ridge ($\alpha = 0$) regressions. The parameter $\lambda$ controls the overall strength of the penalty. Both $\alpha$ and $\lambda$ were chosen by 10-fold cross validation for each training sample. Elastic net regressions were fit using the R package glmnet [25].

### 13 Symmetry properties of the Krawtchouk Polynomial

In this paper, we use several known properties of the Krawtchouk polynomial regarding its symmetries. These includes the symmetry in its signs between the $k$ and $d$ parameter, which is important for dividing the interactions into the locally correlated and anticorrelated groups shown in Figure 2 in the main text. Here we present the proofs for these results for the reader's convenience.

First recall the definition of Krawtchouk polynomial in $d$ of order $k$ with parameters $\alpha$ (number of alleles per site) and $\ell$ (number of sites):

$$w_k^\ell(d) = \alpha^{-\ell}\sum_{q=0}^{k}(-1)^q(\alpha-1)^{k-q}\binom{d}{q}\binom{\ell-d}{k-q}. \tag{121}$$

The Krawtchouk polynomials are a class of orthogonal polynomials central to coding theory [4] and spectral fitness landscape theory [9]. First we show that the Krawtchouk polynomials are symmetric in $k$ and $d$ with respect to the coefficient $m_p = (\alpha-1)^p\binom{\ell}{p}$:

$$(\alpha-1)^d\binom{\ell}{d}w_k^\ell(d) = (\alpha-1)^k\binom{\ell}{k}w_d^\ell(k). \tag{122}$$

To see how we arrive at this property, first note that $\binom{\ell}{d}\binom{d}{q}\binom{\ell-d}{k-q} = \binom{\ell}{k}\binom{k}{q}\binom{\ell-k}{d-q}$. Therefore,

$$\frac{(\alpha-1)^d\binom{\ell}{d}}{(\alpha-1)^k\binom{\ell}{k}}w_k^\ell(d) = \frac{(\alpha-1)^d\binom{\ell}{d}}{(\alpha-1)^k\binom{\ell}{k}}\alpha^{-\ell}\sum_{q=0}^{\ell}(-1)^q(\alpha-1)^{k-q}\binom{d}{q}\binom{\ell-d}{k-q} \tag{123}$$

$$= \alpha^{-\ell}\sum_{q=0}^{\ell}(-1)^q(\alpha-1)^{d-q}\frac{\binom{\ell}{d}}{\binom{\ell}{k}}\binom{d}{q}\binom{\ell-d}{k-q} \tag{124}$$

$$= \alpha^{-\ell}\sum_{q=0}^{\ell}(-1)^q(\alpha-1)^{d-q}\binom{k}{q}\binom{\ell-k}{d-q} = w_d^\ell(k) \tag{125}$$

Next, we recall the well-known orthogonality property of the Krawtchouk polynomial [26]:

$$\sum_{d=0}^{\ell} m_d w_k^\ell(d) w_{k'}^\ell(d) = \alpha^{-\ell}\delta_{k,k'}m_k. \tag{126}$$

Using the symmetry property we proved above, we can rewrite this equation as

$$\sum_{d=0}^{\ell} m_d w_k^\ell(d) w_{k'}^\ell(d) = \sum_{d=0}^{\ell} m_k w_d^\ell(k) w_{k'}^\ell(d) = m_k \sum_{d=0}^{\ell} w_d^\ell(k) w_{k'}^\ell(d) = \alpha^{-\ell} \delta_{k,k'} m_k. \tag{127}$$

Therefore, we find

$$\sum_{d=0}^{\ell} w_d^\ell(k) w_{k'}^\ell(d) = \delta_{k,k'} \alpha^{-\ell}. \tag{128}$$

Now let $W^{(\ell)}$ denote the $(\ell+1) \times (\ell+1)$ matrix with $W^{(\ell)}(j,k) = w_k^\ell(j)$. Then Eq. 128 shows

$$W^{(\ell)} W^{(\ell)} = \alpha^{-\ell} I_{\ell+1}. \tag{129}$$

### 14 Proof for the classification of interactions into locally correlated and anticorrelated groups

Now we use the results from the previous section to show that we can classify pure $k$-th order interactions into a locally correlated group and a locally anti-correlated group. Specifically, we say that a $k$-th order interaction contributes to positive local correlations between phenotypes if its distance covariance function is positive at distance 1, i.e. if $w_k(1) > 0$. Examining the sign of this function is not a trivial task when $k$ is large. However, since $\mathrm{sgn}(w_k^\ell(1)) = \mathrm{sgn}(w_1^\ell(k))$ based on the result above, we can instead examine the function $w_1^\ell(k)$, which is equal to

$$w_1^\ell(k) = \alpha^{-\ell} \left( (\alpha - 1)(\ell - k) - k \right) = \alpha^{-\ell} \left( -\alpha k + (\alpha - 1)\ell \right). \tag{130}$$

We find that $w_1^\ell(k)$ is linear and decreasing in $k$. Therefore, we can find a threshold value $p$, above which the distance 1 covariance becomes negative. Setting Eq. 130 to zero and solving for $k$, we find $p = (1 - \frac{1}{\alpha})\ell$. This threshold has a nice interpretation in that it is equal to the average number of different sites between two randomly chosen sequences. So if $k < p$, the interaction contributes to locally positive correlations. And if $k > p$, the interaction contributes to negative locally negative correlations.

However, $p$ can take integer values given certain $\alpha$ and $\ell$ combinations. So in the case $k = p$, we have a zero covariance at distance 1. To study the behavior of such autocovariance functions, we must look at the sign of the autocovariance funciton at distance 2, i.e. $\mathrm{sgn}(w_k(2))$, which is equal to $\mathrm{sgn}(w_2(k))$. Using Eq. 121, we find

$$w_2(k) = \frac{1}{2}(\alpha - 1)^2 (\ell - k)(\ell - m - 1) - (\alpha - 1)m(\ell - m) + \frac{1}{2} m(m-1). \tag{131}$$

Setting $k = (1 - \frac{1}{\alpha})\ell$, the equation above is equal to

$$-\frac{1}{2}(\alpha - 1)\ell, \tag{132}$$

which is always negative. Therefore, when $k$ equals the expected mutational distance between two random sequences, the $k$-th order interactions do not contribute any correlation between sequences at distance 1 but induce negative correlations at distance 2, so that this order of interactions is best viewed as being locally anti-correlated.

### 15 Proof for results used in Proposition 1

Here we present a novel recursive relation between Krawtchouk polynomials of different orders. This result allows us to study the geometry of the distance correlation in mutational effects, and local epistatic interactions in terms of Krawtchouk polynomials of a lower order, and is used to prove Proposition 1.

**Proposition 2.** *For $m < \ell$ and $d \leq \ell - m$, we have*

$$\sum_{p=0}^{m}(-1)^p\binom{m}{p}w_k^{\ell}(d+p) = \begin{cases} w_{k-m}^{\ell-m}(d) & k \geq m \\ 0 & k < m. \end{cases} \tag{133}$$

To prove this property, we first present the following result.

**Lemma 3.** *Given a polynomial in $x$*

$$(1+(\alpha-1)x)^{\ell-d}(1-x)^d, \tag{134}$$

*the coefficient for $x^k$ is given by the Krawtchouk polynomial in $d$*

$$\sum_{q=0}^{k}(-1)^q(\alpha-1)^{k-q}\binom{d}{q}\binom{\ell-d}{k-q} = \alpha^{-\ell}w_k^{\ell}(d), \tag{135}$$

*according to Eq. 121.*

To prove this, simply expand Eq. 134. Now for the proof of Proposition 2.

*Proof.* Consider the polynomial

$$\sum_{p=0}^{m}(-1)^p\binom{m}{p}(1+(\alpha-1)x)^{\ell-d-p}(1-x)^{d+p}. \tag{136}$$

The coefficient of $x^k$ of this polynomial is

$$\alpha^{-\ell}\sum_{p=0}^{m}(-1)^p\binom{m}{p}w_k^{\ell}(d+p) \tag{137}$$

according to Lemma 3. Now we can rewrite the polynomial in line 136

$$\sum_{p=0}^{m}(-1)^p\binom{m}{p}(1+(\alpha-1)x)^{\ell-d-p}(1-x)^{d+p} \tag{138}$$

$$=(1+(\alpha-1)x)^{\ell-d-m}(1-x)^d\sum_{p=0}^{m}(-1)^p\binom{m}{p}(1+(\alpha-1)x)^{m-p}(1-x)^p \tag{139}$$

$$=(1+(\alpha-1)x)^{\ell-d-m}(1-x)^d(1+(\alpha-1)x-1+x)^m \tag{140}$$

$$=(1+(\alpha-1)x)^{\ell-d-m}(1-x)^d\alpha^m x^m. \tag{141}$$

The lowest degree of terms of the polynomial of the last line is $m$. So the coefficient for $x^k$ is equal to zero if $k < m$. For $k \geq m$, the coefficients for $x^k$ is

$$\alpha^m\alpha^{-\ell-m}w_{k-m}^{\ell-m}(d), \tag{142}$$

using Lemma 3. Since line 136 and line 141 are the same polynomial, we have established that

$$\sum_{p=0}^{m}(-1)^p\binom{m}{p}w_k^{\ell}(d+p) = w_{k-m}^{\ell-m}(d), \tag{143}$$

for $k \geq m$ and is equal to zero if $k < m$. □

### 16 Processing of the *SMN1* splicing dataset

The *SMN1* dataset consists of enrichment ratios (number of output reads/number of input reads) across three libraries, with each library measured in triplicate. Previous analysis discarded two replicates due to low sample quality. Since no library effect was detected [27], we consider the enrichment ratios across 7 replicates as independent samples. Depending on its presence or absence in each input sample, a splice site can have zero to 7 measured enrichment ratios. Out of the 32768 possible splice sites, 2036 are not represented in any replicates, and therefore are considered missing data.

We assume the enrichment ratios across replicates for a given genotype are log-normally distributed. First, for sequences with all positive ratios across $n$ replicates ($1 < n \leq 7$), we use the bias corrected geometric mean [28] as the estimate of the median enrichment ratio using the formula

$$\widehat{\mu} = \exp(\overline{y} - \widehat{\sigma}^2/2n), \tag{144}$$

where $\overline{y}$ and $\widehat{\sigma}^2$ are the arithmetic mean and sample variance of the log-transformed enrichment ratios, respectively. For sequences that have a measured enrichment ratio of zero for at least one replicate, the above equation is inapplicable, and we simply score that sequence as having a phenotype equal to the median of the enrichment ratios across replicates.

We then estimate the variance for this log-normal distribution using the standard formula

$$\sigma^2 = (\exp(\widehat{\sigma}^2) - 1)\exp(2\widehat{\mu} + \widehat{\sigma}^2) \tag{145}$$

For sequences with zero ratios and/or with only 1 replicate, we use the modified formula

$$\sigma^2 = (\exp(\overline{\widehat{\sigma}^2}) - 1)\exp(2\mu' + \overline{\widehat{\sigma}^2}), \tag{146}$$

where $\mu'$ is the log of the median of the enrichment ratios and $\overline{\widehat{\sigma}^2}$ is the mean $\widehat{\sigma}^2$ for all sequences with only positive ratios and at least two replicates.

### 17 Visualization of the *SMN1* splicing landscape

To derive a low dimensional representation of the splicing landscape, we consider a population evolving in continuous time under weak mutation [29–31] with natural selection acting to maintain splicing activity. We first used our method to reconstruct the full landscape consisting of 65536 sequences corresponding to all combinations of alleles at the eight variable positions of the 9-nt splice site. Note that the reconstructed landscape also includes sequences with A or G at the +2 position, which do not constitute valid splice sites. Since these sequences are nonetheless accessible through mutation, we include them but set the PSI of all such sequences to be zero. Next, exon-exon junction sequencing in the original study revealed that a secondary GU at the -2 and -1 positions can be preferentially used over the GU or GC at position +1 and +2 [27], leading to a frameshift in the mature mRNA. Therefore, we set the PSI of all such sequences to be zero. Last, to ensure an appropriate degree of realism for the evolutionary Markov chain, we truncate all predicted PSI values to be between 0 and 100. We model evolution as a continuous-time Markov chain where the population moves between sequences at each fixation event based on fitness values given by the modeled PSI. The rate matrix $\mathbf{Q}$ of the Markov chain is

$$\mathbf{Q}_{x,x'} = \begin{cases} \frac{1}{\alpha-1}\frac{c(f(x')-f(x))}{1-e^{-c(f(x')-f(x))}} & d(x,x')=1 \\ -\sum_{x''\neq x}\mathbf{Q}_{x,x''} & x=x' \\ 0 & \text{otherwise,} \end{cases} \tag{147}$$

where $c$ is the conversion factor that transforms PSI to scaled fitness (Malthusian fitness $\times$ $N_e$). We choose $c$ so that the expected PSI at stationarity is equal to 80. Time is scaled so that the total mutation rate per site is equal to 1. We use the right eigenvectors of $\mathbf{Q}$ associated with the 3 greatest nonzero eigenvalues as coordinates to embed the splicing landscape in three dimensions, where each eigenvector is scaled so that the weighted mean of its squared entries is equal to the relaxation time of the associated eigenmode where the weights are given by the frequency of each genotype at stationarity (see [32] for details). This allows our low-dimensional representation of the landscape to optimally capture the expected time for a population to evolve between sequences [32]. Note that although we included sequences with A or G at position +2 when calculating the embedding coordinates, for simplicity we omitted these sequences when plotting the final visualization.

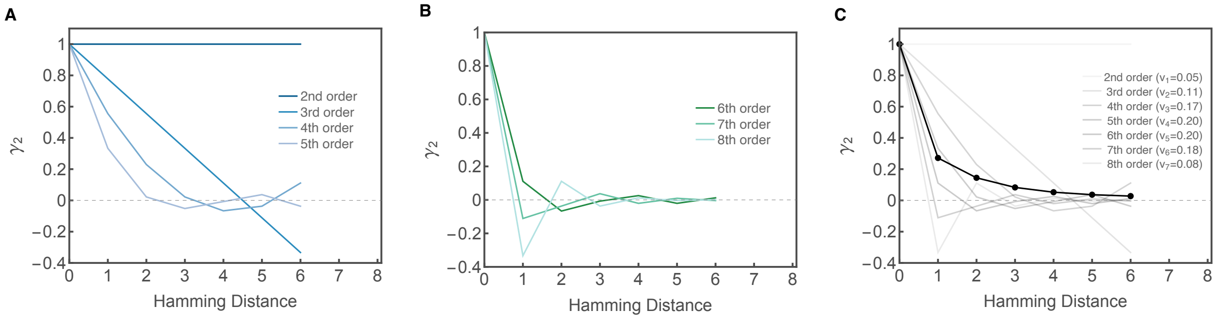


Supplementary Figure 1: Superimposition of distance correlation functions of double mutant epistatic coefficients ($\gamma_2$) for pure $k$-th order interactions for sequences of length 8 with 4 alleles per site. Note that the curves are plotted up to Hamming distance 6, since the maximal distance between two genetic backgrounds with the same pairwise epistatic interaction is 6. (A) Distance correlation function for orders of genetic interaction that produce locally positive correlations (in this case, interaction orders $k$=2–5). Note that the first order curve is not shown, since an additive landscape contains no local pairwise interaction. (B) Distance correlation function for orders of genetic interaction that produce local anti-correlations (in this case, $k$=6–8). (C) $\gamma_2$ of an arbitrary genotype-phenotype map (black line) is a weighted sum of elementary $\gamma_2$ functions (gray lines) with the weights given by the the variance components of mean squared epistatic coefficients of order $\geq 2$.

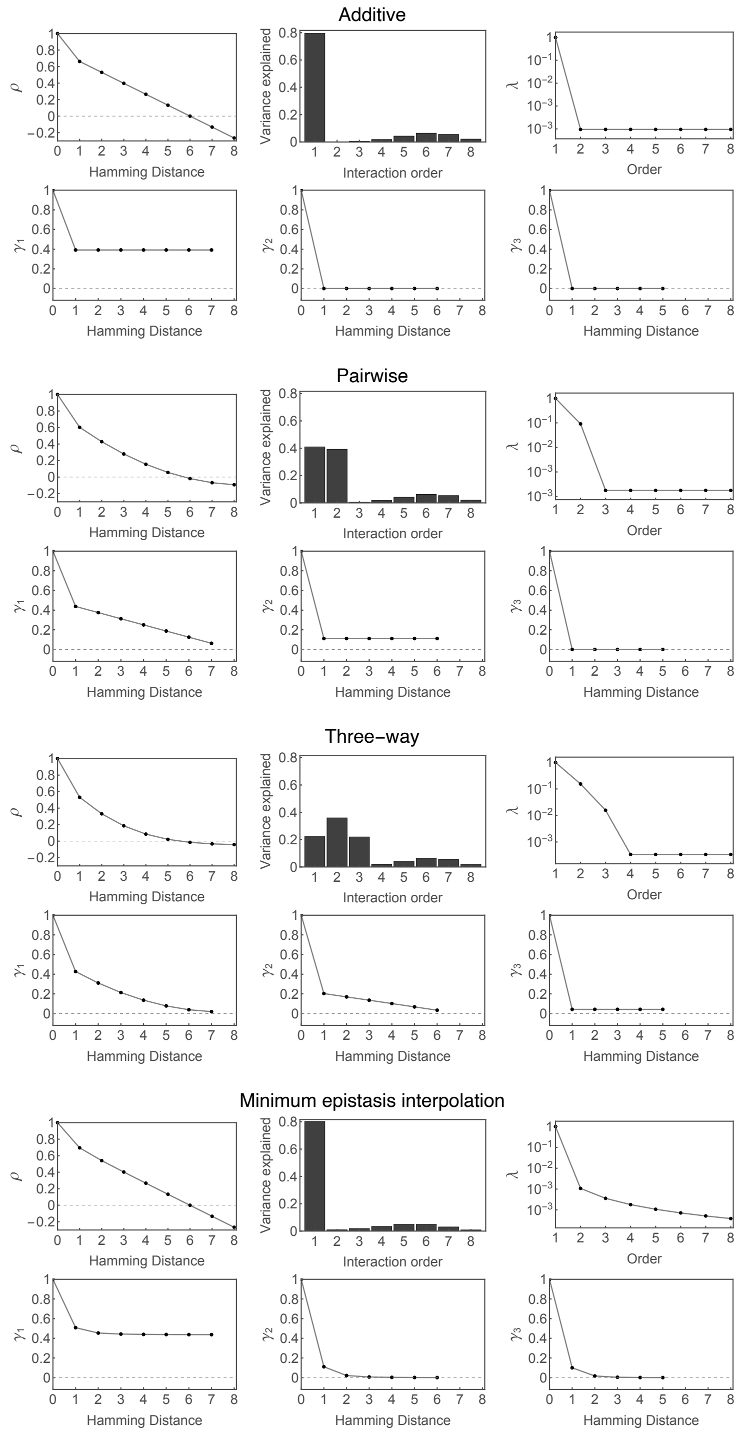


Supplementary Figure 2: Summary statistics for four different models. Each panel shows the empirical distance correlation function ($\rho$), the empirical variance components, mean square regression coefficient for each order ($\lambda$), and distance correlation of epistatic coefficients ($\gamma_k$) of order $k$ = 1-3. Quantities shown here represent the expected summary statistics of the prior distributions of different models, where the uniform interaction terms higher than the order of the model for the additive, pairwise and three-way model correspond to the independent error variance and the lambdas for the minimum epistasis interpolation model control the epistatic portion of the fit. For all models, the expected fraction of variance due to the lower order portion of the model ($k = 1$ for the additive and minimum epistasis interpolation models, $k = 1, 2$ for the pairwise model, and $k = 1, 2, 3$ for the three-way model) is 80%. The lower-order $\lambda_k$ for the additive, pairwise and three-way models were chosen to be equivalent to random field models constructed using one-hot vectors up to the highest order included in the model ($k = 1$ for the additive model, $k = 1, 2$ for the pairwise model, and $k = 1, 2, 3$ for the three-way model), where the coefficients multiplying these one-hot vectors are drawn independently from a standard normal distribution. See also SI Section 9 "Connection with $L_2$-regularized regression and Minimum Epistasis Interpolation".

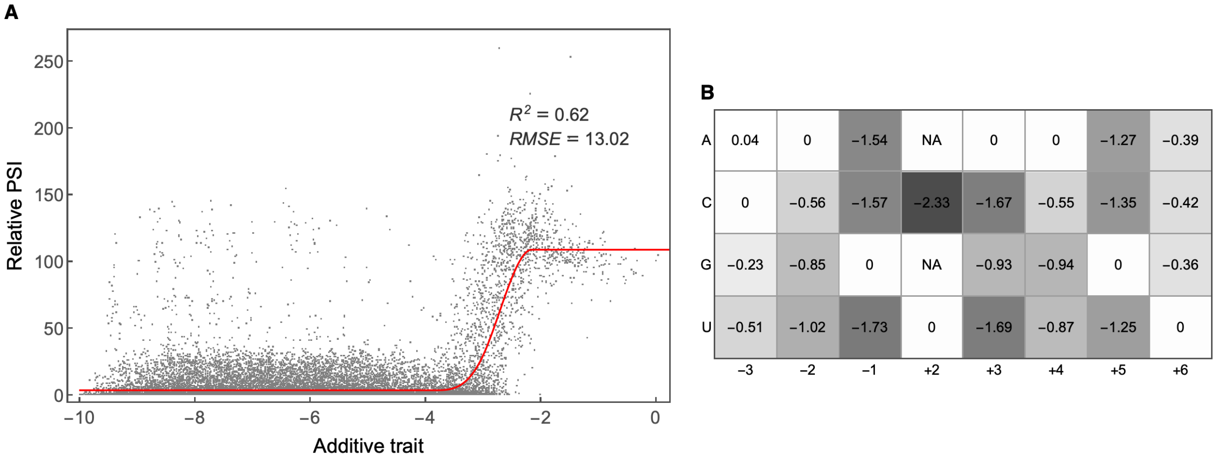


Supplementary Figure 3: Global epistasis model fitted to all 5′ splice sites in the *SMN1* dataset. The global epistasis model assumes there is an unobserved trait that is additive with respect to the effects of mutations and that the observed phenotype is a monotonic nonlinear function of this latent additive trait. (A) shows the relative PSI vs. the inferred additive trait value. Red curve is the inferred nonlinear function from the additive trait to the measurement (PSI) scale. Also shown are the in-sample $R^2$ and Root Mean Square Error ($RMSE$). (B) Mutational effects for the additive trait calculated relative to the consensus nucleotide (indicated by the 0 values in each column). Mutational effects were scaled so that mean mutational effect is equal to $-1$, as a consequence the additive trait can be interpreted in terms of the number of mismatches from the consensus sequence.

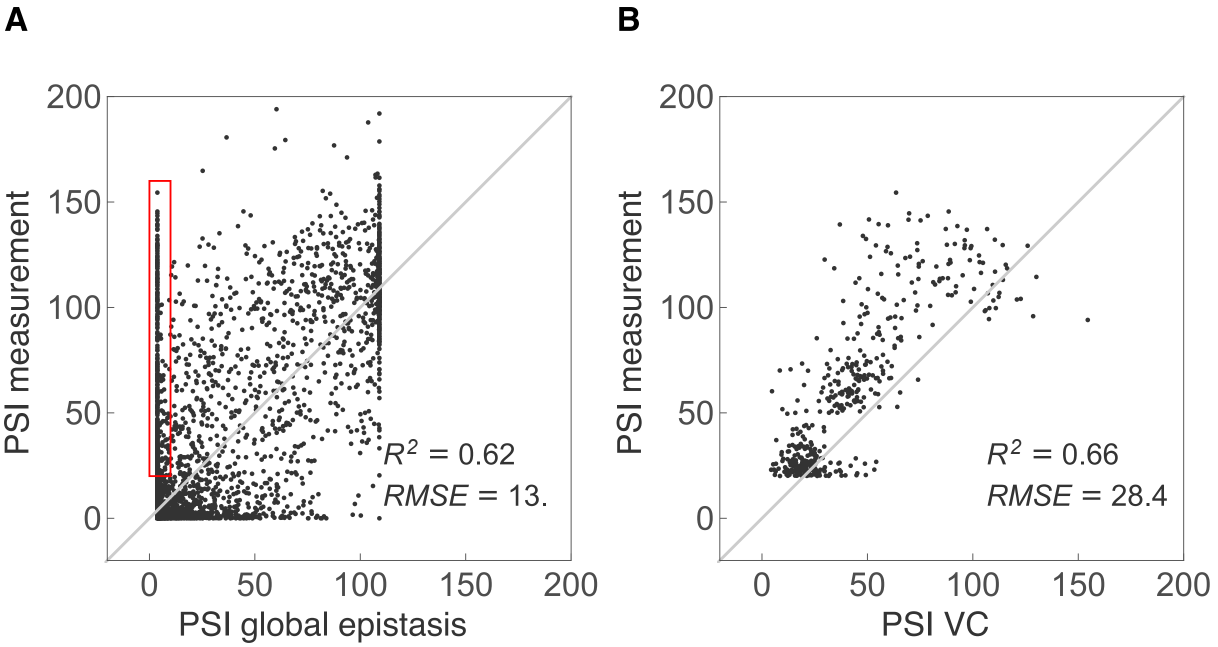


Supplementary Figure 4: The global epistasis model cannot explain the splicing activity of certain 5′ss. (A) Scatter plot showing measured PSI vs. PSI fitted using the global epistasis model. A group of outlier 5′ss with measured PSI $>$ 20 and fittedd PSI $<$ 10 are highlighted by the red rectangle. (B) Scatter plot showing leave-one-out predictions for the boxed outliers using empirical variance component regression.

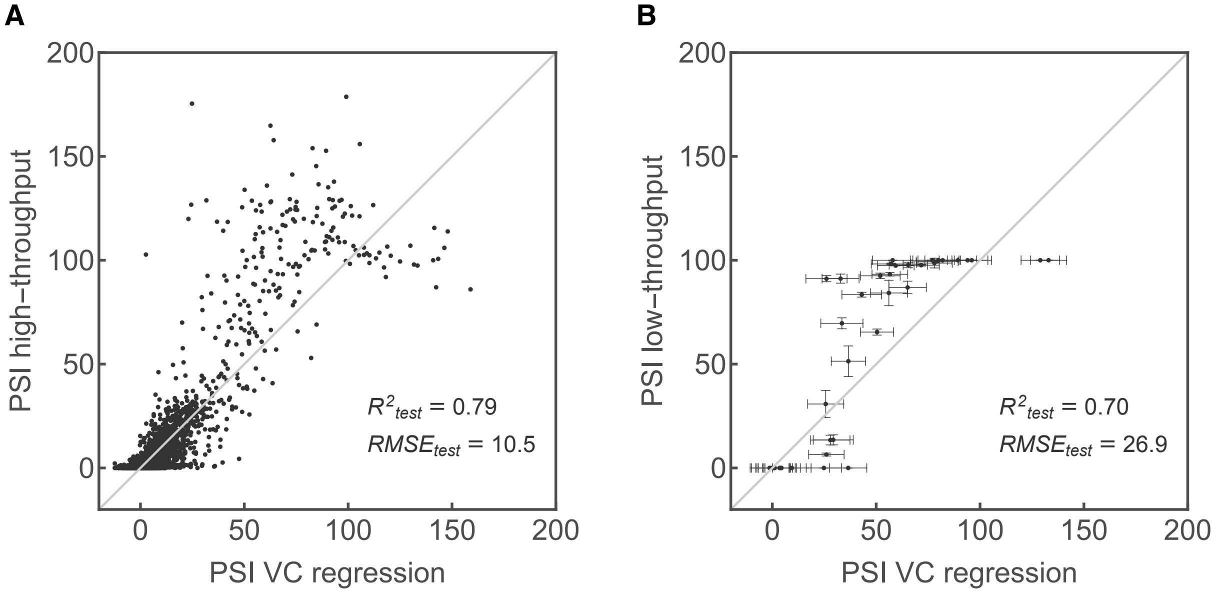


Supplementary Figure 5: Out-of-sample predictions for the SMN1 dataset. (A) Out-of-sample predictions using empirical variance component regression on an 80%-20% training-test split for the high-throughput SMN1 dataset. (B) Out-of-sample predictions using empirical variance component regression based on the entire high-throughput SMN1 dataset for 40 unmeasured SMN1 5′ss not included in the high-throughput dataset. Horizontal error bars correspond to one standard deviation of the posterior distribution. Vertical error bars correspond to one standard deviation around the mean manual validation PSI estimated using three replicates. Data is identical to Figure 5B, but includes the raw empirical variance component predictions rather than restricting these predictions to lie between 0 and 100 PSI.

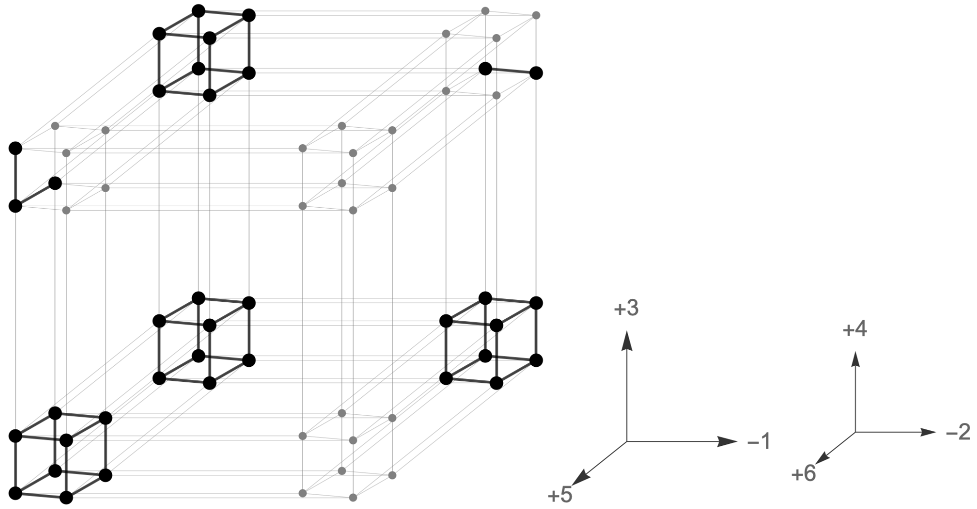


Supplementary Figure 6: Schematic drawing of the $2^6 = 64$ possible combinations of binary states (consensus vs. mutated) on six sites of the 5′ss. Mutations on the three major sites (-1, +3, +5) are represented by the long edges, while mutations on the 3 minor sites (-2, +4, +6) are represented by short edges. Each dot corresponds to the group of all sequences with mutations on a particular subset of the six sites. Groups with 5′ss having PSI > 80 are shown in black with other groups shown in gray. Mutations at -2, +4, +6 that maintain splicing function are shown in black.

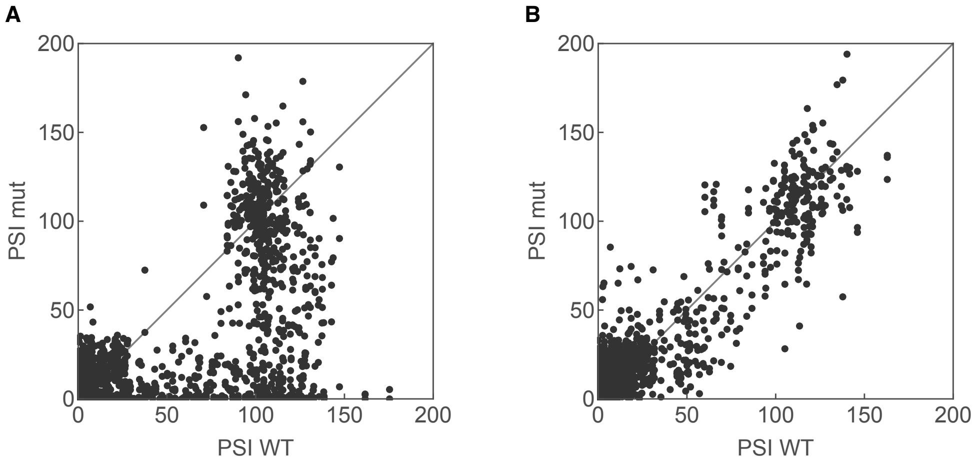


Supplementary Figure 7: Deleterious effects of mutations at position +6 are alleviated by mutations at position +5. (A) Scatter plot showing the PSI of sequences with mutations on +6 vs. the PSI of their counterparts with the consensus U on +6, when site +5 has the consensus G (median mutational effect = −43.0 PSI, calculated in backgrounds with PSI > 80). (B) same as (A), but when +5 contains mutant nucleotides (median mutational effect = −2.0 PSI).

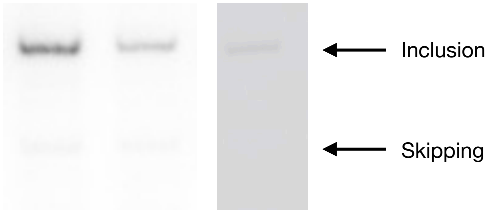


Supplementary Figure 8: Gel image showing three replicates of low-throughput validation of the 5′ss AAG/GUGGAC. Mean PSI $\pm$ 1SD $= 96.9 \pm 5.33$. See *Materials and Methods* for experimental procedures.

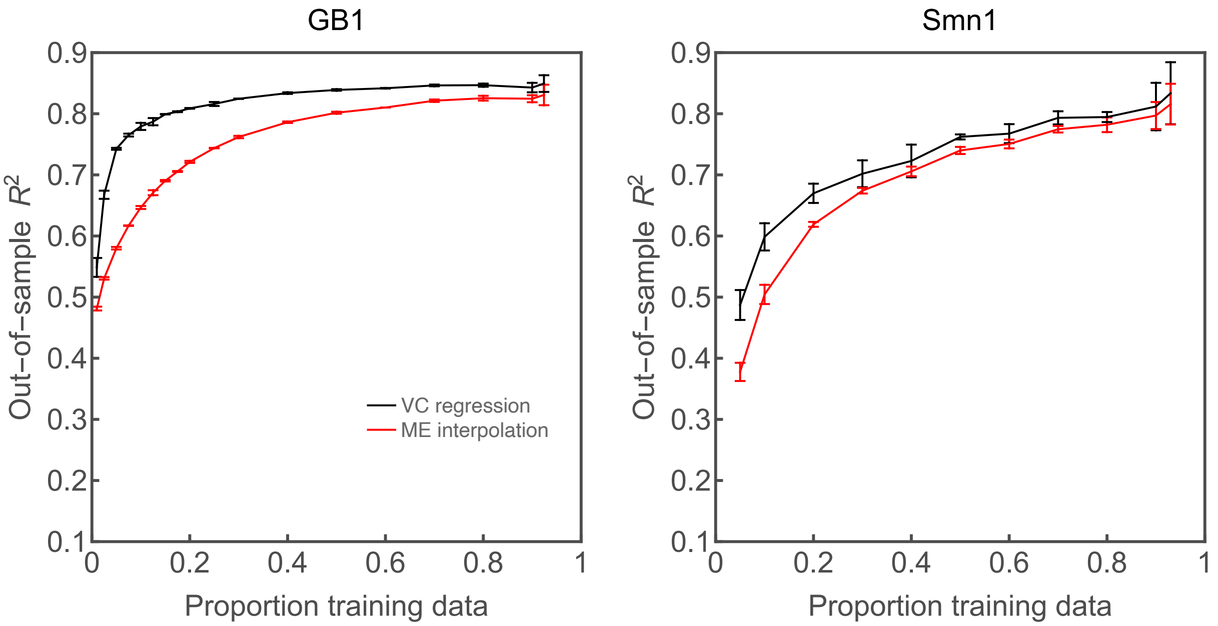


Supplementary Figure 9: Comparison of model performance between empirical variance component regression (VC regression) and minimum epistasis interpolation (ME interpolation) in terms of out-of-sample $R^2$ for a range of training sample sizes calculated for 5 replicates, using the GB1 (left) and SMN1 dataset (right). Error bars represent one standard deviation.

#### References


[1] Jakub Otwinowski and Ilya Nemenman. "Genotype to phenotype mapping and the fitness landscape of the E. coli lac promoter". In: PLoS One 8.5 (2013), e61570.

[2] Rhys M Adams et al. "Epistasis in a fitness landscape defined by antibody-antigen binding free energy". In: Cell systems 8.1 (2019), pp. 86–93.

[3] Trevor Hinkley et al. "A systems analysis of mutational effects in HIV-1 protease and reverse transcriptase". In: Nature Genetics 43.5 (2011), pp. 487–489.

[4] Vladimir I Levenshtein. "Krawtchouk polynomials and universal bounds for codes and designs in Hamming spaces". In: IEEE Transactions on Information Theory 41.5 (1995), pp. 1303–1321.

[5] Robert Happel and Peter F Stadler. "Canonical approximation of fitness landscapes". In: Complexity 2.1 (1996), pp. 53–58.

[6] Peter F Stadler. "Fitness landscapes". In: Biological Evolution and Statistical Physics. Springer, 2002, pp. 183–204.

[7] Peter F Stadler, Robert Happel, et al. "Canonical approximation of landscapes". In: Santa Fe Institute Preprint (1994), pp. 94–09.

[8] Johannes Neidhart, Ivan G Szendro, and Joachim Krug. "Exact results for amplitude spectra of fitness landscapes". In: J. Theor. Biol. 332 (2013), pp. 218–227.

[9] Peter F Stadler and Robert Happel. "Random field models for fitness landscapes". In: J. Math. Biol. 38.5 (1999), pp. 435–478.

[10] Luca Ferretti et al. "Measuring epistasis in fitness landscapes: The correlation of fitness effects of mutations". In: J. Theor. Biol. 396 (2016), pp. 132–143.

[11] Peter F Stadler. "Landscapes and their correlation functions". In: J. Math. Chem. 20.1 (1996), pp. 1–45.

[12] Carl Edward Rasmussen and Christopher K I Williams. Gaussian processes for machine learning. Vol. 1. MIT press Cambridge, 2006.

[13] Richard Hammack, Wilfried Imrich, and Sandi Klavžar. Handbook of product graphs. CRC press, 2011.

[14] Juannan Zhou and David M McCandlish. "Minimum epistasis interpolation for sequence-function relationships". In: Nature communications 11.1 (2020), pp. 1–14.

[15] Luca Ferretti et al. "Measuring epistasis in fitness landscapes: The correlation of fitness effects of mutations". In: Journal of Theoretical Biology 396 (2016), pp. 132–143. ISSN: 10958541. DOI: 10.1016/j.jtbi.2016.01.037.

[16] Claudia Bank et al. "On the (un)predictability of a large intragenic fitness landscape". In: Proc. Natl. Acad. Sci. U.S.A. 113.49 (2016), pp. 14085–14090.

[17] Alexander J Smola and Risi Kondor. "Kernels and regularization on graphs". In: COLT. Vol. 2777. Springer. 2003, pp. 144–158.

[18] Wei-Chia Chen et al. "Field-theoretic density estimation for biological sequence space with applications to 5 splice site diversity and aneuploidy in cancer". In: Proceedings of the National Academy of Sciences 118.40 (2021).

[19] Tinghua Wang, Dongyan Zhao, and Shengfeng Tian. "An overview of kernel alignment and its applications". In: Artificial Intelligence Review 43.2 (2015), pp. 179–192.

[20] JK Haseman and RC Elston. "The investigation of linkage between a quantitative trait and a marker locus". In: Behavior Genetics 2.1 (1972), pp. 3–19.

[21] Michael Betancourt. "A conceptual introduction to Hamiltonian Monte Carlo". In: arXiv preprint arXiv: 1701.02434 (2017).

[22] Radford M Neal. "MCMC using Hamiltonian dynamics". In: Handbook of Markov chain Monte Carlo. Ed. by Steve Brooks et al. 2011, pp. 113–162.

[23] Matthew D Hoffman and Andrew Gelman. "The No-U-Turn sampler: adaptively setting path lengths in Hamiltonian Monte Carlo." In: J. Mach. Learn. Res. 15.1 (2014), pp. 1593–1623.

[24] Hui Zou and Trevor Hastie. "Regularization and variable selection via the elastic net". In: Journal of the Royal Statistic 67.2 (2005), pp. 301–320.

[25] Jerome Friedman, Trevor Hastie, and Rob Tibshirani. "Regularization paths for generalized linear models via coordinate descent". In: Journal of statistical software 33.1 (2010), p. 1.

[26] Ilia Krasikov and Simon Litsyn. "On integral zeros of Krawtchouk polynomials". In: journal of combinatorial theory, Se 74.1 (1996), pp. 71–99.

[27] Mandy S Wong, Justin B Kinney, and Adrian R Krainer. "Quantitative Activity Profile and Context Dependence of All Human 5' Splice Sites". In: Mol. Cell (2018).

[28] TB Parkin and JA Robinson. "Statistical evaluation of median estimators for lognormally distributed variables". In: Soil Science Society of America Journal 57.2 (1993), pp. 317–323.

[29] Yoh Iwasa. "Free fitness that always increases in evolution". In: J. Theor. Biol. 135.3 (1988), pp. 265–281.

[30] Guy Sella and Aaron E Hirsh. "The application of statistical physics to evolutionary biology". In: Proc. Natl. Acad. Sci. U.S.A. 102.27 (2005), pp. 9541–9546.

[31] David M McCandlish, Premal Shah, and Joshua B Plotkin. "Epistasis and the dynamics of reversion in molecular evolution". In: Genetics 203.3 (2016), pp. 1335–1351.

[32] David M McCandlish. "Visualizing fitness landscapes". In: Evolution 65.6 (2011), pp. 1544–1558.
